# CryoSHIELDs: tunable amphipathic peptide additives for cryo-EM

**DOI:** 10.64898/2026.10.01.755993

**Authors:** Caleigh M. Azumaya, Przemysław Dutka, Matthew C. Johnson, Sandeep K. Ravala, Cheng-Guo Wu, Bryce D. Delgado, Francesca Anson, Nairie Michaelian, Christopher W. Koo, Jeffrey Y. K. Tom, Tik Hang Soong, Alister Burt, Shu Ti, Anna C. Howes, Orlando E. Martinez, Aimin Song, Alexis Rohou, Claudio Ciferri, Stanislau Yatskevich

**Affiliations:** Protein Sciences, Genentech Inc., 1 DNA Way, South San Francisco, CA, 94080, USA; Early Discovery Biochemistry, Genentech Inc., 1 DNA Way, South San Francisco, CA, 94080, USA; Thermo Fisher Scientific, 700 Gateway Blvd, South San Francisco, CA, 94080, USA

**Author notes:** These authors contributed equally (Caleigh M. Azumaya, Przemysław Dutka, Matthew C. Johnson). Correspondence: Claudio Ciferri, Stanislau Yatskevich.

## Abstract

While cryogenic electron microscopy (cryo-EM) has transformed structural biology, sample preparation remains a major determinant of success. Specifically, the air-water interface (AWI) can damage complexes, impose preferred orientation, and reduce particle density. Here, we introduce cryoSHIELDs, a library of peptide additives designed to modulate the AWI and improve cryo-EM grid preparation. Across 24 diverse protein systems, cryoSHIELDs improved particle distribution, angular sampling, complex integrity, and, in selected cases, map quality and resolution. CryoSHIELDs enabled structure determination of the OSMRβ–Vixarelimab complex, the multi-subunit CCAN kinetochore complex without chemical cross-linking, and membrane proteins including ABCC6 and the μ-opioid receptor–Gi complex, the latter resolved at 2.3 Å using tenfold lower protein concentration without compromising map quality. CryoSHIELDs provide a simple, tunable, and broadly applicable platform for more reliable cryo-EM in discovery biology and structure-based drug development.

## INTRODUCTION

Cryogenic electron microscopy (cryo-EM) has transformed the structural characterization of biological macromolecules. However, the production of high-quality vitrified specimens remains a primary bottleneck^1–5^. For optimal cryo-EM imaging, particles should be suspended in a thin layer of vitreous ice in a random distribution of orientations. The conventional method for preparing samples requires blotting away excess liquid on a grid to create a free-standing aqueous film a few tens of nanometers thick, which is then vitrified by plunge-freezing in liquid ethane^6^. This procedure generates a system with an exceptionally high surface-to-volume ratio, where low adhesion between water and air molecules allows the cohesive forces of the liquid to form a distinct boundary known as the air-water interface (AWI). Interfacial specimen adsorption frequently causes macromolecular structural denaturation, particle loss, and severe preferred orientation, all of which can preclude high-resolution, isotropic 3D reconstructions^7–10^.

The AWI presents one of the most pervasive problems in cryo-EM sample preparation. Common mitigation strategies include the use of detergents and surfactants^11–14^, modifications of the specimen preparation^15^ or protein constructs^16^ that aim to disrupt protein-AWI interactions, chemical crosslinking^17,18^ to prevent macromolecular unfolding and dissociation of complexes, and the use of custom-made support layers^19–25^ (e.g., graphene or carbon) to sequester proteins away from the AWI. Alternative deposition methods, such as spraying or scribing techniques^26–30^, have also been developed to minimize protein interaction with the AWI and to enable time-resolved studies. While these solutions have been successful for specific targets, they often introduce additional complexity and may require elaborate instrumentation and target-specific optimization. For instance, detergents can destabilize fragile complexes^31^, while crosslinking may lock proteins into non-native conformations, underscoring the need for a more robust, universally applicable preparation tool.

One strategy to mitigate AWI-induced problems is to introduce surface-reactive additives that modify the interactions between macromolecules and the interface, potentially creating a more protein-compatible environment. Designer amphipathic peptides represent a highly tunable class of such additives that, to date, have remained underexplored. In line with this concept, late embryogenesis abundant (LEA) proteins^32^ and a small peptide based on the N-terminus of cyclin B1^33,34^ (BH cryo-tide) have shown promise in a few specific cases. Here, we developed and tested a suite of amphipathic peptides, termed cryoSHIELDs, for mitigating diverse cryo-EM sample preparation pathologies. We found that while a single peptide does not address all cryo-EM sample preparation issues, specific peptides are exceptionally good at addressing particular sample pathologies, such as preferred orientation, particle distribution, or preservation of protein complexes.

## RESULTS

### Sparse matrix screen of cryoSHIELD peptides

We explored two groups of peptides to use as additives in cryo-EM: (i) linear peptides with alternating polar and nonpolar residues along the chain, and (ii) helical peptides where hydrophobic residues line up along one side of an α-helix while polar residues line the opposite side^35,36^ (Table 1 and S1). For the hydrophobic face of linear peptides, we compared aliphatic residues Val and Leu, and aromatic Phe and Trp residues. On the hydrophilic face, we explored Lys, Arg, and Glu residues. Neutral charge peptides are prone to insolubility, self-association, and aggregation, limiting their utility for cryo-EM applications^37^. For helical peptides, we chose sequences based on naturally occurring proteins: part of apo lipoprotein A1 from *Mesocricetus auratus* (cryoH), cecropin B peptide from *Bombyx mori* (cryoM), and a synthetic amphipathic CM15 peptide (cryoB), as they presented diverse surfaces with a range of helical propensity^38^. Consistent with our designs and predictions, amphipathic peptides, but not charged peptides, exhibited surfactant properties in the Marangoni assay^39^, with critical sealing concentration (CSC) values aligning closely with CHAPSO, a commonly used detergent (Table S2). Based on these preliminary values, we chose to use peptides at ∼5-10x of their CSC, 200 μM final concentration, in samples for initial screening.

**Table 1.**
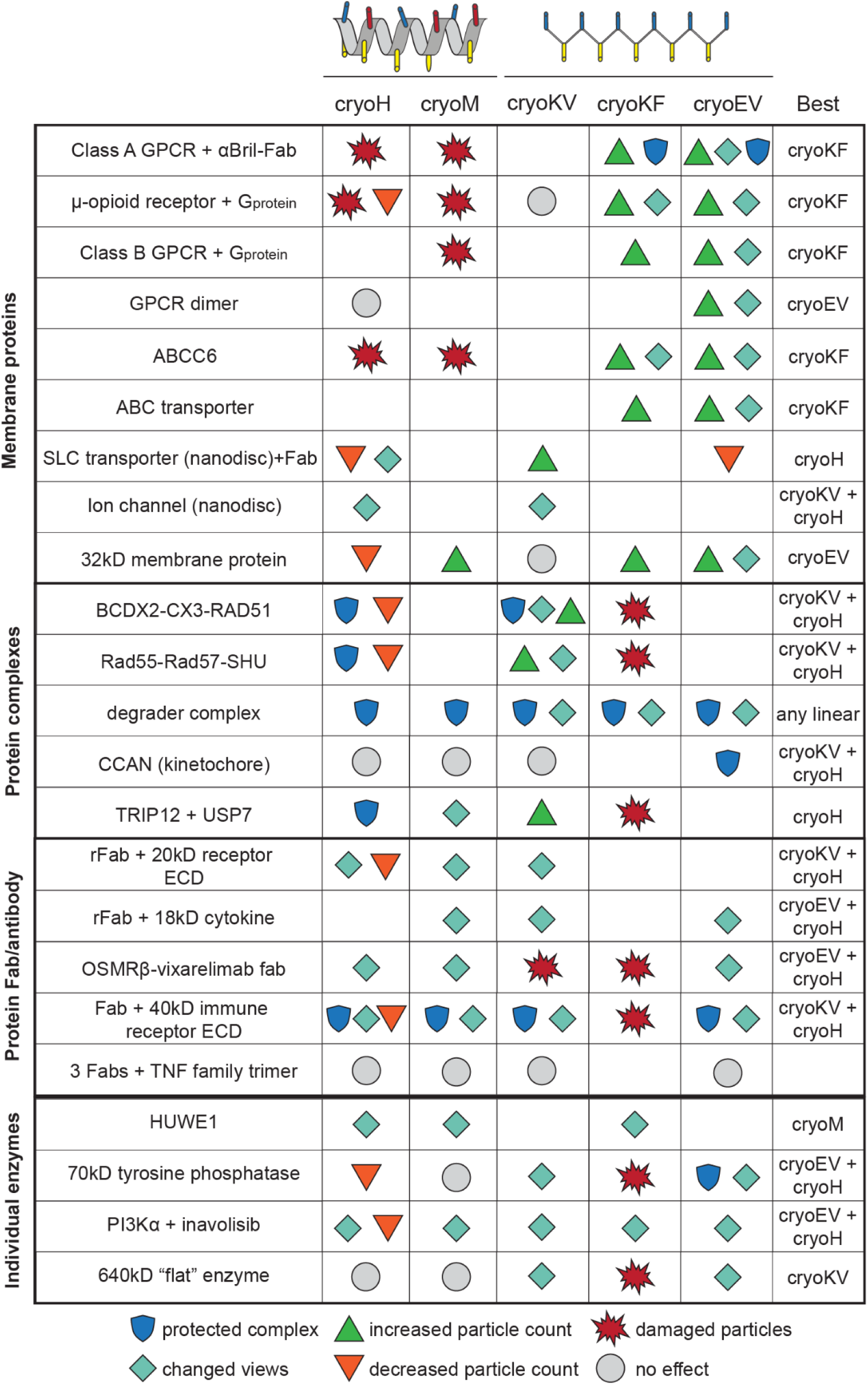
Amphipathic peptides have diverse effects on cryo-EM protein samples. Summary of cryoSHIELD effects observed during routine screening at the Genentech cryo-EM facility. Proprietary proteins have been anonymized, and “rFab” denotes a rigidified Fab^40^. The best cryoSHIELD or combination of cryoSHIELDs is noted in the final column, and for one case, constitutive centromeric associated network (CCAN), is highlighted in green because only a combination of cryoKV + cryoH was effective at preserving the complex integrity. The observed effects of cryoSHIELD addition are defined as follows. **Protect complex**: prevents dissociation of complex components observed in the absence of the additive. **Change views**: altered the particle orientation distribution compared to a sample with no cryoSHIELD. **Increase/Decrease particle count**: qualitatively increased or decreased the particle density observed in micrographs at a constant protein concentration. **Damaged particles**: induced particle damage, most often via large-scale aggregation. **No effect**: yielded no observable difference in complex stability, particle count, or orientation distribution compared to a no additive sample.

|  |  | cryoH | cryoM | cryoKV | cryoKF | cryoEV | Best |
|---|---|---|---|---|---|---|---|
| Membrane proteins | Class A GPCR + αBril-Fab | damaged particles | damaged particles |  | increased particle count, protected complex | increased particle count, changed views, protected complex | cryoKF |
|  | μ-opioid receptor + G<sub>protein</sub> | damaged particles, decreased particle count | damaged particles | no effect | increased particle count, changed views | increased particle count, changed views | cryoKF |
|  | Class B GPCR + G<sub>protein</sub> |  | damaged particles |  | increased particle count | increased particle count, changed views | cryoKF |
|  | GPCR dimer | no effect |  |  |  | increased particle count, changed views | cryoEV |
|  | ABCC6 | damaged particles | damaged particles |  | increased particle count, changed views | increased particle count, changed views | cryoKF |
|  | ABC transporter |  |  |  | increased particle count | increased particle count, changed views | cryoKF |
|  | SLC transporter (nanodisc)+Fab | decreased particle count, changed views |  | increased particle count |  | decreased particle count | cryoH |
|  | Ion channel (nanodisc) | changed views |  | changed views |  |  | cryoKV + cryoH |
|  | 32kD membrane protein | decreased particle count | increased particle count | no effect | increased particle count | increased particle count, changed views | cryoEV |
| Protein complexes | BCDX2-CX3-RAD51 | protected complex, decreased particle count |  | protected complex, changed views, increased particle count | damaged particles |  | cryoKV + cryoH |
|  | Rad55-Rad57-SHU | protected complex, decreased particle count |  | increased particle count, changed views | damaged particles |  | cryoKV + cryoH |
|  | degrader complex | protected complex | protected complex | protected complex, changed views | protected complex, changed views | protected complex, changed views | any linear |
|  | CCAN (kinetochore) | no effect | no effect | no effect |  | protected complex | cryoKV + cryoH |
|  | TRIP12 + USP7 | protected complex | changed views | increased particle count | damaged particles |  | cryoH |
| Protein Fab/antibody | rFab + 20kD receptor ECD | changed views, decreased particle count | changed views | changed views |  |  | cryoKV + cryoH |
|  | rFab + 18kD cytokine |  | changed views | changed views |  | changed views | cryoEV + cryoH |
|  | OSMRβ-vixarelimab fab | changed views | changed views | damaged particles | damaged particles | changed views | cryoEV + cryoH |
|  | Fab + 40kD immune receptor ECD | protected complex, changed views, decreased particle count | protected complex, changed views | protected complex, changed views | damaged particles | protected complex, changed views | cryoKV + cryoH |
|  | 3 Fabs + TNF family trimer | no effect | no effect | no effect |  | no effect |  |
| Individual enzymes | HUWE1 | changed views | changed views |  | changed views |  | cryoM |
|  | 70kD tyrosine phosphatase | decreased particle count | no effect | changed views | damaged particles | protected complex, changed views | cryoEV + cryoH |
|  | PI3Kα + inavolisib | changed views, decreased particle count | changed views | changed views | changed views | changed views | cryoEV + cryoH |
|  | 640kD “flat” enzyme | no effect | no effect | changed views | damaged particles | changed views | cryoKV |

protected complex
increased particle count
damaged particles
changed views
decreased particle count
no effect

As a multi-user, industrial cryo-EM facility, we incorporated cryoSHIELDs into our regular grid preparation and screening workflows against a range of structural targets. Assessed targets included soluble and intramembrane proteins with different stoichiometries and symmetries, stable and unstable complexes, and a broad range of molecular weights. In the absence of cryoSHIELDs, many of these targets exhibited typical cryo-EM grid preparation issues like particle aggregation, complex dissociation, preferred orientation, lack of particles in the ice meniscus, or inconsistent ice thickness. The initial sparse matrix approach allowed us to assess cryoSHIELD effects across different types of cryo-EM targets, and to observe some common trends emerging across twenty-four separate cases (Table 1 and Figure S1).

We observed that different cryoSHIELDs resulted in varied effects across targets. Consistent with the AWI being driven by hydrophobic interactions, charged peptides showed little observable effect on cryo-EM samples (Figure S1A and Table S1). Linear peptides containing Arg at the hydrophilic surface or Trp at the hydrophobic surface frequently, but not always, resulted in protein aggregation or denaturation on the grids, limiting their wide applicability (Figure S1B-S1D). Overall, one of the most potent and reproducible effects of the cryoSHIELDs was enabling difficult cryo-EM targets by altering particle orientations and distribution, which we observed for linear peptides containing Lys or Glu at the hydrophilic surface and either Val or Phe at the hydrophobic surface: cryoKV, cryoEV, and cryoKF (Table 1). Notably, cryoKF aggregated a subset of soluble proteins (Table 1, Figure S1E), but was by far the most advantageous peptide for membrane proteins in detergent. Similarly, we observed that helical cryoH and cryoM peptides were consistently beneficial for cryo-EM samples by improving complex stability. While the helical peptides occasionally improved particle orientation distribution, they were often less efficacious than the linear peptides in this capacity (Table 1). Lastly, we discovered that for soluble proteins a 1:1 mixture of linear and helical peptides resulted in synergistic effects. The combination was able to alleviate multiple sample preparation issues simultaneously, such as orientation and complex stability (Table 1, “Best” column).

### CryoSHIELDs improve angular sampling in orientation-limited specimens

Preferred orientation limits the structural analysis of many biologically and therapeutically relevant assemblies, including Fab–antigen complexes, degrader assemblies, and particles with broad, flat surfaces. Across these sample classes, cryoSHIELDs altered particle orientations, although the extent of improvement depended on both the peptide and the target (Figure 1A-1C, S2, and Table 1). These additives effectively mitigated orientation bias in multiple Fab-antigen complexes, and different cryoSHIELDs yielded distinct view distributions (Figure 1A and S2A-S2B). In most instances, both linear and helical peptides broadened the orientational landscape of the complexes, as evidenced by increased conical FSC area ratio (cFAR) values (Figure 1D and S2I), with helical peptides generally being less efficacious in this capacity. Notable exceptions included cryoKF, which frequently reduced usable particle counts due to aggregation (Figure 1A, S1E, and S2C), and cryoM, which sometimes introduced unique but strong preferred orientations when used alone (Figure 1A and 1D). Among individual cryoSHIELDs, cryoKV, and cryoEV were potent in inducing new and wide distribution of views for most Fab-antigen complexes.

**Figure 1.**
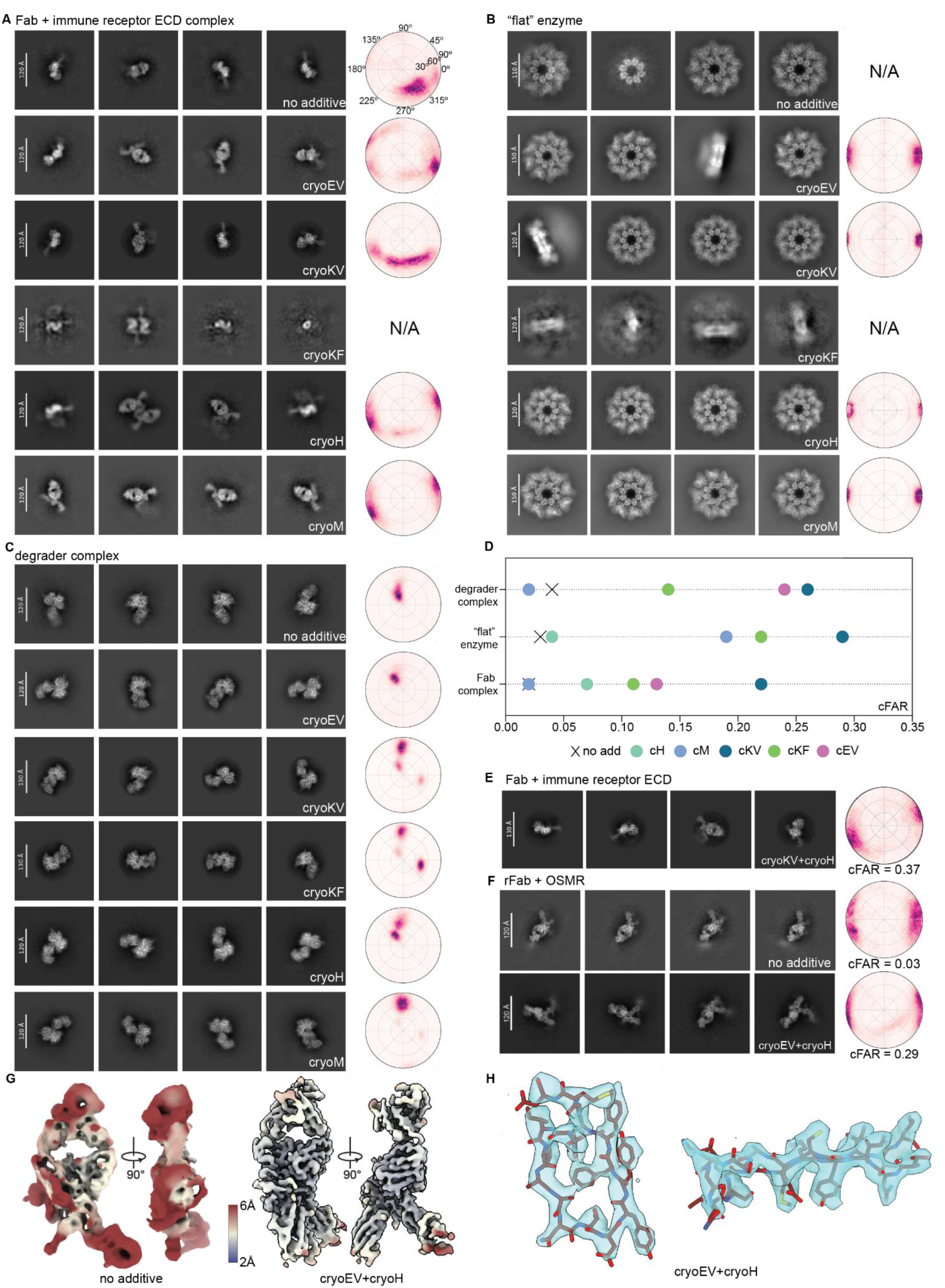
CryoSHIELDs alter the orientational landscape of diverse biological samples. (A) Representative 2D class averages and view direction distributions from 3D reconstructions of a Fab-antigen complex prepared with different cryoSHIELDs. CryoKF aggregated this sample, resulting in low-resolution 2D class averages that precluded structure determination, noted with N/A. Scale bars in 2D averages are indicated in each panel. ECD - extracellular domain. (B) Representative 2D class averages and view direction distributions from 3D reconstructions of an octameric enzyme prepared with different cryoSHIELDs. Datasets that did not yield a sufficient number of particles or views to generate a 3D map are noted with N/A. Scale bars in 2D averages are indicated in each panel. (C) Representative 2D class averages and view direction distributions from 3D reconstructions of a E3 ligase degrader complex prepared with different cryoSHIELDs. Scale bars in 2D averages are indicated in each panel. (D) Conical FSC area ratio (cFAR) values and particle per micrograph numbers for each of the datasets presented in A-C. Values for maps generated without additives are denoted by a black X. For ‘flat’ enzyme, cFAR value was determined from a tilted no-additive dataset. Particles per micrograph were calculated from final 3D reconstructions or “particle-like” 2D classes if no 3D reconstruction was possible. (E) Representative 2D class averages of the Fab-antigen complex from (A) but prepared with cryoKV and cryoH peptide combination. (F) Representative 2D class averages of oncostatin M receptor β subunit (OSMRβ)–Fab complex prepared without additives or with cryoEV-cryoH. (G) Comparison of OSMRβ–Fab reconstructions obtained without additives (left) or with cryoEV–cryoH (right) colored by resolution. (H) Representative atomic model-to-map fits of the OSMRβ–Fab complex illustrate the quality of the resulting density and the resolution of structural details.

We found that combining linear and helical cryoSHIELDs was often equally or more effective than using individual peptides. For the same Fab–immune receptor complex, a 1:1 mixture of cryoKV and cryoH increased cFAR to 0.37, exceeding either peptide alone (Figure 1D and 1E). To further investigate whether improved angular sampling translates into higher-quality reconstructions, we examined the oncostatin M receptor β subunit (OSMRβ) bound to a Fab derived from Vixarelimab. Without additives, the complex exhibited severe preferred orientation and yielded a poorly resolved reconstruction (Figure 1F, 1G and S2D). Combining cryoEV and cryoH enabled a 3.37 Å reconstruction from a small screening dataset, allowing confident fitting of the atomic model into the cryo-EM density (Figure 1H and S2E). Similar benefits were observed for two additional Fab complexes, comprising an 18 kDa cytokine and a 20 kDa receptor extracellular domain (Figure S2A and S2B). Although individual linear peptides, such as cryoEV and cryoKV, induced additional particle orientations, their combination with helical peptides, cryoH or cryoM, further diversified the observed views. Together, these results suggest that combining linear and helical cryoSHIELDs at a final concentration of 200 µM provides an effective starting strategy for mitigating preferred orientation in Fab-containing complexes.

We further assessed the concentration dependence of peptide effects directly using a Fab–antigen complex (Figure S2F–H). As observed for many other Fab-containing samples, the combination of cryoKV and cryoH improved the distribution of particle orientations. Titration of the peptide mixture revealed concentration-dependent changes in particle orientation, with 200 µM producing the most favorable angular distribution and the highest cFAR value for this sample. This is also consistent with CSC values determined using Marangoni assay. We observed notable improvement in cFAR values for peptide concentration starting at 50 µM for this sample (Figure S2G).

The benefits extended to a 640 kDa soluble octameric enzyme with pronounced preferred orientation (Figure 1B). Structure determination had previously only been possible after collecting large, tilted datasets, resulting in prolonged experiments and resolutions that limited the utility of structures in iterative drug design. Testing individual peptides again indicated that cryoKV, cryoEV, cryoM, and cryoH were all able to increase the cFAR of 3D reconstructions in this system (Figure 1D). While cryoM, cryoH, and cryoEV all allowed 3D reconstruction, only 2, 3, and 9% of the views were well aligning side views with these cryoSHIELDs. However, cryoKV induced 20% of well-aligning particles to be high-resolution side views (Figure 1B). This facilitated easier high-resolution structure determination without stage tilting, resulting in a cFAR score ten times higher than a map determined from a tilted sample without additive while maintaining the number of particles per micrograph similar to the no additive condition (Figure 1D, S2I). As a result, the average resolutions of the maps determined for this system have routinely improved, allowing for the timely and confident placement of small molecules and waters. Similar to many other cases, cryoKF, cryoRV, and cryoRF induced heavy sample aggregation of this target (Figure 1B and S1D).

We next examined an E3 ubiquitin ligase–degrader assembly, another sample class frequently affected by preferred orientation^41^ (Figure 1C). Traditional approaches, such as combining graphene oxide with stage tilt during data collection, enabled a ∼3.5 Å reconstruction, but a large data collection and extensive processing were needed to reach this resolution, preventing routine structure-guided drug design approaches. For this method, only 10 particles per micrograph contributed to the final reconstructions from a representative tilted graphene oxide data collection (Figure 1D). Application of most single cryoSHIELDs resulted in a sufficient particle orientational range to reconstruct maps at ∼2.5 Å from at least three times less data, substantially reducing microscope and computational resource investment. For the degrader samples, the cFAR value for samples with a linear cryoSHIELD increased 3-6 fold compared to without an additive (Figure 1D and S2I). Unlike several other soluble targets, this assembly also tolerated cryoRF, emphasizing the target dependence of peptide effects, while peptide combinations similarly improved the distribution of views (Figures S1B, S1D, and S2J). Rosenthal–Henderson analysis showed that cryoKV, cryoKF, and cryoRF datasets achieved higher nominal resolution at comparable particle numbers than the additive-free dataset (Figure S2K).

Overall, most cryoSHIELDs changed the distribution of views of the target molecule, consistent with their function at the AWI. CryoKV was particularly effective across the systems examined, whereas combinations of linear and helical peptides appear to provide additional benefits. We therefore recommend a focused initial screen of 1:1 mixtures of cryoKV or cryoEV with cryoH or cryoM, at a total peptide concentration of 200 µM. These empirical starting conditions reduce the scope of optimization rather than define a universal optimum; cryoSHIELDs’ utility should be judged by particle integrity, angular sampling, and map interpretability together.

### CryoKF improves GPCR cryo-EM behavior

Membrane proteins comprise a large and biologically important protein class and account for 60% of Food and Drug Administration approved drug targets^42,43^. However, their structural characterization is frequently hindered by low expression levels and poor biochemical stability. We characterized the cryoSHIELD toolkit effects across multiple membrane protein families, including GPCRs, ABC transporters, SLC transporters, and an ion channel (Table 1). Our initial sparse matrix revealed that cryoKF and cryoEV, specifically, exerted robust and reproducible effects on GPCRs.

Standard cryo-EM workflows for GPCR-G protein complexes typically necessitate high protein concentrations^44^, usually 5–10 mg/mL, to achieve sufficient particle density in vitreous ice and to preserve the integrity of labile receptor complexes from damage at the AWI. During sample preparation, lauryl maltose neopentyl glycol (LMNG), a detergent commonly used for GPCR purification, can concentrate together with the protein and promote train- or tube-like assemblies that complicate cryo-EM data analysis^45^ (Figure 2A, left). By contrast, cryoKF dramatically enriched GPCR-G protein complex particles within the grid holes, facilitating the use of significantly less purified material without inducing LMNG-driven artifacts (Figure 2A, middle).

**Figure 2.**
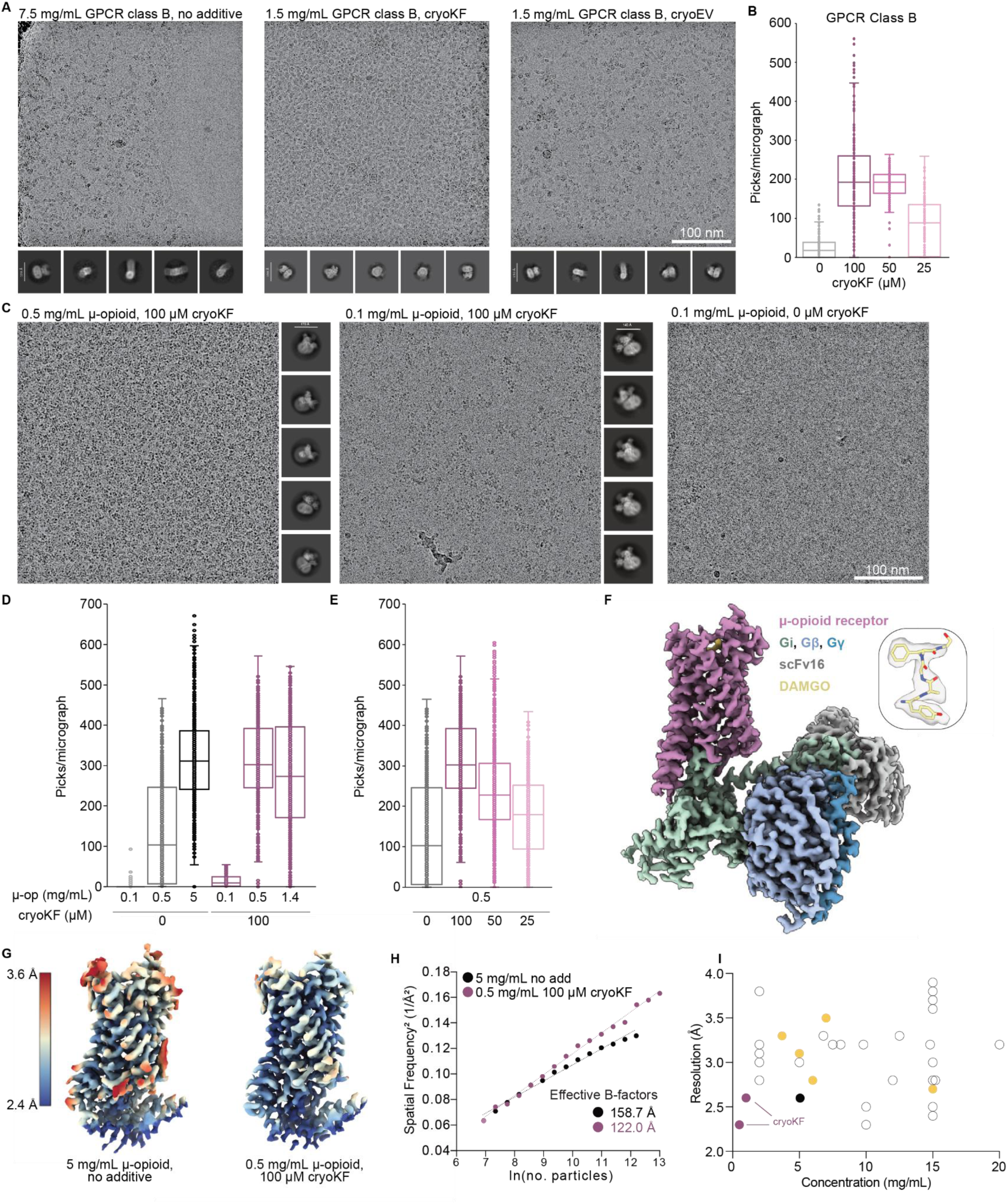
CryoSHIELDs improve the particle distribution of GPCR complexes within vitrified ice. (A) Micrographs and 2D class averages from a Class B GPCR-G protein complex with (left) excess LMNG micelles present at a typical high protein concentration and no additive, (middle) well distributed GPCR-G-protein complexes with 100 µM cryoKF at 1.5 mg/mL, and (right) good particle distribution, but some micelle contamination remaining with 200 µM cryoEV at 1.5 mg/mL. Micrographs scale bar, 100 nm. Scale bars in 2D averages are indicated in each panel. (B) Quantification of nominal particle picks across samples of a Class B GPCR-G protein complex with varying concentrations of cryoKF. 900 micrographs, across 3 grid types, were analyzed for each condition. (C) Representative cryo-EM micrographs and 2D class averages of the μ-opioid receptor–Gi complex prepared at 0.5 or 0.1 mg/mL with 100 µM cryoKF, or at 0.1 mg/mL without additives. The particles are confirmed to be intact complexes in 2D class averages from the cryoKF grids. Micrographs scale bar, 100 nm. Scale bars in 2D averages are indicated in each panel. (D) Quantification of particle picks per micrograph for μ-opioid receptor–Gi samples prepared at the indicated protein concentrations (0.1–5 mg/mL) in the absence or presence of 100 µM cryoKF. (E) CryoKF concentration-dependent enrichment of μ-opioid receptor–Gi particles at a fixed protein concentration of 0.5 mg/mL. (F) Composite cryo-EM reconstruction of the DAMGO-bound μ-opioid receptor–Gi complex determined at 2.3 Å global resolution from a sample containing 0.5 mg/mL protein and 100 µM cryoKF. The μ-opioid receptor, Gαi, Gβ, Gγ, and scFv16 are colored as indicated. Inset shows the DAMGO ligand fitted into the cryo-EM density. (G) Comparison of focused μ-opioid receptor cryo-EM maps obtained at 5 mg/mL without additives (left) and 0.5 mg/mL with 100 µM cryoKF (right), colored by local resolution (2.4–3.6 Å). (H) Rosenthal–Henderson analysis of μ-opioid receptor–Gi datasets obtained at 5 mg/mL without additives or at 0.5 mg/mL with 100 µM cryoKF. Inverse-squared resolution (Å⁻²) is plotted against the natural logarithm of particle number, with linear regression fits shown. Effective B-factors were calculated as twice the reciprocal of the fitted slope, yielding 159 Å² without additives and 122 Å² with cryoKF. (I) Resolution of µ-opioid receptor–G_i_ protein complex structures deposited in the PDB plotted as a function of protein concentration. Maps presented in this manuscript are colored in pink for cryoKF containing conditions or in black for the no additive condition, and maps determined in the presence of DAMGO are colored in yellow. PDB accession codes for all data points are supplied in Table S3.

This also resulted in 2D class averages with high-resolution structural features without the need for any elaborate data processing. We saw similar, but less pronounced, results with cryoEV (Figure 2A, right). This enrichment was dose-dependent: particle density within the holes decreased as the cryoKF concentration was titrated down (Figure 2B and S3A). Concentrations of 0.5–1.5 mg/mL consistently yielded complete grid coverage with high-quality particles.

To directly benchmark cryoKF in a well-characterized model system, we applied it to the μ-opioid receptor in complex with heterotrimeric G protein G_i_ and the agonist peptide DAMGO, enabling comparison to previously reported structures obtained at conventional, high protein concentrations (Table S3). Consistent with our observations for other GPCR targets, cryoKF promoted pronounced particle enrichment in vitreous ice and produced a uniform particle distribution across grid holes at 0.5 mg/mL receptor concentration (Figure 2C, left). Strikingly, even at 0.1 mg/mL, cryoKF-containing samples maintained intact, discrete complexes that generated well-resolved 2D class averages; whereas, samples prepared without cryoKF, contained no detectable particles (Figure 2C, middle and right, and Figure S3B). To quantify the effect of cryoKF on particle enrichment, we compared μ-opioid receptor–Gi samples across different protein and peptide concentrations (Figure 2D and 2E). At 0.5 mg/mL receptor and 100 µM cryoKF, particle density was comparable to that obtained at 5 mg/mL without additives. Furthermore, at a fixed receptor concentration of 0.5 mg/mL, particle enrichment increased with cryoKF concentration from 25 to 100 µM, demonstrating a dose-dependent effect. Overall, cryoKF could reduce the protein concentration required for µ-opioid receptor structure determination by ten-to fifty-fold relative to standard workflows.

Using cryoKF, we determined a 2.3 Å structure of the µ-opioid receptor–G_i_ protein complex from a typical 6,800 micrograph dataset collected at 0.5 mg/mL receptor concentration (Figure 2F and S3B). Increasing the protein concentration to 1 mg/mL resulted in a slightly lower nominal resolution of 2.6 Å, potentially reflecting excessive particle crowding in the presence of cryoKF. To directly evaluate whether the substantial reduction in protein concentration or inclusion of the cryoKF additive compromised structural quality, we collected an additional dataset without additives at 5 mg/mL, a concentration commonly used for GPCR structure determination (Figure S3B). This yielded a global 2.6 Å reconstruction, compared with 2.3 Å for the cryoKF-containing sample at a ten-fold lower concentration. Importantly, the cryo-EM density improvement was particularly apparent in the local resolution of the receptor transmembrane region, where the cryoKF map displayed better-resolved helical density (Figure 2G). Consistently, Rosenthal– Henderson analysis revealed a steeper slope for the cryoKF dataset, corresponding to a lower apparent B-factor and more efficient recovery of high-resolution information with increasing particle numbers (Figure 2H). CryoKF also altered the angular distribution of the receptor complex (Figure S3C), suggesting that improvements in particle distribution and angular sampling likely collectively contribute to reconstruction quality. Thus, cryoKF enabled a ten-fold reduction in protein concentration without compromising, and in some cases while improving, the quality of the resulting structure.

Comparison with previously published μ-opioid receptor–Gi structures further illustrates that our cryoKF-assisted reconstruction achieves high-resolution at substantially lower protein concentrations than typically reported (Figure 2I and Table S3). However, given the differences in experimental conditions across published studies, our conclusions are primarily supported by the direct comparisons presented above. We also evaluated the remaining cryoSHIELDs using the μ-opioid receptor–Gi complex. Consistent with observations for other GPCRs, cryoEV modestly increased particle density and diversified angular sampling, whereas most other peptides had little effect or yielded less well-resolved 2D class averages (Figures S3D and S3E).

To extend these observations, we prepared another class B GPCR-G protein complex in the presence of cryoKF and achieved a 2.95 Å reconstruction at just 0.1 mg/mL receptor concentration (Fig. S4A and S4B). In contrast, grids prepared in parallel at 0.1 mg/mL without cryoKF contained no particles across multiple attempts, and adequate particle coverage for this target without additives could only be achieved at concentrations of 5–10 mg/mL (Figure S4C). Thus, for this complex, cryoKF allows successful structure determination while using as little as fifty-fold lower protein concentration. Lastly, cryoEV or cryoKF in combination with cryoH also enriched a GPCR dimer lacking bound G proteins (Figures S4D and S4E), extending the utility of cryoSHIELD across GPCR classes.

By facilitating a significant reduction in the required protein concentration and improving the distribution of views as well as particle quality, cryoKF addresses many bottlenecks of GPCR cryo-EM structure determination. This advantage should be especially valuable for complexes that can be assembled without orthosteric ligands, allowing a single protein preparation to support screening of multiple ligands and thereby accelerating high-throughput GPCR structure-determination pipelines.

### CryoKF facilitates ABCC6 structure determination

Beyond their application to GPCRs, we found that cryoKF and cryoEV provided benefits across diverse membrane protein classes solubilized in common detergents, such as LMNG or Glyco-Diosgenin (GDN) (Table 1). To evaluate the utility of these peptides for novel, challenging targets, we chose to apply the cryoSHIELD toolkit to the ABCC6 transporter, a membrane protein with no reported experimental structure. Mutations in ABCC6 cause pseudoxanthoma elasticum (PXE) and generalized arterial calcification of infancy (GACI), diseases characterized by ectopic calcification of elastic tissues such as skin, eyes, and the cardiovascular system^46,47^. Similar to many other membrane proteins, ABCC6 suffers from low expression yields. At a concentration of 3 mg/mL, particles were visible within the vitreous ice layer, but very few 2D class averages displayed distinct structural features, and the few well-resolved 2D classes exhibited a preferred orientation, preventing high-resolution 3D reconstructions (Figure 3A, left and S5A).

**Figure 3.**
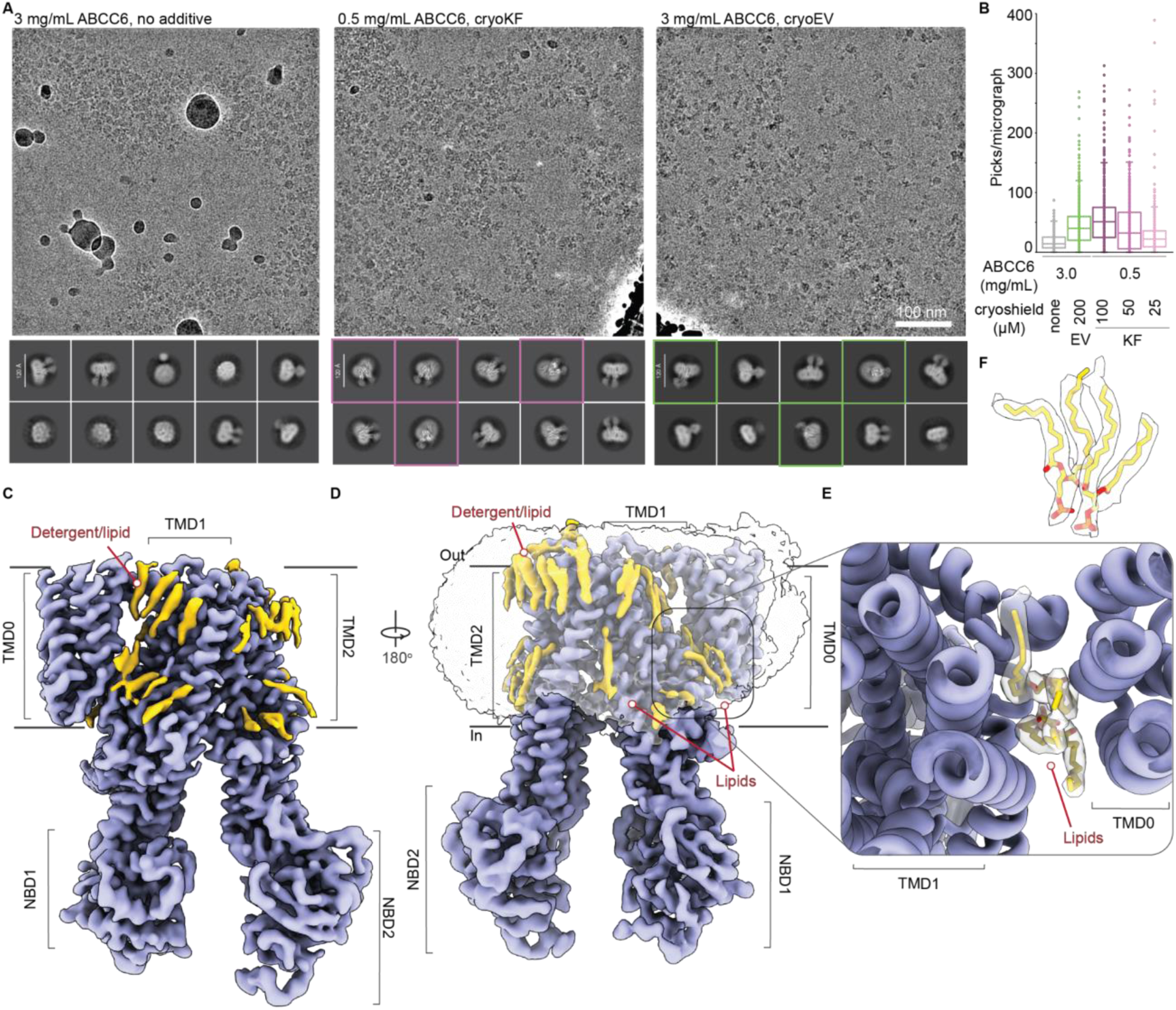
CryoSHIELDs enable structure determination of ABCC6. (A) Representative micrographs and 2D class averages, for ABCC6 vitrified at 3 mg/mL without additives, 0.5 mg/mL with 100 µM cryoKF, and 3 mg/mL with 200 µM cryoEV. Scale bar, 100 nm. Scale bars in 2D averages are indicated in each panel. Unique views not present in the no additive sample are highlighted with colored boxes. (B) ABCC6 particles per micrograph across different conditions. 900 micrographs, across 3 grid types, were analyzed for each condition. (C) Composite cryo-EM map of ABCC6 determined at 2.49 Å resolution with cryoKF additive. The protein density map is colored in purple and additional density attributed to detergent, cholesterol, or lipids is colored in gold (D) Composite cryo-EM map of ABCC6 with the superimposed detergent micelle density in transparent representation. (E) A zoom in on the TMD1-TMD0 interface viewed in cartoon representation, with lipid model colored in gold and built into the transparent cryo-EM density map. (F) Interfacial lipid model built into the cryo-EM density map.

From equivalently sized screening datasets, cryoKF both diversified particle orientations and facilitated uniform particle distribution at a six-fold lower protein concentration (0.5 mg/mL), mirroring our observations with GPCRs (Figure 3A and 3B). CryoEV, analogous to its effect on GPCRs, resulted in more intact ABCC6 particles at 3 mg/mL concentration and additionally a more diverse distribution of views compared to no additive grids (Figure 3A and 3B). A full data collection of the cryoKF condition enabled ABCC6 structure determination at a global 2.49 Å resolution (Figure 3C-D, and S5B).

The cryo-EM structure of ABCC6, captured in an inward-facing conformation, revealed two canonical transmembrane (TM) domains (TMD1 and TMD2), two canonical nucleotide-binding domains (NBD1 and NBD2), as well as an additional N-terminal TMD (TMD0) characteristic of a few specific ABC subfamily C members (Figure 3C and 3D). TMD0 is rigidly anchored against TMD1, an interaction stabilized by two well-ordered lipid molecules sequestered at the domain interface (Figure 3E and 3F). The presence of numerous other lipid and detergent densities surrounding the ABCC6 scaffold indicates that the cryoKF additive does not perturb or displace the local lipid environment (Figure 3C-3F).

While ABCC6 shares high homology with its paralog ABCC1^48^ and some homology with other members ABCC2^49^ and ABCC8^50^, it exhibits a distinct functional orientation for its accessory domain TMD0 (Figure S5C). In ABCC1, TMD0 mediates homodimerization, whereas in ABCC8, it serves as a regulatory tether for the K_ATP_ channel. In contrast, the ABCC6 TMD0 is rotated and tilted inward towards TMD1, likely because of the preserved structural lipids that stabilize this interface.

The determination of the ABCC6 architecture serves as a proof-of-concept, demonstrating that cryoKF can transform previously challenging, low-yield membrane proteins into high-resolution targets for structural biology and drug discovery. In this case, the principal benefit of cryoKF was likely the emergence of well-resolved side and high-tilt views that were largely absent without the additive, substantially improving angular sampling. The preservation of fine-scale features, such as the TMD0-stabilizing lipids, demonstrates that cryoSHIELDs improve sample behavior without compromising structural integrity.

### CryoSHIELD effects on other membrane proteins

To further establish the general applicability of cryoSHIELDs, we tested a second ABC transporter characterized by an exceptionally poor yield (∼20 μg of purified protein from 2 L of Expi293F expression). In the absence of additives, a 3 mg/mL sample yielded almost no usable particles and poorly resolved 2D class averages (Figure S5D and S5E). Conversely, the inclusion of cryoKF ensured robust particle coverage and well-resolved 2D classes at a 0.5 mg/mL protein concentration. In this sample, cryoEV was critical for improved particle distribution and provided unique high-tilt views. Combining datasets from cryoKF and cryoEV samples enabled structure determination for this challenging target.

We also addressed the challenge of membrane proteins reconstituted into lipid nanodiscs, which sometimes exhibit preferred orientations^51^. We applied a suite of cryoSHIELDs to a few nanodisc-embedded protein families. Globally, we observed that nanodisc-reconstituted samples behaved more similarly to soluble than detergent-solubilized proteins. We observed that cryoH specifically induced side views for an SLC transporter with a Fab bound to its extracellular face (Figure S5F). This view enabled structure determination of this previously intractable system. While other cryoSHIELDs, namely cryoKV and the combination of cryoH and cryoEV, increased the particle count per micrograph, they did not induce side views (Figure S5G). Similar enrichment of side views was observed for a nanodisc-reconstituted ion channel when a combination of cryoH and cryoKV were used (Table 1).

In summary, while cryoKF and cryoEV are extremely effective for detergent-solubilized membrane proteins, nanodisc-reconstituted samples behave more similarly to soluble protein targets, where cryoH alone or in combination with a linear peptide tends to benefit membrane protein complexes through reorientation.

### CryoSHIELDs stabilize protein complexes

One of the main reported functions of the LEA proteins is preserving protein complexes on cryo-EM grids^32^. We therefore hypothesized that cryoSHIELDs would similarly protect protein complexes prone to denaturation or dissociation at the AWI (Table 1). One of the cases we examined in detail was that of the *Bombyx mori* 12-subunit holocentric inner kinetochore CCAN (<u>C</u>onstitutive <u>C</u>entromere-<u>A</u>ssociated <u>N</u>etwork) complex^52^. Conventionally, CCAN structural studies require sophisticated “on-column” crosslinking protocols to maintain integrity^52,53^. Samples vitrified in the absence of crosslinkers and cryoSHIELD additives yielded micrographs with apparent particle density, but 2D classification failed to produce coherent or structurally informative averages (Figure 4A). While cryoEV and cryoH yielded some intact 2D class averages, the cryoKV + cryoH mixture preserved the CCAN holocomplex completely without any exogenous crosslinking agents (Figure 4A, 4B and S6A). Other individual peptides or LEA proteins were insufficient to yield any 2D class averages for CCAN (Figure S6A).

**Figure 4.**
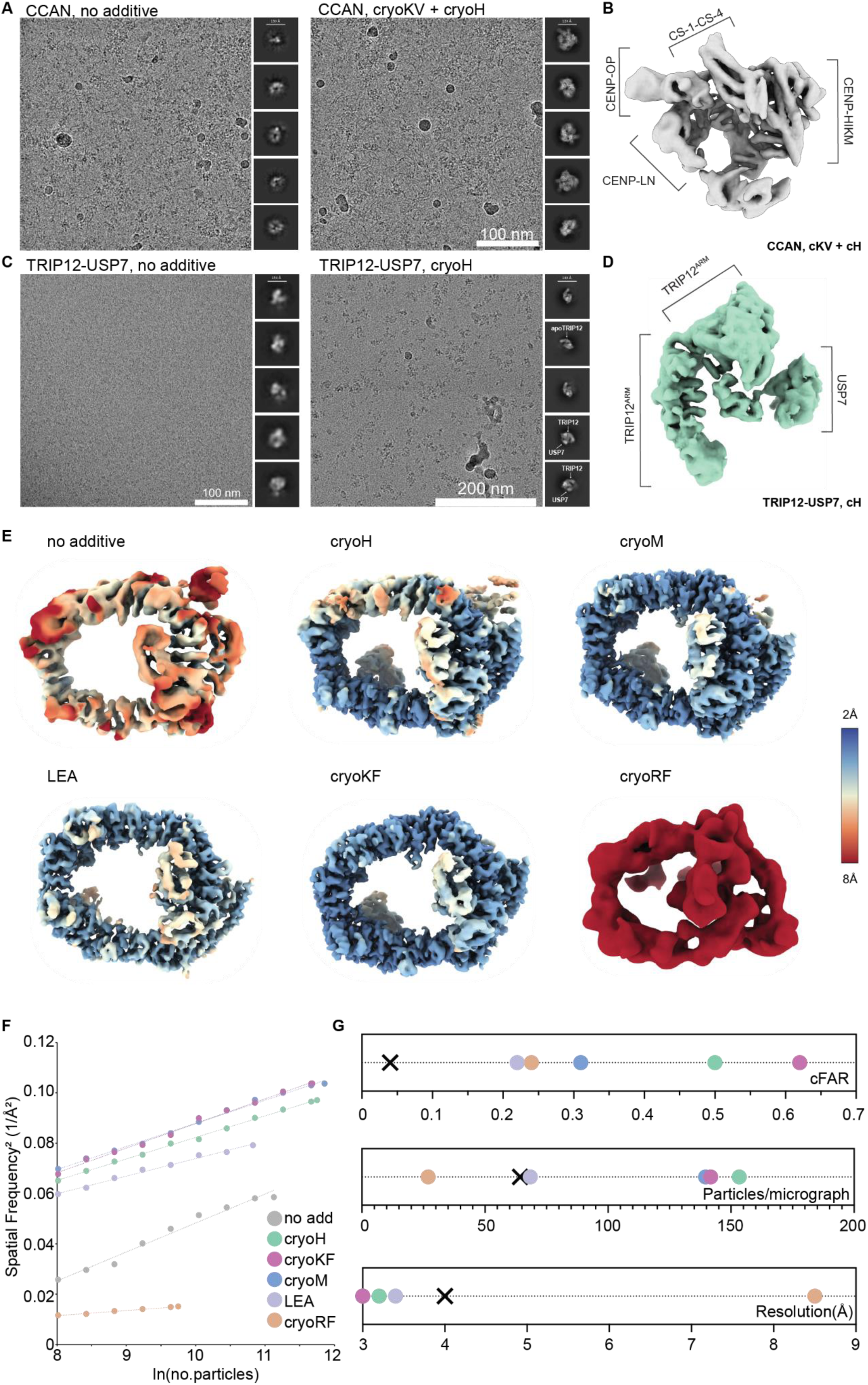
CryoSHIELDs stabilize complexes and improve particle alignment. (A) Representative micrographs and 2D class averages of the 12-subunit CCAN complex in the presence and absence of a 1:1 cryoH-cryoKV mixture. Scale bar, 100 nm. (B) Cryo-EM CCAN reconstruction from a 1:1 cryoH-cryoKV mixture screening dataset. A map could not be generated from the no additive sample. (C) Representative micrographs and 2D averages of the TRIP12-USP7 complex in the presence and absence of cryoH. Scale bars are 100 nm and 200 nm. (D) Cryo-EM TRIP12-USP7 3D reconstruction from the cryoH screening dataset. A map could not be generated from the no additive sample. (E) Representative maps generated from 4000 micrograph datasets for HUWE1 vitrified with different cryoSHIELDs and the LEA protein. All maps are colored by their local resolution with a scale from 2 to 8 Å. (F) Rosenthal–Henderson analysis for the final particles used in reconstructions for each of the datasets shown in panel E. The cryoRF dataset suffered from extensive aggregation and therefore had many fewer particles for reconstruction. (G) Comparison of conical FSC Area Ratio (cFAR) scores, particles per micrograph, and average resolution in each of the HUWE1 datasets in panel E.

We extended this stabilization principle to the TRIP12 E3 ubiquitin ligase, which we discovered forms a previously uncharacterized complex with the USP7 de-ubiquitinase (Figure S6B). Although individual TRIP12 (apo-TRIP12) resulted in well-resolved 2D class averages (Figure S6C), similar to reported TRIP12 structures^54^, the TRIP12-USP7 complex was very sensitive to denaturation during cryo-EM grid preparation (Figure 4C). Traditional chemical crosslinking failed to resolve this issue, yielding only amorphous globular particles unsuitable for high-resolution analysis despite monodisperse distribution on grids (Figure S6D). However, the inclusion of cryoH during grid preparation successfully stabilized the interaction, revealing an additional density (Figure 4C and D). Although a population of apo-TRIP12 remained visible, the complex was sufficiently stabilized for structural characterization in the absence of any chemical crosslinking. While other additives like cryoM provided unique orientations of the TRIP12 scaffold, we were unable to determine if they prevented the dissociation of the USP7 subunit (Figure S6E).

The cryoEV peptide showed an additional benefit when applied to a GPCR-BRIL fusion bound to an anti-BRIL Fab^55^. Not only did cryoEV increase the variety of particle orientations in the 2D class averages, similar to earlier experiments, but it also increased the number of Fab-bound complexes, while the no additive sample primarily contained free Fab fragments (Figure S6F). Given the inherent difficulty in stabilizing inactive GPCR complexes, the enhancement in the stability of commonly used BRIL-fusion constructs is valuable for structure determination.

Overall, we observed that cryoH was particularly efficacious as a general stabilizer for multi-protein assemblies across several test cases (Table 1). We do note, however, that the use of cryoH is often associated with a modest reduction in overall particle density, as observed in our CCAN and TRIP12-USP7 datasets (Figure 4A, 4C and S6A). Nonetheless, the benefit of the complex stabilization provided by cryoH far outweighs the reduction in particle count.

### CryoSHIELDs effect on optimized samples

We investigated whether cryoSHIELDs affect “routine” targets, i.e. samples amenable to high-resolution structure determination without overt preparation challenges. Using the large E3 ubiquitin ligase HUWE1 as a model system^56,57^, we systematically compared particle distribution, angular sampling, and reconstruction quality across equivalent datasets prepared with different additives (Figures 4E–4G and S7A). In the absence of additives, HUWE1 yielded a 4.6 Å reconstruction, whereas cryoH, cryoM, and cryoKF improved the resolution to 3.44 Å, 3.23 Å, and 3.22 Å, respectively (Figure S7B). Importantly, these improvements were accompanied by substantially better-resolved structural features, as demonstrated by representative atomic model-to-map fits, providing direct evidence that cryoSHIELDs improve structural interpretability beyond simply increasing particle numbers (Figure S7C). The LEA protein similarly improved resolution to 3.42 Å, whereas cryoRF induced aggregation and reduced resolution to 7.51 Å. We speculate this performance gain could be attributed to three factors: (i) thinner, more uniform ice, evident from improved micrograph contrast at the same defocus (Figure S7A); (ii) enhanced preservation of intact particles by cryoSHIELDs, consistent with their stabilizing effects on protein complexes and reflected in particle counts per micrograph (Figure 4G); and (iii) improved distribution of views for certain cryoSHIELDs, demonstrated by improved cFAR values (Figure 4G), similar to that observed for other protein samples (*e.g.* Figure 1). By optimizing these parameters simultaneously, cryoSHIELDs augment the quality of even “standard” cryo-EM specimens, thereby maximizing data utility and reducing required microscope time.

Lastly, we identified a performance “ceiling” for inherently ideal cryo-EM specimens. Adeno-associated viruses (AAVs), which naturally partition into thin ice with near-perfect angular distributions, derived no additional benefit from cryoSHIELD inclusion (Figure S7D). Curiously, peptides induced some degree of AAV preferred orientation, consistent with cryoSHIELDs modifying the AWI. These results suggest that while cryoSHIELDs are not required for specimens already perfectly suited for cryo-EM, they represent a powerful optimization tool for the vast majority of other biological targets limited by lack of symmetry, AWI-mediated instability, or orientation bias.

### Systematic comparison of cryoSHIELD effects

To systematically evaluate the impact of all cryoSHIELDs and peptide combinations on particle distribution and interaction with the AWI, we used a phosphoinositide 3-kinase-alpha (PI3Kα) complex bound to the small-molecule inhibitor, GDC-0077 (Inavolisib), as a model system^58^ (Figure 5A, S8, and S9). For each condition, we collected comparable single-particle datasets, applied minimal processing to obtain 2D classes and view distribution plots, and acquired tomograms from equivalent grid regions to assess particle distribution within the ice. Without additives, PI3Kα exhibited preferred orientation, similar to previously reported highly anisotropic cryo-EM maps^59^, making it a challenging case with poorly defined protein densities (Figure 5B). Although conventional detergents (e.g., fos-choline-12), LEA protein, and BH cryo-tide shifted view distributions, consistent with their activity at the AWI, none enabled isotropic reconstructions (Figure 5A and S8). Similarly, individual cryoSHIELDs altered the distribution of views, also consistent with cryoSHIELDs functioning at the AWI. However, single peptides were insufficient to achieve isotropic reconstructions for the PI3Kα complex. To overcome these limitations, we explored combinations of peptides. Similar to the cryoSHIELD mixtures being useful for Fab-antigen and other protein complexes, the most effective condition for PI3Kα complex was a combination of linear cryoEV and helical cryoH, which markedly improved the cFAR score from 0.04 without peptide to 0.38 for the combination and resulted in substantially improved cryo-EM density (Figure 5B, 5C and S8). Interestingly, this improvement did not reflect a simple additive combination of the orientations observed for each peptide individually. Instead, the cryoEV–cryoH mixture produced a distinct distribution of views, with 2D classes differing from those dominant in the cryoEV-only condition, indicating that combining linear and helical cryoSHIELDs allows access to new regions of orientation space.

**Figure 5.**
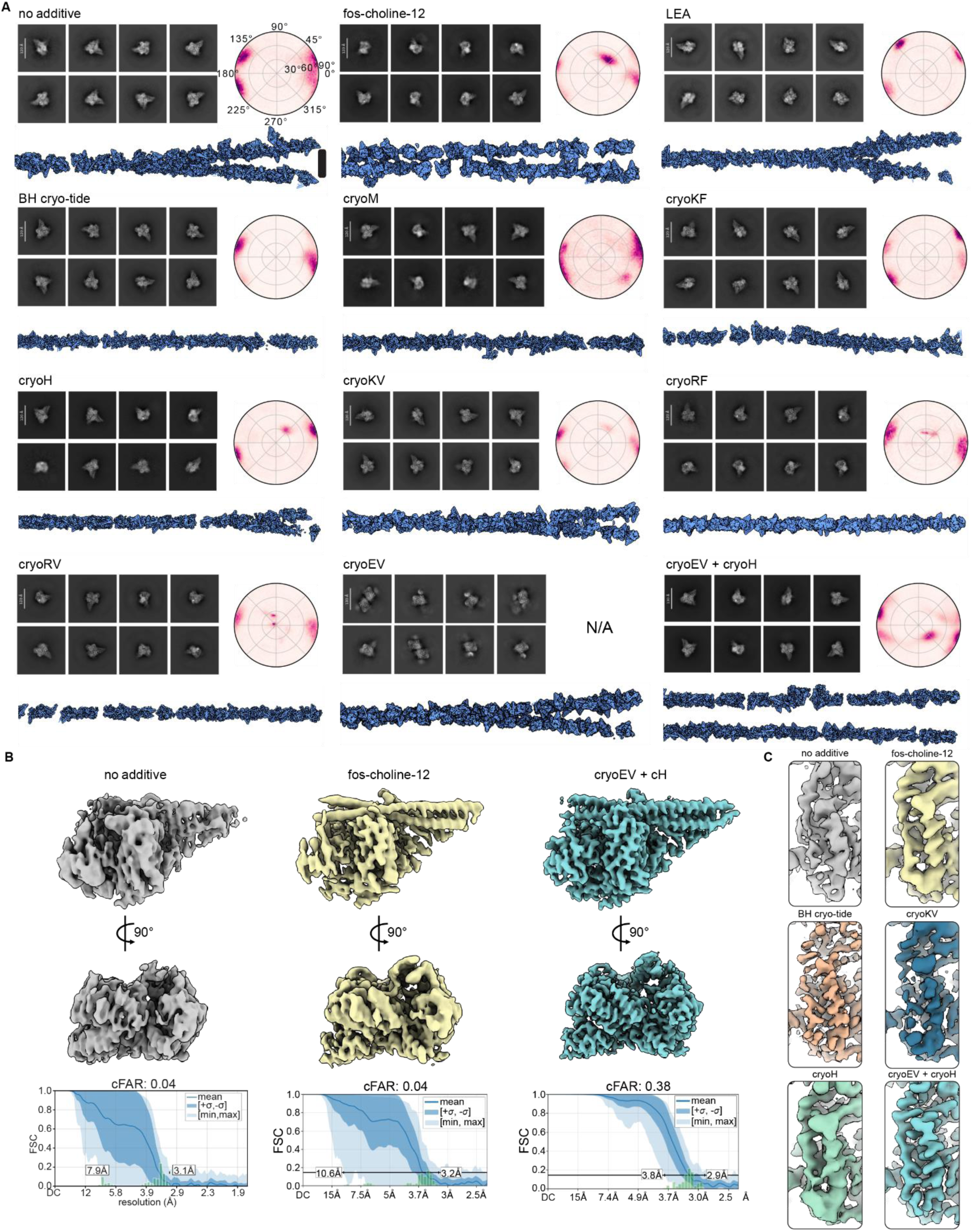
Effect of additives on PI3Kα complex. (A) Analysis of PI3Kα particle behavior under different additive conditions. Top left: representative 2D class averages. Scale bars in 2D averages are indicated in each panel. Top right: viewing direction distributions. Bottom: cross-sectional schematic diagrams illustrating the spatial distribution of particles within vitrified ice holes. Particle poses are estimates derived from template matching; each diagram represents a cross-section spanning approximately half of the foil hole (center to edge, right to left). The cryoEV dataset did not yield sufficient orientations to produce a 3D map. Scale bar in the first panel is 25 nm, the thickness of ANT 1.2/1.3 grids. (B) Comparison of reconstructed maps obtained for selected conditions (top) and corresponding conical FSC Area Ratio (cFAR) plots and scores (bottom). (C) A close-up view of the cryo-EM density map of the PI3Kα complex reconstructed in the presence of different additives.

We also used the PI3Kα complex as a case study to examine how cryoSHIELDs influence particle distribution within vitrified ice and local ice thickness. In the tomograms examined, most individual cryoSHIELDs were associated with particles arranged predominantly in a single interfacial layer, often with a shallow ice meniscus toward the hole center (Figure 5A and S9). Because these observations were derived from a single grid per condition, and ice thickness was estimated from interparticle spacing, we interpret these measurements as useful local ice thickness estimates to correlate with single-particle analysis rather than universally reproducible effects of individual peptides. Nevertheless, several distinct behaviors emerged. Combinations of linear and helical peptides exhibited more complex behavior, in some cases promoting particle partitioning into two layers consistent with increased ice thickness (e.g., cryoKV with either helical peptide, and cryoEV with cryoH), whereas other combinations (e.g., cryoKF with either helical peptide) retained a uniform monolayer. CryoEV showed substantial variation in estimated ice thickness across the sampled holes, although whether this reflects an intrinsic property of the peptide or variability in grid preparation remains unclear. CryoRV similarly confined PI3Kα particles predominantly to a single layer; however, additional density was observed at the opposite interface, potentially corresponding to peptide assemblies or aggregates of uncertain composition. Despite the estimated increased ice thickness observed in the cryoEV-cryoH combination, this condition yielded the most isotropic reconstruction (Figure 5B and S8). In this condition, particles occupied two distinct layers separated by approximately 60 nm, suggesting that alternative adsorption geometries, rather than prevention of interfacial adsorption, may contribute to improved angular sampling induced by the peptides. Together, these observations demonstrate that cryoSHIELDs modify, rather than abolish, interfacial particle organization and local ice geometry in distinct ways. Our observations suggest that while individual cryoSHIELDs can introduce new particle orientations, their distinct behaviors provide a modular framework for mix-and-match optimization. Consistent with our observations across other orientation-limited systems, these results suggest that combining a linear peptide (cryoKV or cryoEV) with a helical peptide (cryoH or cryoM) provides a particularly effective and broadly applicable starting point for soluble protein cryo-EM grid optimization, while allowing further refinement for individual targets.

We also examined the concentration dependence of the cryoEV–cryoH mixture on the angular distribution of PI3Kα particles (Figure S10A). Increasing peptide concentration progressively altered the distribution of particle orientations, accompanied by improved cFAR scores and corresponding improvements in the cryo-EM reconstructions. The effect became pronounced at 50 µM of each peptide and increased further at higher peptide concentrations. Thus, as observed for the Fab–antigen complex described above (Figure S2F–H), the effects of cryoSHIELDs on particle orientation were concentration dependent, with the strongest improvements observed at the higher peptide concentrations tested.

Given that PI3Kα particles remained localized at the AWI interface, we next investigated whether direct peptide–protein interactions contribute to the observed changes in particle orientation. We performed surface plasmon resonance (SPR) experiments using immobilized PI3Kα, validating the assay with a phosphorylated PDGFR-derived peptide (740Y-P), which exhibited concentration-dependent binding consistent with previous reports^60^ (Figure S10B). Most cryoSHIELDs showed little or no detectable association at the concentrations tested (up to 10 µM), whereas cryoM exhibited appreciable concentration-dependent binding and cryoKF produced weaker, noncanonical apparent association profiles (Figure S10B). Interestingly, the cryoEV– cryoH combination, which substantially improved PI3Kα angular sampling (cFAR from 0.04 to 0.38), showed no appreciable binding under these conditions. These observations suggest that detectable peptide–protein binding is not required for improved angular sampling, although interactions at higher peptide concentrations or specifically at the peptide-modified AWI cannot be excluded. Conversely, the association observed for cryoM and cryoKF may contribute to their tendency to induce distinct views. To assess whether cryoSHIELDs influence protein stability, we performed nano-differential scanning fluorimetry (nanoDSF) at peptide concentrations used for cryo-EM (200 µM). While inavolisib substantially increased PI3Kα thermal stability, cryoSHIELDs produced negligible changes (Figure S10C and S10D). Together, these results suggest that cryoSHIELD-mediated improvements in PI3Kα cryo-EM behavior are not explained by thermal stabilization in solution, while peptide-specific interactions may contribute to the effects of individual additives.

### CryoSHIELD effects on CTF estimation and FFT signal

To assess whether cryoSHIELDs themselves could interfere with cryo-EM imaging and data processing, we prepared grids containing individual peptides in the absence of target proteins, alongside buffer-only, control peptide, and conventional detergent conditions (Figure S11). Inspection of representative micrographs revealed no obvious peptide-induced artifacts or large-scale assemblies (Figure S11A). We subsequently collected screening datasets for each condition and evaluated their effects on contrast transfer function (CTF) estimation and relative ice thickness as determined by cryoSPARC (Figures S11B–S11D). Interestingly, analysis of the Fourier transforms and their one-dimensional rotational averages revealed a distinct diffraction peak at approximately 4.7 Å in grids containing linear cryoSHIELDs (cryoKV, cryoEV, and cryoKF), which was absent from helical peptides, control peptides, and conventional detergents (Figures S11E and S11F). This periodicity is characteristic of the inter-strand spacing in β-sheets, suggesting that linear cryoSHIELDs formed ordered peptide assemblies. Importantly, this diffraction feature did not prevent reliable CTF estimation or high-resolution reconstruction of protein structures.

To investigate whether this β-sheet organization was intrinsic to the peptides or associated with cryo-EM grid preparation, we characterized their secondary structures in solution using circular dichroism (CD) spectroscopy (Figure S12A). Most cryoSHIELDs exhibited predominantly random coil conformations in solution, with only cryoKF showing some β-sheet propensity. Together with the diffraction measurements, these observations are consistent with the idea that linear cryoSHIELDs undergo structural reorganization into ordered β-sheet assemblies during grid preparation, potentially driven by their interaction with the AWI. Although these experiments do not directly establish peptide localization at the interface, they are consistent with peptides functioning at the AWI.

Under certain conditions, linear cryoSHIELDs also produced larger, micron-scale sheet-like artifacts, particularly following prolonged storage at 4°C (>2 weeks) or extensive agitation (Figure S12B). The previously described amphipathic peptide BH cryo-tide also formed extended filamentous assemblies exhibiting a characteristic 4.7 Å diffraction feature, consistent with β-sheet-like molecular organization (Figure S12C). Two-dimensional classification of these filaments revealed distinct patterns of periodic striations, suggesting that the peptides can assemble into ordered structures with different packing arrangements (Figure S12D). These observations highlight the importance of peptide handling and stock preparation (detailed in the Methods section). Among the cryoSHIELDs tested, cryoKF required the greatest concentration optimization, frequently inducing aggregation of proteins at 200 µM, particularly at 4°C (Figure S1E). This effect could be mitigated by reducing the peptide concentration to 50–100 µM and adding it at room temperature immediately before vitrification. This approach retains its potent particle-enrichment effects on detergent-solubilized membrane proteins (Figures 2 and 3). In contrast, cryoRF and cryoRV frequently promoted protein aggregation even at reduced concentrations, limiting their general applicability (Figures S1B and S1D). Together, these findings identify practical considerations for cryoSHIELD use.

## DISCUSSION

We have developed and characterized cryoSHIELDs, a library of amphipathic peptide additives that mitigate common challenges in cryo-EM sample preparation across diverse biologically and therapeutically relevant protein systems. Evaluated across 24 targets, cryoSHIELDs improved particle recovery, angular sampling, preservation of labile protein complexes, and, in selected cases, reconstruction quality and resolution. Direct structural comparisons also demonstrate that these benefits extend beyond changes in particle distribution to improvements in map interpretability, facilitating high-resolution structure determination of previously challenging targets.

We observed distinct, target-dependent activities within the cryoSHIELD toolbox (Figure 6A). Linear peptides cryoKV and cryoEV were particularly effective at diversifying particle orientations, whereas cryoKF enabled substantial enrichment of detergent-solubilized membrane proteins, albeit with a propensity to aggregate certain soluble complexes. Among helical peptides, cryoM frequently introduced unique particle orientations, while cryoH was particularly effective at preserving fragile multiprotein assemblies and modifying the orientation of membrane proteins in lipid nanodiscs.

**Figure 6.**
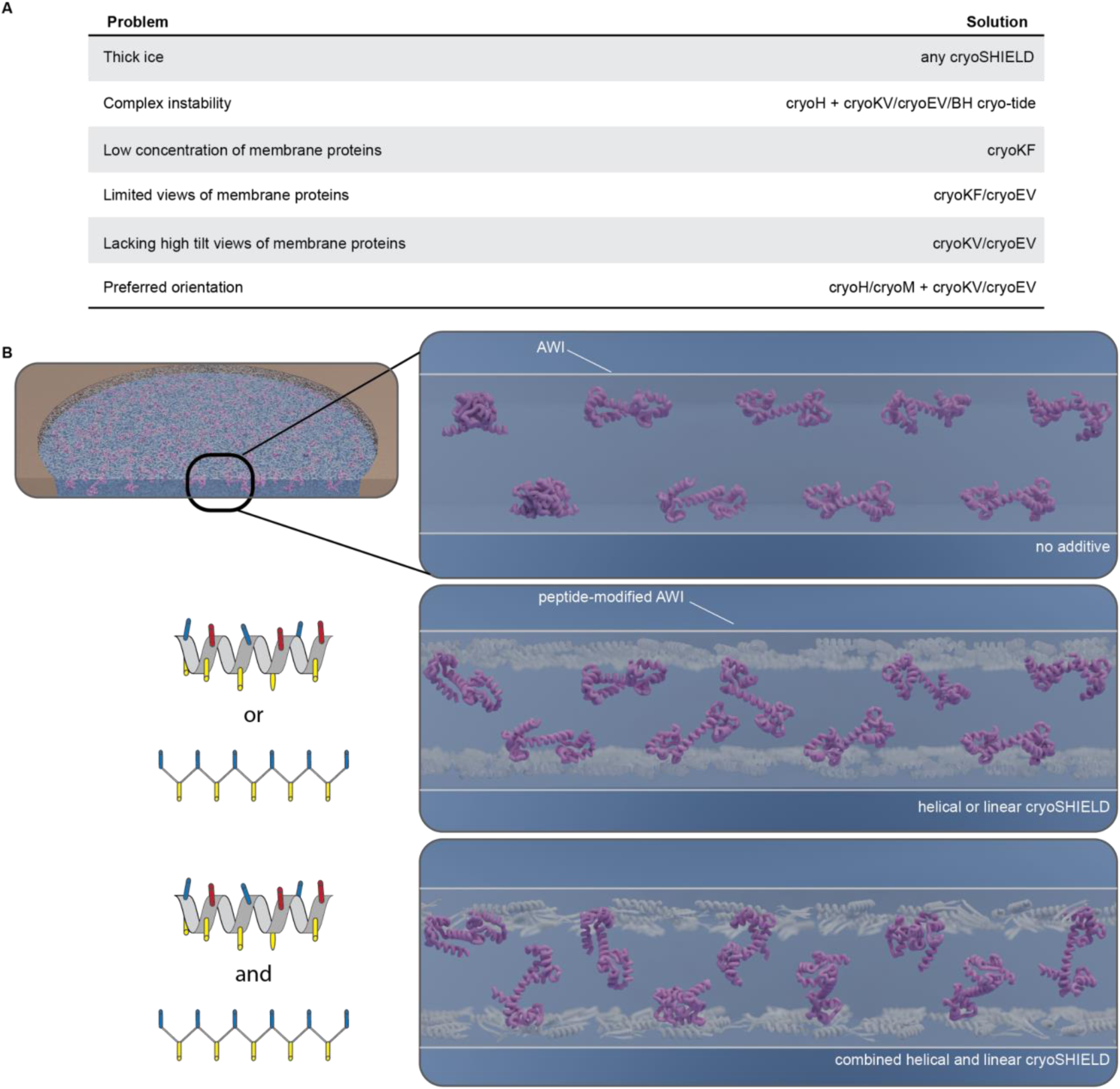
Mechanism of cryoSHIELD function. (A) Table of common cryo-EM sample preparation problems that cryoSHIELDs can address and initial recommendations based on the data in this paper. (B) Schematic illustration of the proposed mechanism of cryoSHIELD activity at the air–water interface (AWI). Without additives, particles may adsorb directly to the AWI, frequently adopting restricted orientations and/or denaturing. CryoSHIELDs are proposed to modify the interfacial environment, allowing particles to interact with the peptidic surface while potentially preserving their structural integrity and promoting alternative adsorption geometries. Different peptide structures and combinations may generate distinct interfacial environments, enabling particles at opposing interfaces to adopt complementary orientations and thereby improving angular sampling without necessarily eliminating preferred orientation.

The benefits of these additives are often synergistic, particularly when pairing linear and helical variants. Binary combinations, such as cryoKV + cryoH/M, cryoEV + cryoH/M, or cryoKF + cryoH/M, proved highly versatile and represent a powerful starting point for sample optimization. Although final peptide selection remains empirical, our extensive characterization provides target-class-dependent recommendations that considerably reduce the number of conditions required in initial screening. CryoSHIELDs have consequently become a routine component of our cryo-EM workflow, with applications extending to more than 40 protein systems. Overall, cryoSHIELDs provide a cost-effective, high-throughput, and target-class-dependent toolkit for overcoming the sample-specific bottlenecks that have historically limited the pace of cryo-EM structure determination (Fig. S13).

Mechanistically, our observations are consistent with cryoSHIELDs primarily modifying protein interactions with the AWI, rather than preventing interfacial adsorption (Figure 6B, Movie S1). Cryo-electron tomography revealed that particles frequently remain associated with one or both interfaces, with certain peptide combinations promoting distinct adsorption geometries that contribute to improved angular sampling. Consistent with their surface activity, linear cryoSHIELDs exhibited a characteristic ∼4.7 Å diffraction signal on cryo-EM grids, suggestive of ordered β-sheet-like assemblies, whereas CD measurements indicated predominantly disordered conformations in solution (Figures S11 and S12). Furthermore, SPR measurements revealed little detectable association between PI3Kα and most peptides, although cryoM and cryoKF exhibited appreciable interactions (Figure S10). Notably, the cryoEV–cryoH combination substantially improved angular sampling without apparent SPR binding, suggesting that stable peptide–protein interactions are not present or essential for this effect. NanoDSF measurements likewise showed no substantial stabilization of PI3Kα in solution. Together, these findings support a model in which cryoSHIELDs create a modified interfacial environment that can preserve particle integrity and permit alternative adsorption orientations, with direct peptide–protein interactions potentially contributing to the effects of individual additives. Although direct localization and molecular organization of cryoSHIELDs at the AWI remain unresolved, this model provides a framework for understanding their complementary activities and can guide further optimization.

### Limitations of the Study

Although cryoSHIELDs were evaluated across 24 diverse protein systems, their effects remain target-dependent, and no single peptide universally ‘solves’ cryo-EM sample preparation. Our attempts to correlate protein physicochemical properties, including hydrophobicity, surface charge distribution, and isoelectric point, with cryo-EM performance did not identify robust predictive relationships, although related classes of proteins often exhibited similar responses to individual cryoSHIELDs. Thus, our recommendations provide empirically informed starting points rather than a fully predictive framework. Certain peptides can also promote protein aggregation or exacerbate preferred orientation, as observed for cryoKF with some soluble complexes and for AAV particles, respectively. Reducing peptide concentration can mitigate aggregation in some cases, although this may not resolve all adverse effects. Furthermore, peptide CSC values determined in PBS may not directly translate to other buffer conditions. The molecular mechanism of cryoSHIELD activity also remains incompletely understood. Although tomography, diffraction, and biophysical measurements are consistent with modification of particle adsorption at the air– water interface, the localization and molecular organization of the peptides at the interface have not been directly visualized. Furthermore, potential peptide–protein interactions were examined primarily using PI3Kα and cannot be excluded for other targets or under interfacial conditions. Finally, cryo-EM grid preparation is inherently variable, and certain analyses, particularly tomography and indirect ice-thickness estimation, were based on limited sampling and may not fully capture grid-to-grid variability. Two examples we have observed of apparent failure cases are the TNF family trimer bound to 3-Fabs in a C3 symmetric organization in Table 1 and a Fab-MHC complex sample. In both cases, single and combinations of cryoSHIELDs were ineffective at resolving the observed preferred orientation. Other sample modifications such as detergents, other additives, sample PEGylation, and graphene oxide supports were also ineffective in these cases. These limitations highlight the need for further mechanistic characterization of cryoSHIELD activity, development of new cryoSHIELD peptides, and the need for guided target-specific optimization.

## Resource Availability

Further information and requests for resources and reagents should be directed to the lead contact, Stanislau Yatskevich.

## Materials Availability

Unique reagents generated in this study may be requested through Genentech’s Material Transfer Agreement (MTA) program at https://www.gene.com/scientists/mta. Requests are subject to material availability, approval, and execution of an MTA prior to transfer. Questions regarding material requests or MTA documentation should be directed to.

## Data and code availability

● Cryo-EM maps and atomic coordinates generated in this study have been deposited in the Electron Microscopy Data Bank (EMDB) and Protein Data Bank (PDB). For ABCC6, the NBD1-focused refinement (EMD-77250), NBD2-focused refinement (EMD-77249), consensus refinement (EMD-77251), and composite map (EMD-77282) have been deposited in the EMDB, and the refined atomic coordinates have been deposited in the PDB (PDB: 35XZ). For PI3Kα, reconstructions obtained in the presence of cryoEV– cryoH (EMD-77258) and in the absence of cryoSHIELDs (EMD-77259) have been deposited in the EMDB. The OSMRβ–Vixarelimab-derived Fab complex prepared with cryoEV–cryoH has been deposited as EMD-79601, with corresponding atomic coordinates deposited as PDB: 39MH. For the DAMGO-bound μ-opioid receptor–Gi complex, consensus and locally refined maps obtained at 1.0 mg/mL with cryoKF have been deposited as EMD-77232 and EMD-77231, respectively; consensus and locally refined maps obtained at 0.5 mg/mL with cryoKF have been deposited as EMD-79600 and EMD-7998, respectively; and consensus and locally refined maps obtained at 5 mg/mL without peptide have been deposited as EMD-79365 and EMD-79366, respectively.
● This paper does not report original code.
● Any additional information required to re-analyze the data reported in this paper is available from the lead contact upon request.

## Acknowledgement

The authors thank Gladys Boenig, Amy Byrnes, Rina Fong, Ireneusz Gorniak, Hoangdung Ho, Angela Oh, Sergei Pourmal, Dawei Sun, Tim Wendorff, and Ping Wu for providing samples used for cryoSHIELD testing. We also thank Wenping Chang for help with peptide synthesis. We are grateful to the Biomolecular Resources group for their help with molecular biology across projects. Additionally, we thank Bryan Martin for early collaborations and insightful discussions regarding peptide characterization.

## Author contribution

S.Y. conceived the study. Methodology: C.M.A, P.D., M.C.J., S.Y.; Software: A.B.; Validation: C.M.A., P.D., M.C.J, S.Y.; Formal Analysis: C.M.A, P.D., M.C.J., S.Y, B.D.D, C.W.K., T.H.S.; Investigation: J.Y.K.T., A.S. (peptide synthesis), S.T., N.M., S.K.R., C.W., C.W.K., B.D.D., S.Y. (sample preparation), C.M.A., P.D., M.C.J., B.D.D., C.W.K., T.H.S., O.E.M., S.Y. (cryo-EM data collection), S.K.R., F.A. (biophysical data collection); Resources: A.R., C.C.; Data Curation: C.M.A, P.D., M.C.J, S.Y.; Writing – Original Draft: C.M.A., P.D., M.C.J., S.Y; Writing – Review & Editing: all authors; Visualization: C.M.A, P.D., M.C.J., A.C.H., S.Y; Supervision: A.R., C.C., S.Y.; Project Administration: C.M.A.

## Declaration of Generative AI and AI-assisted technologies in the writing process

During the preparation of this work, the authors used ChatGPT 5.5 or 6.0. Thinking to assist with language editing and proofreading. After using this tool or service, the authors reviewed and edited the content as needed and take full responsibility for the content of the publication.

## STAR★Methods

### Key resource table

| **REAGENT or RESOURCE** | **SOURCE** | **IDENTIFIER** |
|---|---|---|
| **Antibodies** |  |  |
| Recombinant scFv16 antibody fragment | This work; Maeda et al.<sup>70</sup> | See methods |
| Vixarelimab-derived Fab (Vixa.4DS) against OSMRbeta | This work | See methods |
| Anti-FLAG M2 agarose affinity resin | Sigma-Aldrich | A2220 |
| **Chemicals, peptides, and recombinant proteins** |  |  |
| cryoKV amphipathic peptide | GenScript (custom synthesis); this paper | Sequence: KAKVKVKVKVKVKVKVKAK; Table S1 |
| cryoMAX3 amphipathic peptide | GenScript (custom synthesis); this paper | Sequence: VKVKVKTKVPPTKVKTKVKV; Table S1 |
| cryoEV amphipathic peptide | This paper (in-house solid-phase synthesis) | Sequence: EAEVEVEVEVEVEVEVEAE; Table S1 |
| cryoKF amphipathic peptide | GenScript (custom synthesis); this paper | Sequence: KAKFKFKFKFKFKFKFKAK; Table S1 |
| cryoKL amphipathic peptide | GenScript (custom synthesis); this paper | Sequence: KAKLKLKLKLKLKLKLKAK; Table S1 |
| cryoRV amphipathic peptide | GenScript (custom synthesis); this paper | Sequence: RARVRVRVRVRVRVRVRAR; Table S1 |
| cryoRF amphipathic peptide | GenScript (custom synthesis); this paper | Sequence: RARFRFRFRFRFRFRFRAR; Table S1 |
| cryoW1 amphipathic peptide | GenScript (custom synthesis); this paper | Sequence: MIKVKLNWKFKVNFEINVKI; Table S1 |
| cryoW2 amphipathic peptide | GenScript (custom synthesis); this paper | Sequence: KWKALWKLLESLWKLLWEAE; Table S1 |
| 3xFLAG peptide | Sigma-Aldrich | F4799 |
| cryoH amphipathic peptide | GenScript (custom synthesis); this paper | Sequence: KLQELQEKLTPLGEDLRDRMRHHVDALRTKMT; Table S1 |
| cryoM amphipathic peptide | GenScript (custom synthesis); this paper | Sequence: KWKVFKKIEKMGRNIRNGIVKAGPAIAVLGEAKAL; Table S1 |
| cryoB amphipathic peptide | GenScript (custom synthesis); this paper | Sequence: KWKLFKKIGAVLKVL; Table S1 |
| cryoCHARGE amphipathic peptide | GenScript (custom synthesis); this paper | Sequence: KLKEEEERQKREEEERIKRLEELEAKRKEEERLEQEKRERK; Table S1 |
| BH cryo-tide | Genscript (custom synthesis); citation<sup>34</sup> | Sequence: MALKVTKNSKINAENKAKINM; Table S1 |
| PDGFR-derived 740Y-P phosphorylated peptide | Selleckchem | S7865 |
| Inavolisib (GDC-0077) | This paper (in-house produced) | N/A |
| DAMGO agonist peptide | MedChemExpress | HY-P0210 |
| Biotin-LPXTG sortase labeling reagent | This paper (in-house produced) | N/A |
| Sortase A enzyme | This paper (in-house produced) | N/A |
| TEV protease | This paper (in-house produced) | N/A |
| Benzonase nuclease | This paper (in-house produced) | N/A |
| Apyrase | New England Biolabs | M0398L |
| Lauryl maltose neopentyl glycol (LMNG) | Anatrace | NG310 |
| Cholesteryl hemisuccinate tris salt (CHS) | Anatrace | CH210 |
| Glyco-diosgenin (GDN) | Anatrace | GDN101 |
| Digitonin | Cayman Chemical | 14952 |
| CHAPSO | Anatrace | C317 |
| PEIpro transfection reagent | Sartorius | 101000029 |
| Sheared herring sperm DNA | Promega | D181A |
| Expi293 Expression Medium | Gibco | A1435101 |
| Fos-choline-12 | Anatrace | F308 |
| IGEPAL CA-630 | Sigma-Aldrich | I8896 |
| Triton X-100 | Thermo Fisher Scientific | 28314 |
| Triton X-1114 | Thermo Fisher Scientific | 28332 |
| Purified recombinant late embryogenesis abundant (LEA) protein | This paper, based on Abe et al.<sup>32</sup> | See Methods |
| Purified recombinant PI3Kalpha heterodimer: p110alpha P3-N1068 / p85alpha A318-N600 | This paper | See Methods |
| Purified recombinant mu-opioid receptor OPRM1 (F158W)-GNAI1/B1/G2-scFv16complex | This paper | See Methods |
| Purified recombinant ABCC6 tagged transporter | This paper | See Methods |
| Purified recombinant OSMRbeta-vixarelimab-derived Fab complex | This paper | See Methods |
| Purified recombinant HUWE1 E3 ubiquitin ligase sample | Yatskevich et al<sup>57</sup> | N/A |
| Purified recombinant Rad55-Rad57-SHU complex | Koo et al<sup>71</sup> | N/A |
| Purified recombinant BCDX2-CX3-RAD51 complex | Koo et al<sup>72</sup> | N/A |
| **Deposited data** |  |  |
| ABCC6 NBD1 focused cryo-EM map | EMDB | EMD-77250 |
| ABCC6 NBD2 focused cryo-EM map | EMDB | EMD-77249 |
| ABCC6 consensus cryo-EM map | EMDB | EMD-77251 |
| ABCC6 composite cryo-EM map | EMDB | EMD-77282 |
| ABCC6 refined atomic coordinates | PDB | PDB: 35XZ |
| mu-opioid-Gi (5 mg/mL): consensus map | EMDB | EMD-79365 |
| mu-opioid-Gi (5 mg/mL): local map | EMDB | EMD-79366 |
| mu-opioid-Gi (0.5 mg/mL, cryoKF): consensus map | EMDB | EMD-79600 |
| mu-opioid-Gi (0.5 mg/mL, cryoKF): local map | EMDB | EMD-79982 |
| mu-opioid-Gi (1.0 mg/mL, cryoKF): consensus map | EMDB | EMD-77232 |
| mu-opioid-Gi (1.0 mg/mL, cryoKF): local map | EMDB | EMD-77231 |
| PI3Kalpha + cryoEV-cryoH reconstruction | EMDB | EMD-77258 |
| PI3Kalpha without peptide reconstruction | EMDB | EMD-77259 |
| OSMR in complex with rigid Vixarelimab Fab with cryoEV-cryoH, cryo-EM map | EMDB | EMD-79601 |
| OSMR in complex with rigid Vixarelimab Fab with cryoEV-cryoH, coordinates | PDB | 39MH |
| Molecular movie of proposed cryoSHIELD function | This paper | Movie S1 |
| **Experimental models: Cell lines** |  |  |
| Hi5 insect cells (Trichoplusia ni) | Gibco | B85502 |
| Expi293F suspension cells | Gibco | A14527 |
| CHO cells | Gibco | R80007 |
| **Recombinant DNA** |  |  |
| pAcGP67A PI3Kalpha (p110alpha-p85alpha-T2A) expression construct | This paper | See methods |
| OPRM1(F158W)-LgBiT-TEV-vGFP-twin-Strep plasmid | This paper | See methods |
| GNAI1(S47C/G202T/G203A/E245A/A326S) plasmid | This paper | See methods |
| GNB1(His8,HiBiT)-GNG2 bicistronic plasmid | This paper | See methods |
| scFv16-His8 expression plasmid | This paper | N/A |
| pRK5 ABCC6-TEV-eGFP-3xFLAG plasmid | This paper | See methods |
| OSMRbeta expression construct | This paper | See methods |
| Vixarelimab-derived Fab expression plasmid(s) | This paper | See methods |
| **Software and algorithms** |  |  |
| CryoSPARC, including cryoSPARC Live | Punjani et al.<sup>73</sup>; Structura Biotechnology | https://cryosparc.com |
| AlphaFold2 | Jumper et al.<sup>74</sup> | https://github.com/google-deepmind/alphafold |
| Coot v0.9.6 | Emsley et al.<sup>75</sup> | https://www2.mrc-lmb.cam.ac.uk/personal/pemsley/coot/ |
| PHENIX v2.0 | Adams et al.<sup>76</sup> | https://phenix-online.org |
| UCSF ChimeraX v1.20 | Pettersen et al.<sup>77</sup> | https://www.cgl.ucsf.edu/chimerax/ |
| SmartScope | Bouvette et al.<sup>78</sup> | https://github.com/NIEHS/SmartScope |
| SerialEM | Mastronarde<sup>79</sup> | https://bio3d.colorado.edu/SerialEM/ |
| Etomo / IMOD | Kremer et al.<sup>80</sup> | https://bio3d.colorado.edu/imod/ |
| IsoNet denoising software | Liu et al.<sup>81</sup> | https://github.com/IsoNet-cryoET/IsoNet2 |
| GraphPad Prism 11 | GraphPad Software | https://www.graphpad.com/scientific-software/prism/ |
| Orientation distribution plots | This paper | https://gist.github.com/alisterburt/7bcdfc902213434ded8c8aa0770744f2 |
| **Other** |  |  |
| Sensor Chip SA (streptavidin) | Cytiva | Cat# BR100032 |
| HisTrap Excel Ni affinity column | Cytiva | 17371205 |
| Strep-Tactin XT 4Flow affinity resin | IBA | 2-5010-010 |
| QF 1.2/1.3, Cu 300 mesh cryo-EM grids | Quantifoil | N1-C14nCu30-01 |
| UF 1.2/1.3, Au 300 mesh cryo-EM grids (25-nm foil) | Quantifoil | N1-A14dAu30-01-t25 |
| ANT 1.2/1.3, Au 300 mesh cryo-EM grids | Molecular Dimensions | MDL-M01 |
| ExpiFectamine 293 Transfection Kit | Thermo Fisher Scientific | A14525 |
| EDTA-free protease inhibitor tablets | Roche | 11836170001 |

### Experimental models and study participant details

Human Expi293F cells (Thermo Fisher Scientific) were used for recombinant expression of the μ-opioid receptor–Gi complex, ABCC6 and TRIP12-USP7. Cells were maintained in Expi293 Expression Medium at 37°C and transiently transfected using the ExpiFectamine 293 transfection kit (Thermo Fisher Scientific) according to the manufacturer’s instructions. Transfections were performed at cell densities of 3.0–3.5 × 10⁶ cells/mL. Following transfection, cultures were supplemented with 2 mM valproic acid and 5 mM sodium propionate after 16 h and maintained for an additional 24 h before harvesting.

Trichoplusia ni Hi5 insect cells were used for recombinant expression of the human PI3Kα complex using the baculovirus expression system. Cells were infected with recombinant baculovirus at a multiplicity of infection (MOI) of 1 and maintained at 27°C for 48 h before harvesting.

Chinese hamster ovary (CHO) cells were used for recombinant expression and secretion of scFv16 and the engineered Vixarelimab Fab. Fab heavy- and light-chain expression constructs were transiently co-transfected into CHO cells using PEIpro reagent.

## Method Details

### Peptide synthesis

All peptides except cryoEV were synthesized by GenScript at >95% purity without N- or C-terminal modifications. Initial screening used peptides without trifluoroacetic acid exchange; subsequent batches prepared with trifluoroacetic acid exchange behaved equivalently. Multiple independently synthesized batches were tested and produced consistent results for each peptide.

The cryoEV peptide was synthesized using automated microwave-assisted solid-phase 9-fluorenylmethyloxycarbonyl (Fmoc) chemistry, which was performed on preloaded TenteGel resin (0.20 mmol/g, 0.5g) following a published procedure^82^. The obtained crude peptide (156 mg) was analyzed by liquid chromatography-mass spectrometry (LC-MS, C18 column, 4.6 mm × 50 mm, 5 μm, 100 Å, 1.0 mL/min, 2-min gradient from 5 to 95% aqueous acetonitrile containing 10 mM ammonium bicarbonate, 220 nm) and purified by reverse-phase high-performance liquid chromatography (RP-HPLC, C18 column, 21 mm × 100 mm, 5 μm, 100 Å, 20 mL/min, 30-min gradient from 0 to 20% aqueous acetonitrile containing 10 mM ammonium bicarbonate) to afford purified cryoEV as a white lyophilized powder (40 mg, *m*/*z*: [M-2H]^2-^ calculated for C_101_H_161_N_21_O_44_ 1185.5447, found 1185.5428).

Helical peptides, cryoM and cryoH, and cryoEV were prepared in 20 mM HEPES pH 8.0. Linear peptides, cryoKV, cryoKF, cryoKL, cryoRV, and cryoRF, were prepared in 20 mM MES pH 6.0. Peptides were dissolved directly in the buffer solution by gentle tapping. Harsh treatment such as prolonged vigorous pipetting or excessive vortexing promotes aggregation of linear peptides while having little effect on the helical peptides. Peptides were subsequently aliquoted into small batches of 20-40 µL and snap-frozen in liquid nitrogen. Subsequently, individual aliquots were thawed and used within 2 weeks.

Unless otherwise indicated, individual peptides were used at 200 µM, whereas 1:1 peptide mixtures were used at 200 µM total concentration (100 µM each). cryoKF alone was used at 100 µM.

### Marangoni Assay

Peptide stocks at 2 mM were diluted in PBS at 1:10, 1:20, 1:40, 1:80, and 1:160 for initial testing. Peptide dilutions were made directly before 10 µL was pipetted onto freshly cleaved mica. Immediately after, 0.5 µL of nonafluorobuytl methyl ether (NFBME) was applied to the middle of the droplet with a Hamilton syringe and we watched for the Marangoni effect as described in Han and Glaeser 2021^39^. If the final dilution was still exhibiting skin formation and protection from the organic solvent, dilutions of 1:200, 1:400, and 1:600 were tested.

Detergent stocks at 25 mM (above the CMC of all tested detergents) were diluted in PBS at 1:10, 1:20, and 1:40 for initial testing.

### Molecular Biology

All constructs were synthesized by Genscript and validated by either Sanger sequencing or whole plasmid sequencing.

#### PI3Kα

The coding sequences for the human PI3Kα complex subunits, p110α (UniProt ID: P42336) and p85α (UniProt ID: P27986), were engineered for co-expression in a bicistronic format. The catalytic subunit p110α comprised residues P3–N1068, while the regulatory subunit p85α was truncated to include the N-terminal SH2 (nSH2) and inter-SH2 (iSH2) domains (residues A318– N600). Both subunits were cloned into the pAcGP67A vector for expression in insect cells. For simplicity, this construct will be referred to as PI3Kα throughout the text. To promote stoichiometric production of the heterodimer, the two coding sequences were linked via a T2A self-cleaving peptide. A 6×His affinity tag was introduced at the N-terminus of the p85α construct to facilitate purification of the recombinant complex. All constructs were verified by DNA sequencing prior to baculovirus generation.

#### μ-opioid-G_i_ complex

Human OPRM1 (Uniprot ID: P35372) bearing the thermostabilizing F158W mutant and human GNAI1 (Uniprot ID: P63096) with the dominant-negative mutations (S47C, G202T, G203A, E245A, A326S) were each cloned into a mammalian vector using a CMV promoter. OPRM1 contained an N-terminal HA signal sequence and Flag tag as well as C-terminal LgBiT, TEV protease site, the Venus variant of the green fluorescent protein (vGFP), and twin-strep tags. GNAI1 was cloned with no tags. Human GNB1 (Uniprot ID: P62873) and human GNG2 (Uniprot ID: P59768) were cloned into a bi-cistronic mammalian vector. GNB1 contained a N-terminal His8 tag and C-terminal HiBiT tag, while GNG2 was untagged. scFv16 was cloned into a mammalian expression vector with a C-terminal His8 tag and N-terminal VG1 signal peptide.

#### ABCC6

Human wild-type ABCC6 (Uniprot ID: O95255) was cloned into pRK5 vector. TEV cleavage site followed by eGFP and 3xFLAG tag were added at the C-terminus of ABCC6.

#### TRIP12-USP7

Human USP7 (UniProt ID: Q93009) was cloned into the pRK5 vector with an N-terminal FLAG tag followed by a TEV protease cleavage site. Human TRIP12 (UniProt ID: Q14669) was cloned with a C-terminal TEV-cleavable 3×FLAG tag followed by a Twin-Strep tag into the pRK5 vector.

#### Late embryogenesis abundant protein

Late embryogenesis abundant (LEA) protein from R. varieornatus, amino acids K58-A181 (UniProt ID: A0A0E4AVP3), was cloned with N-terminal 8xHis tag for expression in insect cells into the pAcGP67A vector.

#### Oncostatin M receptor subunit beta

Human oncostatin M receptor β (OSMRβ; UniProt Q99650), comprising domains 1–4 (residues M1–E427), followed by a C-terminal GNS linker and an 8×His tag, was cloned into a mammalian pRK expression vector.

#### Vixarelimab

Rigidified Vixarelimab fab heavy- and light-chain constructs were designed using the rigid Fab technology as previously described^40^ using 4 intrachain disulfides (4DS). Both heavy and light chain constructs contained N-terminal secretion signal peptides, and the heavy chain additionally contained a C-terminal GSGS linker followed by a 6×His tag. The heavy- and light-chain sequences were cloned separately into mammalian pRK expression vectors.

### Protein Expression

#### PI3Kα and late embryogenesis abundant protein

Recombinant PI3Kα and LEA protein was expressed in insect cells using the baculovirus expression system. Trichoplusia ni (Hi5) cells were infected with recombinant baculovirus at a multiplicity of infection (MOI) of 1. Infected cultures were incubated at 27 °C and harvested 48 h post-infection.

#### μ-opioid-Gi complex

All plasmids were transiently co-transfected into Expi293F cells (Gibco) at a 2:5:1 [w:w:w] ratio (OPRM1:GNAI1:GNB1-GNG2) using the ExpiFectamine293 kit (ThermoFisher Scientific) following manufacturer’s protocol. The total DNA amount per liter of cells was kept fixed according to the manufacturer’s recommendation. Expressions ranged from 2-4 L of cells, and the plasmid and transfection reagent amounts indicated were scaled accordingly. Expi293F cells were transfected between 3.0 × 10^6^ to 3.5 × 10^6^ cells/ml; in Expi293 expression media. The cells were incubated at 37 °C for 16 hours, after which point 2 mM (final concentration) valproic acid and 5 mM (final concentration) sodium propionate was added to the cell culture and the cells were cultured at 37 °C for an additional 24 hours. Cells were harvested by centrifugation, washed once with phosphate-buffered saline, flash-vitrified in liquid nitrogen, and stored at −80 °C until use for protein purification.

#### ABCC6 and TRIP12-USP7 complex

ABCC6 plasmid or TRIP12 and USP7 plasmids at 1:1 plasmid ratio were transfected into Expi293F (Gibco) cells using the ExpiFectamine293 kit (Thermo Fisher Scientific) following the manufacturer’s protocol. Expi293F cells were transfected at 3.0 × 10^6^ cells/ml. After transfection, 1 L of cells were incubated at 37 °C for 16 hours, after which 2 mM (final concentration) valproic acid and 5 mM (final concentration) sodium propionate were added to the cell culture, and the cells were cultured at 37 °C for an additional 24 hours. Cells were harvested by centrifugation, washed once with phosphate-buffered saline, flash-vitrified in liquid nitrogen, and stored at −80°C until use for protein purification.

#### Vixarelimab, scFv16, and oncostatin M receptor subunit beta

Rigid vixarelimab fab and scFv16 were expressed by transient transfection in CHO-K1 derived cells (ATCC, CCL-61). Oncostatin M receptor subunit beta was expressed in HEK293 cells. Briefly, per 1 Liter of cell culture at a density of 12 million cells/mL, the transfection mixture contained 3 mg DNA (1 mg coding DNA plasmids, 0.3 mg/L XBP1 expression plasmid, and 1.7 mg sheared herring sperm DNA (Promega, D181A)) combined with 4.5 mL of PEIpro (Sartorius, 101000029). Cells were cultured at 33 °C for 7 days using in-house-developed media.

### Protein Purification

#### PI3Kα

All purification steps were performed at 4 °C unless otherwise stated. Cell pellets from infected insect cultures were lysed using Detergent/Magnetic (DETMAG) method in lysis buffer (20 mM Tris-HCl, pH 8.0, 500 mM NaCl, 10 mM imidazole, 5% (v/v) glycerol, 1 mM TCEP) supplemented with protease inhibitors (Roche protease inhibitor cocktail), benzonase, and IGEPAL CA-630 to a final concentration of 0.2% (v/v). Ni-NTA magnetic beads (28 mL of a 25% slurry, pre-equilibrated in lysis buffer) were added directly to the clarified lysate at a ratio of ∼20 mg protein per 1 mL of settled beads. The lysate–bead mixture was incubated for 1 h at 4 °C with shaking to allow binding of the protein. Beads were collected using a magnetic wand and transferred into Falcon tubes containing wash buffer (20 mM Tris-HCl, pH 8.0, 500 mM NaCl, 10 mM imidazole, 5% (v/v) glycerol, 1 mM TCEP). The resin was washed four times with wash buffer followed by elution using elution buffer (20 mM Tris-HCl, pH 8.0, 500 mM NaCl, 400 mM imidazole, 5% (v/v) glycerol, 1 mM TCEP).

For removal of the N-terminal 6×His tag on p85α, pooled Ni-NTA eluates were incubated with TEV protease and dialyzed overnight against dialysis buffer (20 mM Tris-HCl, pH 8.0, 500 mM NaCl, 5% (v/v) glycerol, 1 mM TCEP). Following cleavage, the sample was subjected to size-exclusion chromatography (SEC) using a Superdex column equilibrated in SEC buffer (25 mM HEPES, pH 7.5, 150 mM NaCl, 5% (v/v) glycerol, 1 mM TCEP).

Prior to cryo-EM, vitrified preparations of purified PI3Kα protein were thawed from –80 °C storage and subjected to SEC to ensure monodispersity. SEC was performed on a Superdex S200 Increase 3.2/300 column (Cytiva) using a MicroÄKTA system equilibrated in SEC buffer (25 mM HEPES, pH 7.5, 150 mM NaCl, 1 mM TCEP). Peak fractions corresponding to the heterodimeric complex were collected and adjusted to a final protein concentration of 1 mg/mL (6 µM). For cryo-EM, the protein was incubated with inavolisib at a 20-fold molar excess (120 µM) for 20 minutes at room temperature, resulting in a final DMSO concentration of 2%.

#### Late embryogenesis abundant protein

All purification steps were performed at 4 °C unless otherwise stated. Cell pellets were resuspended in 50 mM Tris-HCl pH 8.0, 300 mM NaCl, 10 mM imidazole, 1 mM DTT without EDTA and supplemented with protease inhibitor cocktail EDTA-free tablets, 1 mM PMSF, and Benzonase. Cells were lysed on ice by sonication and followed by centrifugation for 1 hr at 40k rpm. The supernatant was incubated with nickel-nitriloacetic acid Ni-NTA agarose beads (Qiagen) for 3 hr with gentle rotation, followed by washing beads in 50 mM Tris-HCl pH 8.0, 300 mM NaCl, 1 mM DTT with 20 mM imidazole. LEA protein was eluted from the beads in 50 mM HEPES pH 7.5, 300 mM NaCl, 250 mM imidazole, 1 mM DTT, 1 mM PMSF. The protein was subsequently concentrated and further purified using HiLoad 16/600 Superdex 75 prep grade column (Cytiva) in buffer containing 50 mM HEPES pH 7.5, 300 mM NaCl, 1 mM TCEP, 10% glycerol. Protein was then concentrated to 1 mM final concentration and snap frozen in liquid nitrogen.

#### μ-opioid-Gi complex

The protein purification was performed on ice or at 4°C unless otherwise noted. The cell pellet from 2 liter of culture (∼40 g) was resuspended at ∼2.5 ml/g of lysis buffer [Buffer LB: 100 mM HEPES-NMDG (N-methyl-d-glucamine) (pH 7.6), 100 mM NaCl, 5 mM MgCl_2_, 10% (w/v) Glycerol 1 µg/ml Benzonase, and ROCHE EDTA-free protease inhibitor tablets]. The cell pellet was lysed by 10-15 passes of dounce homogenization using the 100 ml ‘Type B’ dounce (KIMBLE). The lysed material was then slowly stirred and supplemented with 100 µM TCEP (Tris(2-carboxyethyl)phosphine hydrochloride; Sigma), 10 µg/ml scFv16, 25 mU/ml Apyrase (NEB), and 10 µM DAMGO. The sample was then allowed to incubate at room temperature for two hours. After two hours of incubation, the lysed material was transferred back to 4C and solubilized with 1% (w/v) LMNG (Lauryl Maltose Neopentyl Glycol; Anatrace) and 0.1% (w/v) CHS (Cholesteryl Hemisuccinate Tris Salt; Anatrace) added dropwise from a 10% (w/v) stock prepared in water. The sample was stirred for one hour to extract the membrane proteins. Detergent-solubilized proteins were separated from the insoluble fraction by centrifugation (∼186,000g for 1 hour; Ti45 Rotor at 40,000 rpm). The supernatant after centrifugation was subsequently filtered. 4 ml of Streptactin XT 4Flow resin (IBA) was equilibrated in wash buffer [25 mM HEPES-NMDG pH 7.6, 100 mM NaCl, 25 µM TCEP, 10 µM DAMGO, 5% glycerol, 0.01% LMNG, 0.001% CHS] and then added to the supernatant for 1 hour of batch binding. The supernatant-resin mixture was then transferred to a gravity column and the resin was allowed to pack. The resin was subsequently washed with 10CV of wash buffer. The sample was eluted from the resin using 5 CV of elution buffer [25 mM HEPES-NMDG pH 7.6, 100 mM NaCl, 25 µM TCEP, 10 µM DAMGO, 5% glycerol, 0.01% LMNG, 0.001% CHS, 10 µg/ml scFv16, 50 mM Biotin]. The eluted protein was concentrated using a 100-kDa MWCO concentrator (Amicon) and applied to a Superose 6 Increase 10/300 GL size exclusion column (Cytiva) that was equilibrated in 25 mM HEPES-NMDG pH 7.6, 100 mM NaCl, 10 µM DAMGO, 0.00075% LMNG, 0.000075% CHS, 0.00025% GDN. Peak fractions were evaluated for purity by SDS– polyacrylamide gel electrophoresis and used immediately for cryo-EM studies.

#### ABCC6

The cell pellet was resuspended in a 1:5 (weight:volume) volume of lysis buffer (50 mM HEPES pH 7.5, 300 mM NaCl, 3 mM MgCl_2_, benzonase (1 μg ml^−1^), 1 mM phenylmethylsulfonyl fluoride, and protease inhibitor tablets (Roche). Cells were lysed by Dounce homogenization, and the ABCC6 complex was subsequently solubilized using 1% digitonin (Cayman Chemical) for 2 hours at 4 °C under gentle agitation. The cell lysate was clarified by ultracentrifugation at 125,000g for 45 min, and the supernatant containing the solubilized protein was collected for affinity purification by batch binding to 1 ml of M2-agarose FLAG resin (Sigma-Aldrich) for 1 hour at 4 °C. The protein was then washed with 10 column volumes of GDN wash buffer (50 mM HEPES pH 7.5, 300 mM NaCl, 3 mM MgCl_2_, 0.05% GDN) and eluted in wash buffer supplemented with 3xFLAG peptide (ThermoFisher Scientific). The protein was then concentrated and further purified using Superose 6 3.2/300 (Cytiva) in buffer containing 25 mM HEPES pH 7.5, 150 mM NaCl, 2 mM MgCl_2_, 0.05% GDN. The peak fraction containing ABCC6, as judged by SDS-PAGE gel, was either used directly for preparing cryoKF-containing grids or concentrated to 3 mg/ml to prepare all other grid types.

#### TRIP12-USP7

The cell pellet was resuspended in a 1:5 (weight:volume) volume of lysis buffer (50 mM HEPES pH 7.5, 300 mM NaCl, benzonase (1 μg ml^−1^), 10% glycerol, 1 mM phenylmethylsulfonyl fluoride, and protease inhibitor tablets (Roche). Cells were lysed by sonication on ice at 4 °C. The cell lysate was clarified by ultracentrifugation at 125,000g for 30 min, and the supernatant containing the TRIP12-USP7 complex was collected for affinity purification by batch binding to 5 ml of M2-agarose FLAG resin (Sigma-Aldrich) for 1 hour at 4 °C with gentle shaking. The protein was then washed with 10 column volumes of wash buffer (50 mM HEPES pH 7.5, 300 mM NaCl, 10% glycerol) and eluted in wash buffer supplemented with 3xFLAG peptide (Thermo Fisher Scientific). The protein was then concentrated and further purified using Superose 6 3.2/300 (Cytiva) in buffer containing 25 mM HEPES pH 7.5, 150 mM NaCl, 5% glycerol, 1 mM TCEP.

#### Oncostatin M receptor subunit beta

The filtered supernatant was loaded onto a Nickel Excel column (Cytiva, 20 mL). The column was washed with buffer A (50 mM sodium phosphate, pH 8.0, 200 mM NaCl, 10 mM imidazole), followed by buffer B (buffer A supplemented with 0.1% Triton X-114 and 0.1% Triton X-100). Bound protein was eluted with buffer C (50 mM sodium phosphate, pH 8.0, 200 mM NaCl, 400 mM imidazole). Fractions containing OSMRβ were pooled and further purified by size-exclusion chromatography on a Superdex 200 prep grade column (Cytiva) equilibrated in 20 mM HEPES, pH 7.5, and 150 mM NaCl. Fractions containing pure, monodispersed OSMRβ were pooled, concentrated, flash-frozen in liquid nitrogen, and stored at −80°C until further use.

#### Vixarelimab and scFv16

For Vixarelimab and scFv16, cells were removed by centrifugation at 10,000 × g, and the culture supernatant was filtered through a 0.2 µm membrane. The clarified supernatant was loaded onto Nickel Excel column (Cytiva) equilibrated in 25 mM Tris, pH 7.5, and 300 mM NaCl. The column was washed with the same buffer supplemented with 0.1% Triton X-114 and 0.1% Triton X-100, and bound protein was eluted with 25 mM Tris, pH 7.5, 300 mM NaCl, and 400 mM imidazole. Fractions containing the target of interest, as assessed by SDS-PAGE, were pooled and further purified by size-exclusion chromatography onto a Superdex 200 prep grade column (Cytiva) equilibrated in 20 mM histidine acetate, pH 5.5, and 150 mM NaCl. Fractions containing purified fab or scFv16 were pooled, concentrated, flash-frozen in liquid nitrogen, and stored at −80°C until further use.

For OSMRβ-Vixarelimab complex formation, proteins were combined at 2:1, OSMRβ:vixarelimab molar ratio. The sample was incubated overnight at 4°C and passed over a Superose 6 3.2/300 column (Cytiva) equilibrated in 25 mM HEPES pH 7.5 and 150 mM NaCl. The center peak fraction was used for cryo-EM analysis.

### Grid Preparation

Samples were plunge-vitrified in liquid ethane using a MarkIV Vitrobot (Thermo Fisher Scientific). Grids were blotted for 3-4 s at 7-8 force and the blotting chamber was maintained at 4 °C and 100% humidity during freezing for all samples. Sample concentrations, peptide conditions, and grid types are denoted in Table S4. Peptides were added at 200 µM final concentration, apart from cryoKF, which was added at 100 µM final concentration, directly before grid preparation unless otherwise noted. CryoKF was brought to room temperature and added immediately before freezing with no wait time on the Vitrobot to avoid protein aggregation.

### Cryo-EM Data Collection

Movies were collected across four microscopes and magnification, pixel size, total electron dose, and electron flux for individual datasets are captured in Table S5. Data collected on the Titan Krios G2, Titan Krios G3i, and Glacios2 were collected on Falcon4i detectors using semi-automated imaging in EPU (Thermo Fisher Scientific). Both Titan Krios microscopes are outfitted with a SelectrisX (Thermo Fisher Scientific) energy filter with width set to 10 eV. Data on the Glacios with a K3 direct electron detector operated in counted mode (Gatan) were collected using semi-automated imaging scripts in Smartscope^78^ and SerialEM^79^. A nominal defocus range of −0.8 µm to −2.0 µm was used for data collection, and screening datasets were collected at −1.5 µm to −2.5 µm defocus.

### SPA Data Processing

#### PI3Kα

For each cryoSHIELD condition, single-particle datasets of comparable size were collected to enable systematic comparison. Data was processed in cryoSPARC v4.7.1^73^ using a standard pipeline consisting of 2D classification, heterogeneous refinement, and non-uniform refinement. The statistics for cryo-EM map determination are provided in Table S6.

#### μ-opioid-G_i_ complex

A modified version of the automated GPCR structure determination workflow^83^ was used to process the data. During the live session, movies were upsampled by a factor of 2. To accelerate initial processing steps, motion-corrected micrographs were Fourier-cropped to an effective pixel size of 0.5× the upsampled pixel size. Initial particle picking was performed using templates derived from previous μ-opioid receptor structures. All subsequent processing steps were carried out as described in the automated GPCR workflow. The final particle stack was re-upsampled by a factor of 2 during reference-based motion correction to recover high-resolution information, followed by non-uniform refinement and local refinement. The statistics for cryo-EM map determination are provided in Table S6.

#### ABCC6

A standard data processing workflow was used for new structure determination. CryoSPARC v4.7.1^73^ live was used for automatic motion correction and CTF estimation using standard parameters. Blob picker was used for particle picking, with 360 Å box size and four-times binning. Three rounds of 2D classification using 100 classes, 80 iterations and 400 particles per class was used to obtain well-resolved 2D class averages that showed secondary structural features. Using these particles, three *ab initio* classes were generated using standard parameters. The best resolved *ab initio* class was further refined using non-uniform refinement. This refined map was used as input for two rounds of heterogenous refinement against initial total blob picks against five decoy noise volumes using standard parameters. The particles were then re-extracted without binning and further refined using non-uniform refinement to generate a final consensus ABCC6 map. Masks around nucleotide binding domain 1 and nucleotide binding domain 2 were used for local refinement around the respective regions using pose/gaussian shift prior method. The final composite map was generated using ChimeraX-1.20^77^. The statistics for cryo-EM map determination are provided in Table S6.

#### HUWE1

4,000 movies were processed in cryoSPARC Live by patch motion correction and patch CTF estimation. Micrographs with a CTF fit resolution better than 5 Å were retained. Particles were initially picked with the blob picker and subjected to 2D classification to generate class averages for use as templates in subsequent template picking. Particles were picked with the template picker and filtered with the micrograph junk detector. Selected particles were extracted at 3 Å/pixel and subjected to 2D classification. Classes showing clear particle features were retained. *Ab initio* reconstruction into two classes was performed on all selected particles, and a second *ab initio* reconstruction with two classes was performed on 100 particles to generate ‘decoy’ noise classes for subsequent heterogeneous refinements. Two rounds of heterogeneous refinement were then carried out, with particles from the best class carried forward after each round. Particles from the final best class were re-extracted at 1.2 Å/pixel and refined by non-uniform refinement. Global CTF refinement was performed and a subset of 160,000 particles were refined with non-uniform refinement. All available particles were used when fewer than 160,000 were available (cryoRF condition). Local refinement and ResLog analysis were performed in cryoSPARC.

#### OSMR

For the cryoEV + cryoH dataset, CryoSPARC v5.0.7 live was used for blob picking, with a particle diameter range of 80–120 Å. After export and junk detection, 2D classification was run using a batch size of 250, 170 Å circular mask, and 150 classes, and good 2D class averages were selected. Using these particles, *ab initio* reconstruction was performed with six classes, followed by heterogeneous refinement. Particles from the best two classes were extracted at 1.15 Å/pixel. Using the extracted particles, *ab initio* reconstruction was performed with four classes, followed by heterogeneous refinement, and three good classes were chosen. Non-uniform refinement was performed on 570,298 particles. Particles were sorted through two rounds of subset according to per-particle scale greater >1 and a final non-uniform refinement was performed on 142,234 particles. The global final resolution was 3.37 Å. Local resolution was calculated in cryoSPARC at FSC=0.5.

For the no additive dataset, CryoSPARC v5.0.7 live was used for blob picking, with a particle diameter range of 80–120 Å. After export and junk detection, 2D classification was run using a batch size of 250, 170 Å circular mask, and 150 classes, and good 2D class averages were selected. Using these particles, *ab initio* reconstruction was performed with four classes, followed by heterogeneous refinement. Particles from the best classes were subjected to another round of 2D classification and good 2D class averages were selected. These class averages were used for template picking and particles were extracted at 2.31 Å/pixel. Three rounds of 2D classification using a batch size of 300, a 160 Å circular mask, and 100 classes were run, and any reasonable 2D class averages were selected. Using these particles, *ab initio* reconstruction was performed with six classes, followed by heterogeneous refinement. 8,511 particles from one reasonable class were run through non-uniform refinement. The final resolution was 4.09 Å. Local resolution was estimated in cryoSPARC at FSC=0.5.

#### Other

For any dataset yielding a map for view distribution calculation, data was processed in cryoSPARC live and cryoSPARC v4.7.1 using a standard pipeline consisting of motion and CTF correction, blob and template picking, 2D classification, heterogeneous refinement, and non-uniform refinement. View distributions were plotted using the python script *ordist.py* from the non-uniform refinement output file.

### Model building

AlphaFold2^74^ ABCC6 model was fitted into the cryo-EM density map and then manually rebuilt using Coot 0.9.6^75^ and then refined using Phenix 2.0^76^ real-space refinement using standard parameters against composite ABCC6 cryo-EM map. The statistics for the model are provided in Table S6.

### Data visualization

All molecular model visualization was done using ChimeraX 1.20^77^. The distribution of view plots were visualized using a custom script (https://gist.github.com/alisterburt/7bcdfc902213434ded8c8aa0770744f2).

### cFAR

All Conical Fourier Shell Correlation Area Ratio (cFAR) values presented were calculated using cryoSPARC.

### Particle Density Quantification

For datasets from which maps were reconstructed (Figure 1, Figure 4, Figure S3E), even if anisotropic, the particles included in the final reconstruction were utilized, and the number of particles per micrograph was counted using a manual curation job in cryoSPARC and plotted in a box plot (Microsoft Excel). These datasets were all over 1000 micrographs. Data used for quantitative comparison (Figures 2B, 2D, 2E, and 3B), an UltrAufoil 1.2/1.3, Quantifoil C/Cu 1.2/1.3, and an ANT 1.2/1.3 grids were frozen for each condition and 300 micrographs were collected from 6 different squares on each grid for every condition presented. These micrographs were subjected to patch motion correction, patch CTF estimation, blob picking and 2D classification to generate templates. Template picking and one round of 2D classification were performed, and averages clearly depicting GPCR-G protein complexes or ABCC6 were selected and plotted in a box plot (Microsoft Excel). For screening datasets that did not yield a sufficient number of micrographs or particles for 3D reconstruction (Figures S4C, S4E, S5E, & S5G), two rounds of 2D classification were conducted subsequent to template picking. Any class averages exhibiting particle-like characteristics (micellar shape with extracellular density) were included as picks per micrograph. The number of micrographs analyzed per dataset is specified in the figure legends. Box plots display median, interquartile range, and full data range. All individual data points are shown.

### Tilt-Series Acquisition and Tomogram Reconstruction

Tilt-series were collected on a 300 kV Titan Krios microscope (Thermo Fisher Scientific) equipped with a Falcon 4i (Thermo Fisher Scientific) direct electron detector and a Selectris X (Thermo Fisher Scientific) energy filter. Multi-frame images were acquired using the PACE-tomo script^84^ within SerialEM^79^ at a nominal magnification of 64,000× corresponding to a pixel size of 1.897 Å, with a constant target defocus of −5.0 μm and a 5 eV energy filter slit width. Dose-symmetric tilt-series were collected over an angular range of −54° to +54° in 2° increments, with a total accumulated dose of ∼120 e⁻/Å².

Tilt-series were pre-processed using an internal implementation of WarpTools^85^; https://zenodo.org/records/15749944) for frame-level motion correction and per-tilt CTF estimation. Tilt-series alignment was performed using WarpTools’ integrated Etomo-based^80^ patch-tracking workflow. CTF-corrected tomograms were reconstructed in WarpTools at a pixel size of 10 Å. Tomograms used for visualization were denoised using IsoNet^81^.

### Particle Visualization

Particle positions were determined by template matching with pose optimization, followed by score-based thresholding, using an internal implementation of WarpTools. While the resulting positions of PI3Kα particles are expected to be accurate, their orientations should be considered approximate, as no sub-tomogram averaging was performed to refine angular assignments. For three-dimensional visualization, a low-pass filtered map of PI3Kα was placed back at each template-matched position in ChimeraX^77^ using the ArtiaX plugin^86^. Grayscale tomographic slices were visualized and displayed using IMOD^80^.

### Ice Thickness Estimation

Ice thickness was estimated from tomographic reconstructions of the PI3Kα complex by measuring inter-particle distances along the beam axis. For conditions in which particles were confined to a single monolayer, the inter-particle distance was defined as 0 nm. Where particles partitioned into two distinct layers, the distance between the two layers was measured directly from the tomograms. To convert these inter-particle measurements into an estimate of total ice thickness, we adopted the assumption, consistent with previous reports, that the vitreous ice layer terminates immediately above the particle layer at each interface. Accordingly, we added the longest dimension of the PI3Kα complex (∼14 nm) and a margin of 3× the pixel size (∼3 nm) to each measurement to account for the particle volume excluded from the inter-particle distance and residual estimation error, respectively. This approach provides an estimate of the local ice thickness at both the center and the edge of each hole. Ice thickness distributions were plotted and visualized using Prism11 (GraphPad Software).

### Surface Plasmon Resonance

Surface plasmon resonance (SPR) experiments were performed using a Biacore 1S+ system (Cytiva) in running buffer containing 50 mM HEPES, pH 8.0, 150 mM NaCl, 0.5% PEG-3350, 1% DMSO, and 1 mM DTT. PI3Kα was biotinylated at its N-terminus using in-house-produced sortase A and a biotin-LPXTG reagent by overnight incubation at 4°C. For this, a batch of PI3Kα was cleaved using TEV protease to leave N-terminal glycine for conjugation. Biotinylated PI3Kα was subsequently purified by size-exclusion chromatography in 25 mM HEPES, pH 7.5, and 150 mM NaCl, and immobilized on a Sensor Chip SA (Cytiva, BR100032). Three flow cells were immobilized with PI3Kα at densities ranging from approximately 480 to 1,200 response units (RU). The sensorgrams presented correspond to the highest immobilization density (∼1,200 RU).

Binding measurements were performed using a multi-cycle injection protocol with two-fold serial peptide dilutions starting at maximum concentrations of either 1 or 10 µM. The 10 µM series (0.31–10 µM) is presented, as the lower concentration series did not provide additional interpretable binding responses. Higher peptide concentrations produced substantial nonspecific sensor-surface responses, precluding reliable assessment of protein binding. Sensorgrams were reference-subtracted against a protein-free SA surface. The sensor surface was regenerated with 40% DMSO between cycles. A PDGFR-derived phosphopeptide (740Y-P; Selleckchem, S7865) was used as a positive control. Sensorgrams were analyzed using Biacore Evaluation software, with steady-state affinity analysis performed where appropriate. Experiments were performed in three independent biological replicates with comparable results.

### Thermal Shift Assay

The thermal shift assays was performed using label-free Differential Scanning Fluorimetry (nanoDSF) on a Prometheus NT.48 instrument (NanoTemper Technologies). P3IKα was prepared at final concentrations of 0.4 mg/mL in 25 mM HEPES pH 7.5 and 150 mM NaCl. Where indicated, cryoSHIELD peptides were added to a final concentration of 200 μM. The samples were loaded into standard capillaries and subjected to a linear temperature ramp from 15 °C to 90 °C at a heating rate of 1°C/min. Intrinsic tryptophan and tyrosine fluorescence emission changes were recorded at 350 nm and 330 nm.

For data analysis, the fluorescence intensity ratio (F350/F330) was calculated, and its first derivative with respect to temperature was determined and baseline-corrected. The resulting derivative curves were integrated and normalized to a 0–100% scale. For the inavolisib-treated control, the first derivative exhibited an inverted transition; therefore, its sign was reversed before integration to allow consistent fitting of the unfolding transitions across conditions. Apparent melting temperatures (Tₘ) were determined from the transition midpoint (V50) of a Boltzmann sigmoidal nonlinear regression fitted to the normalized curves using GraphPad Prism. Experiments were performed in duplicate using independently prepared biological samples, and Tₘ values are reported as mean ± standard deviation (SD).

### Circular dichroism

Far-UV circular dichroism (CD) measurements were performed using a Jasco J-815 CD spectropolarimeter equipped with a 1 mm path-length quartz cuvette at room temperature. CryoSHIELD peptides were diluted from 2 mM stock solutions to a final concentration of 0.3 mg/mL in filtered phosphate-buffered saline (PBS). Spectra were collected over a wavelength range of 200-300 nm, scan speed of 100 nm/min, and data interval of 1 nm. Three consecutive scans were acquired for each sample, averaged, and corrected by subtraction of a corresponding buffer spectrum collected under identical conditions. The resulting spectra were expressed as raw ellipticity (mdeg) and plotted to assess peptide secondary structure in solution. Measurements were repeated using three independently prepared samples.

---

**Figure S1.**
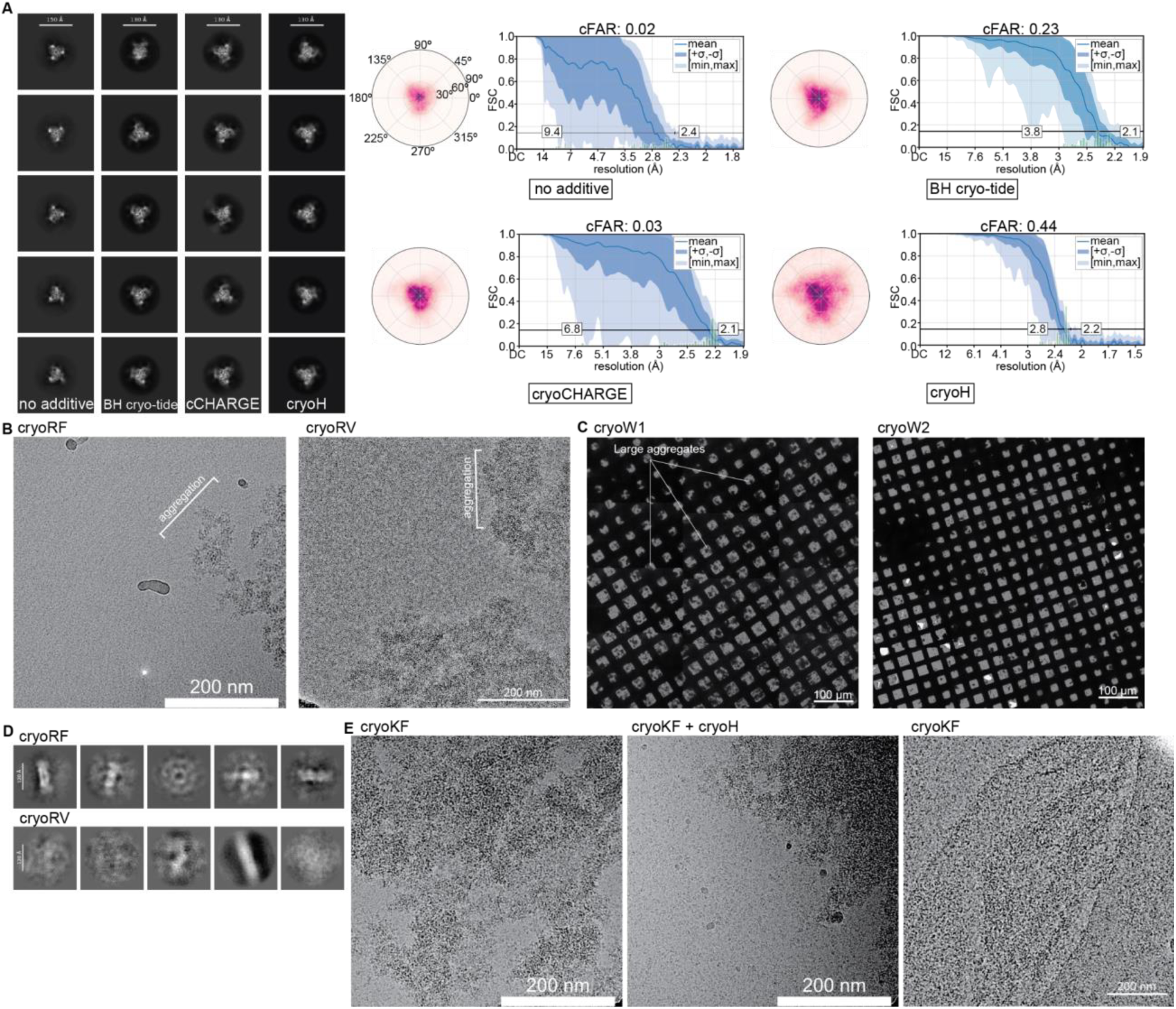
Negative data from cryoSHIELD designs. (A) Representative 2D averages of a homotrimeric immune ligand bound to receptor extracellular domains (ECDs) along with the viewing direction distribution and cFAR plots for their 3D reconstructions with C3 symmetry. Scale bars in 2D averages are indicated in each panel. (B) Example micrographs of Arg-containing cryoSHIELDs that have caused aggregation for soluble proteins, a nucleosome-binding protein, and a soluble enzyme, respectively. Scale bars, 200 nm. (C) Grid atlases showing micron-sized aggregation of the TRIP12-USP7 complex and BCDX2-CX3-RAD51 complex with Trp-containing cryoSHIELDs. Scale bar, 100 µm. (D) Representative 2D averages from “flat” enzyme samples (Figure 1B) that aggregated when mixed with RV and RF. (E) Example micrographs of cryoKF aggregating 70 kDa protein tyrosine phosphatase and a Fab + immune receptor complex (Figure 1A) when applied alone or mixed with cryoH. Scale bars, 200 nm. Related to Table 1.

**Figure S2.**
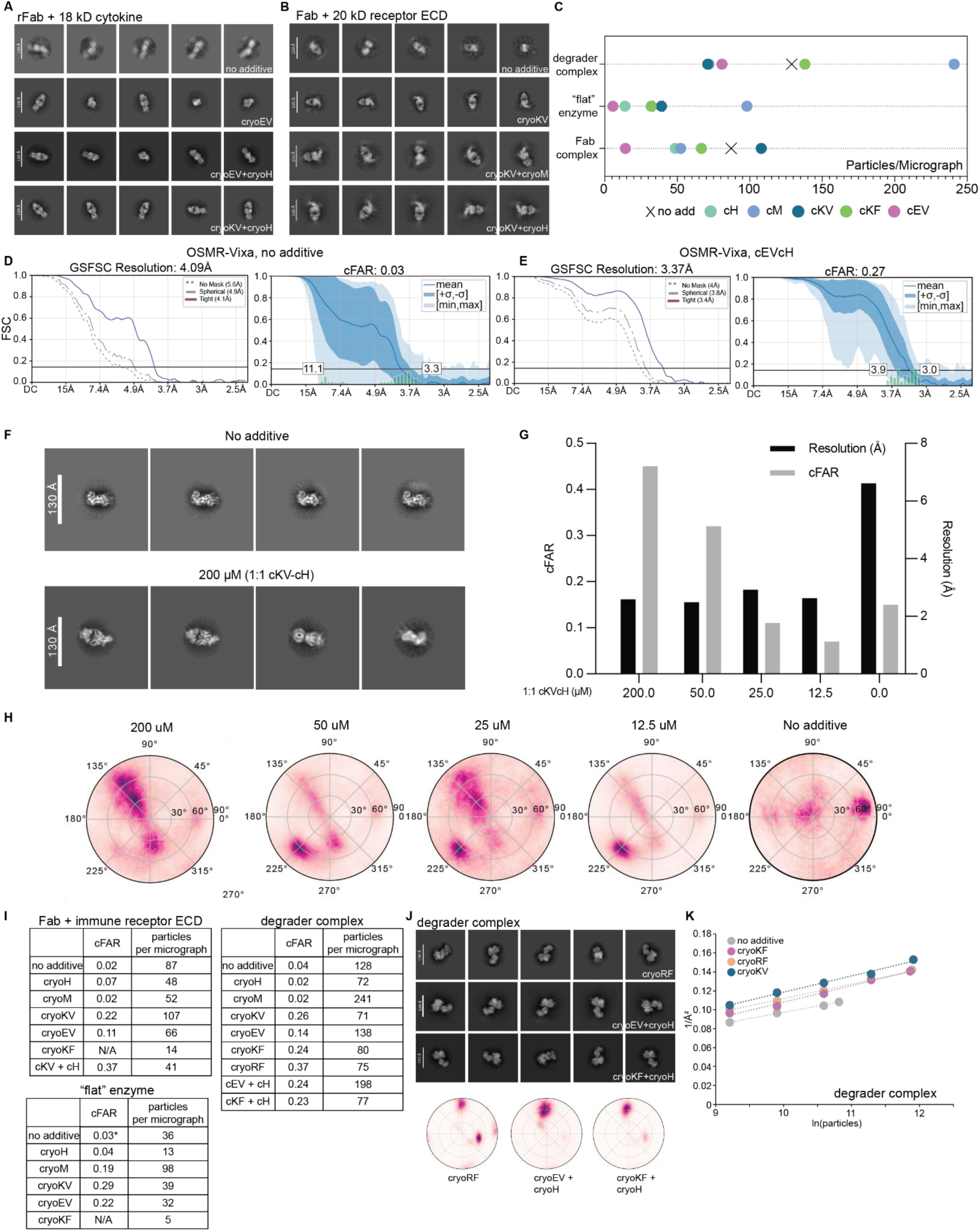
CryoSHIELDs address orientation bias across samples. (A and B) Representative 2D class averages of Fab complexes bound to an 18 kDa cytokine (A) or a 20 kDa receptor extracellular domain (B), prepared without additives, with individual linear cryoSHIELDs, or with linear– helical peptide combinations, as indicated. Combining linear and helical peptides introduced additional particle orientations beyond those obtained with individual peptides. (C) Plot of the particle pick counts for Figure 1A-C cryo-EM cases for different types of cryoSHIELDs. (D, E) Comparison of OSMRβ–Fab cryo-EM reconstruction statistics obtained without additives (D) or with cryoEV–cryoH (E). Gold-standard FSC (GSFSC) and cFAR curves are displayed. (F) Representative 2D class averages of the Fab–antigen complex prepared without additive or in the presence of 200 µM 1:1 cryoKV-cryoH (cKV–cH). Scale bars, 130 Å. (G) cFAR (gray) and map resolution (black) plots of Fab-antigen complex in (F) obtained following grid preparation with decreasing concentrations of 1:1 cryoKV-cryoH mix (total peptide concentration: 200, 50, 25, 12.5, and 0 µM). (H) Angular distribution plots of particles from the corresponding cryo-EM datasets in (G). (I) Summary of cFAR values and particles per micrograph for the Fab–immune receptor ECD complex, a 640 kDa soluble octameric enzyme with a flat architecture, and an E3 ubiquitin ligase–degrader complex under the indicated cryoSHIELD conditions. Values were obtained from representative cryo-EM datasets. N/A indicates conditions for which reliable cFAR estimation was not possible. *-The no-additive flat-enzyme reconstruction was obtained using tilted data collection. (J) Representative 2D class averages and corresponding angular distribution plots of an E3 ubiquitin ligase–degrader complex prepared with cryoRF, cryoEV–cryoH, or cryoKF–cryoH, illustrating peptide-dependent changes in particle orientation. (K) Rosenthal–Henderson analysis of the degrader complex reconstructed without additives or with cryoKF, cryoRF, or cryoKV. Inverse-squared resolution (1/Å²) is plotted against the natural logarithm of the number of particles used for reconstruction, with dotted lines representing linear fits. Related to Figure 1.

**Figure S3.**
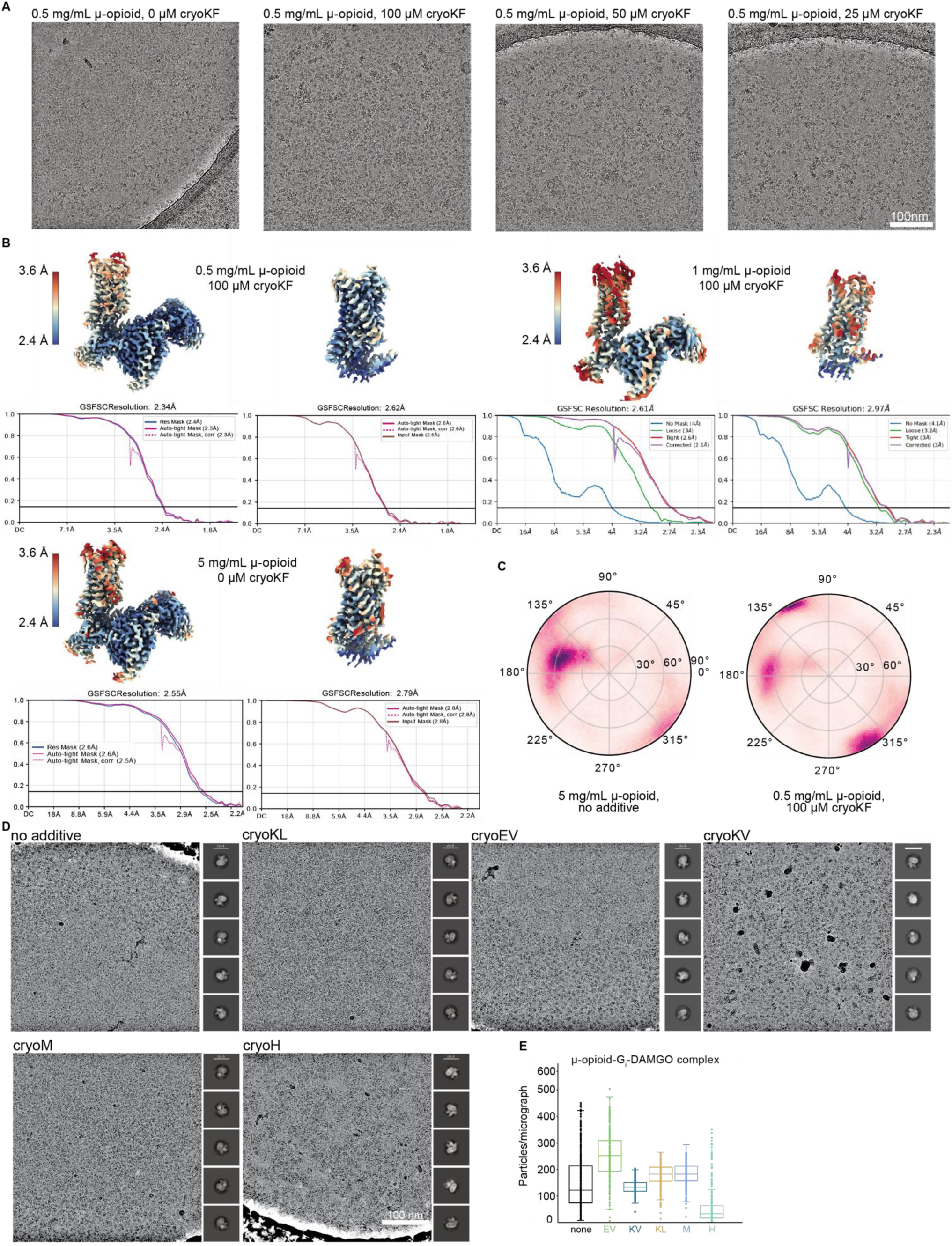
CryoKF enables high-resolution reconstruction of the μ-opioid receptor–Gi complex at reduced protein concentrations. (A) Representative cryo-EM micrographs of the μ-opioid receptor–Gi complex prepared with increasing concentrations of cryoKF (0, 25, 50, and 100 µM), illustrating peptide-dependent changes in particle density. (B) Cryo-EM reconstructions of the μ-opioid receptor–Gi complex prepared at 0.5 or 1 mg/mL with 100 µM cryoKF, or at 5 mg/mL without additives. Global reconstructions and corresponding focused receptor maps are colored by local resolution, with gold-standard Fourier shell correlation (FSC) curves shown below. The 0.5 mg/mL cryoKF condition yielded global and focused receptor resolutions of 2.34 Å and 2.59 Å, respectively, compared with 2.61 Å and 2.97 Å at 1 mg/mL with cryoKF, and 2.55 Å and 2.79 Å at 5 mg/mL without additives. FSC curves obtained with different masking procedures are shown as indicated. (C) Angular distribution plots of the μ-opioid receptor–Gi complex reconstructed at 5 mg/mL without additives or at 0.5 mg/mL with 100 µM cryoKF, illustrating changes in orientations between the two conditions. (D) Representative cryo-EM micrographs and corresponding 2D class averages of μ-opioid receptor–Gi complexes prepared without additives or with cryoKL, cryoEV, cryoKV, cryoM, or cryoH. Scale bar, 100 nm. (E) Quantification of particle density (particles per micrograph) for the μ-opioid receptor–Gi–DAMGO complex under the peptide conditions shown in (D). Box plots summarize the distribution of particle counts across analyzed micrographs, counted from 3D reconstructions of 1000 movie datasets, illustrating peptide-dependent differences in particle enrichment. Related to Figure 2.

**Figure S4.**
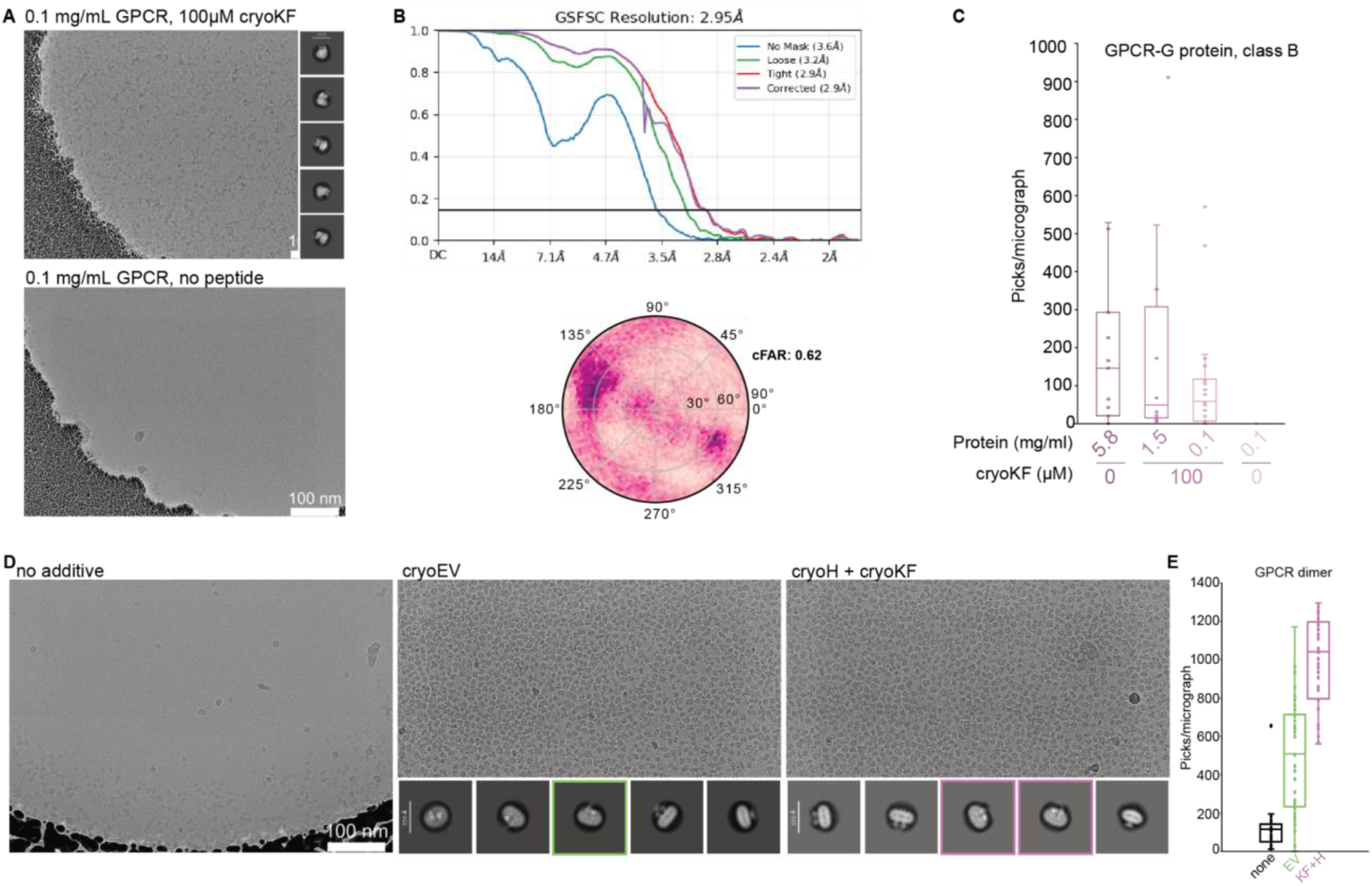
CryoSHIELD effects on other GPCRs. (A) Representative cryo-EM micrographs and 2D class averages of a class B GPCR–G protein complex prepared at 0.1 mg/mL in the presence or absence of 100 µM cryoKF. Scale bar, 100 nm. Scale bars in 2D averages are indicated in each panel. (B) Gold-standard Fourier shell correlation (FSC) curves and angular distribution plot for the class B GPCR–G protein complex reconstructed from the 0.1 mg/mL cryoKF-treated sample, yielding a global resolution of 2.95 Å and a conical FSC area ratio (cFAR) of 0.62. (C) Quantification of particle density for the class B GPCR–G protein complex prepared at different protein concentrations (5.8, 1.5, and 0.1 mg/mL), with or without 100 µM cryoKF, as indicated. 900 micrographs, across 3 grid types, were analyzed for each condition. (D) Representative cryo-EM micrographs and 2D class averages of a GPCR dimer prepared without additives, with cryoEV, or with a combination of cryoH and cryoKF. Both peptide-containing conditions increased particle density, with the cryoH–cryoKF combination producing particularly pronounced enrichment and unique views (pink boxes). Scale bars, 100 nm. Scale bars in 2D averages are indicated in each panel. (E) Quantification of particles per micrograph for the GPCR dimer under the conditions shown in (D). Box plots illustrate the distributions of particle counts, demonstrating substantial enrichment with cryoEV and further enrichment with the cryoH–cryoKF combination. Related to Figure 2.

**Figure S5.**
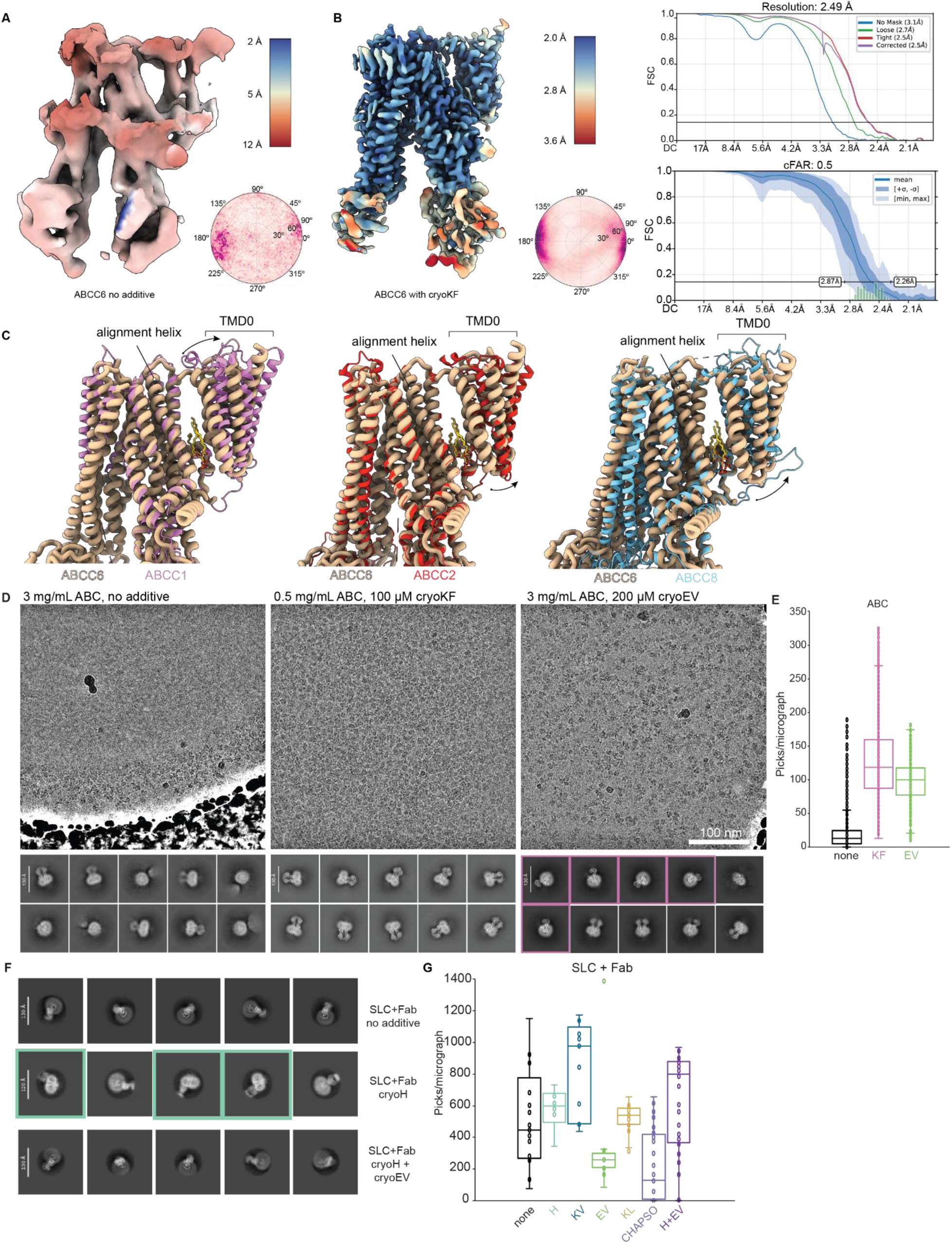
CryoSHIELDs increase particles for low-yield membrane proteins and enable structural determination of nanodisc samples by inducing side views. (A) Consensus cryo-EM reconstruction of ABCC6 colored by local resolution displayed together with the angular distribution of views. (B) Composite cryo-EM reconstruction of ABCC6 colored by local resolution displayed together with the angular distribution of views, the FSC, and the cFAR graphs. (C) Molecular model comparison between ABCC6 and ABCC1 (left), ABCC2 (center) and ABCC8 (right), with focus on TMD0. All structures were aligned on TMD1 helices adjacent to the TMD0 domain. (D) Representative micrographs and 2D averages of a low-yield ABC transporter vitrified with no additive, 100 µM cryoKF, and 200 µM cryoEV. No additive and cryoEV samples were vitrified at 3 mg/mL and the 100µM cryoKF sample was vitrified at 0.5 mg/mL. Scale bar, 100 nm. Scale bars in 2D averages are indicated in each panel. (E) Data corresponds to ABC transporter from S5D and picks per micrograph across conditions are counted and displayed. Picks were counted from particle-like 2D averages after two rounds of sorting. A 3D reconstruction could not be generated from the no additive sample for comparison of true particles. (F) 2D averages for a Fab-bound SLC transporter vitrified with different cryoSHIELDs at 200µM additive. Scale bars in 2D averages are indicated in each panel. (G) Nominal particles per micrograph count for the Fab-bound SLC transporter in panel F vitrified with different cryoSHIELDs at 200µM. Due to limited dataset size, a 3D reconstruction could not be generated in all cases so particle counts were derived from particle-like 2D class averages after two rounds of sorting. Related to Figure 3.

**Figure S6.**
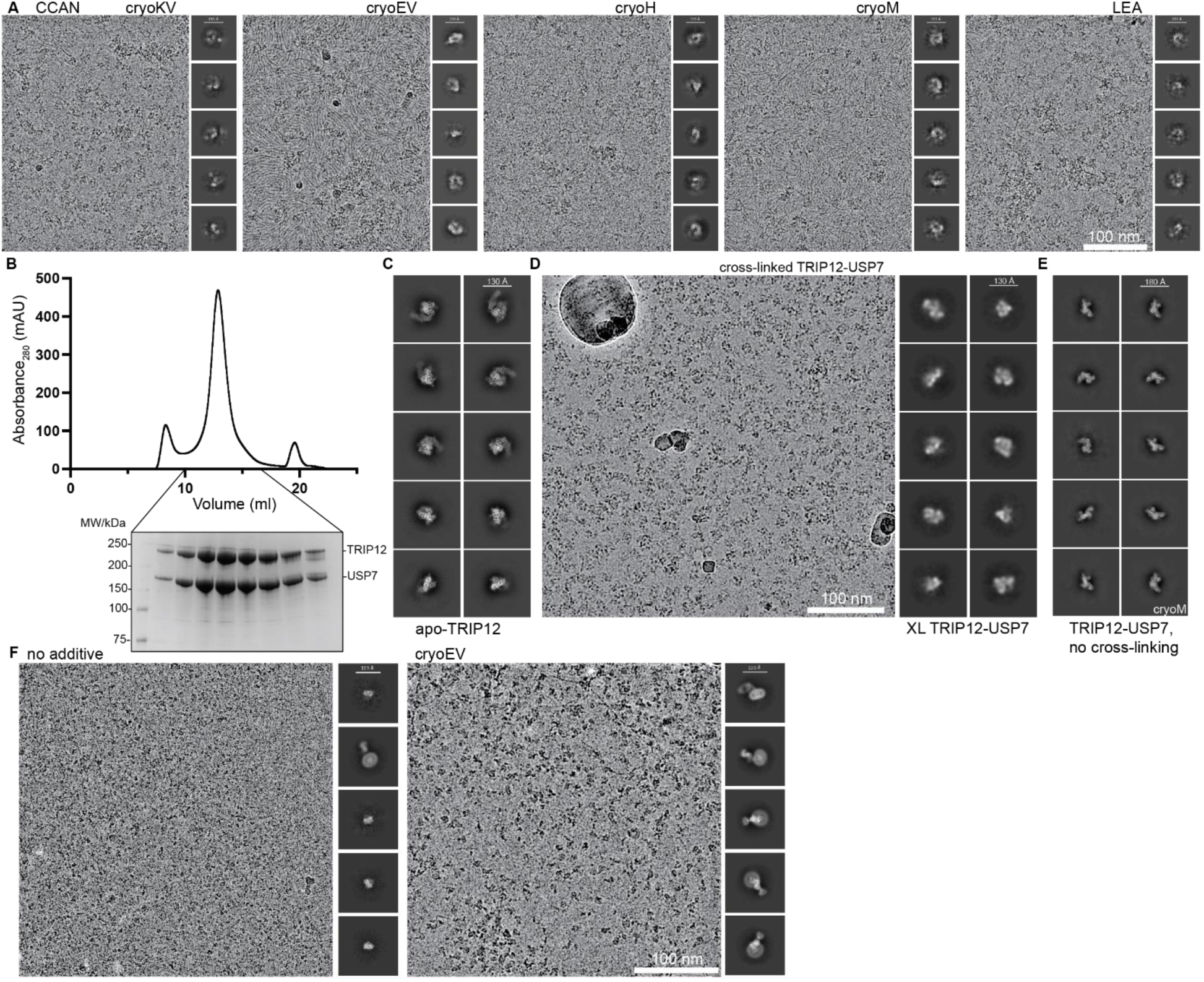
CryoSHIELDs can stabilize complexes against AWI. (A) Representative micrograph and uninterpretable 2D averages from single cryoSHIELD-treated samples for the CCAN complex. Scale bar, 100 nm. Scale bars in 2D averages are indicated in each panel. (B) Size exclusion chromatography and SDS-PAGE gel indicating complex formation between TRIP12 and USP7. (C) 2D class averages of TRIP12 in the absence of USP7. Scale bars in 2D averages are indicated in each panel. (D) Representative micrograph and uninterpretable 2D class averages of a TRIP12-USP7 sample that was crosslinked before grid preparation. Scale bar, 100 nm. Scale bars in 2D averages are indicated in each panel. (E) 2D class averages of unmodified TRIP12–USP7 prepared with cryoM, showing a unique view of the TRIP12 vitrified with cryoM. Scale bars in 2D averages are indicated in each panel. (F) Representative micrographs and 2D class averages of ɑ-BRIL Fab-BRIL-Class B GPCR fusion complex with and without cryoEV. Primarily Fabs are present in the no additive sample, while intact complexes make up the majority of the particles in the cryoEV sample. Scale bar, 100 nm. Scale bars in 2D averages are indicated in each panel. Related to Figure 4.

**Figure S7.**
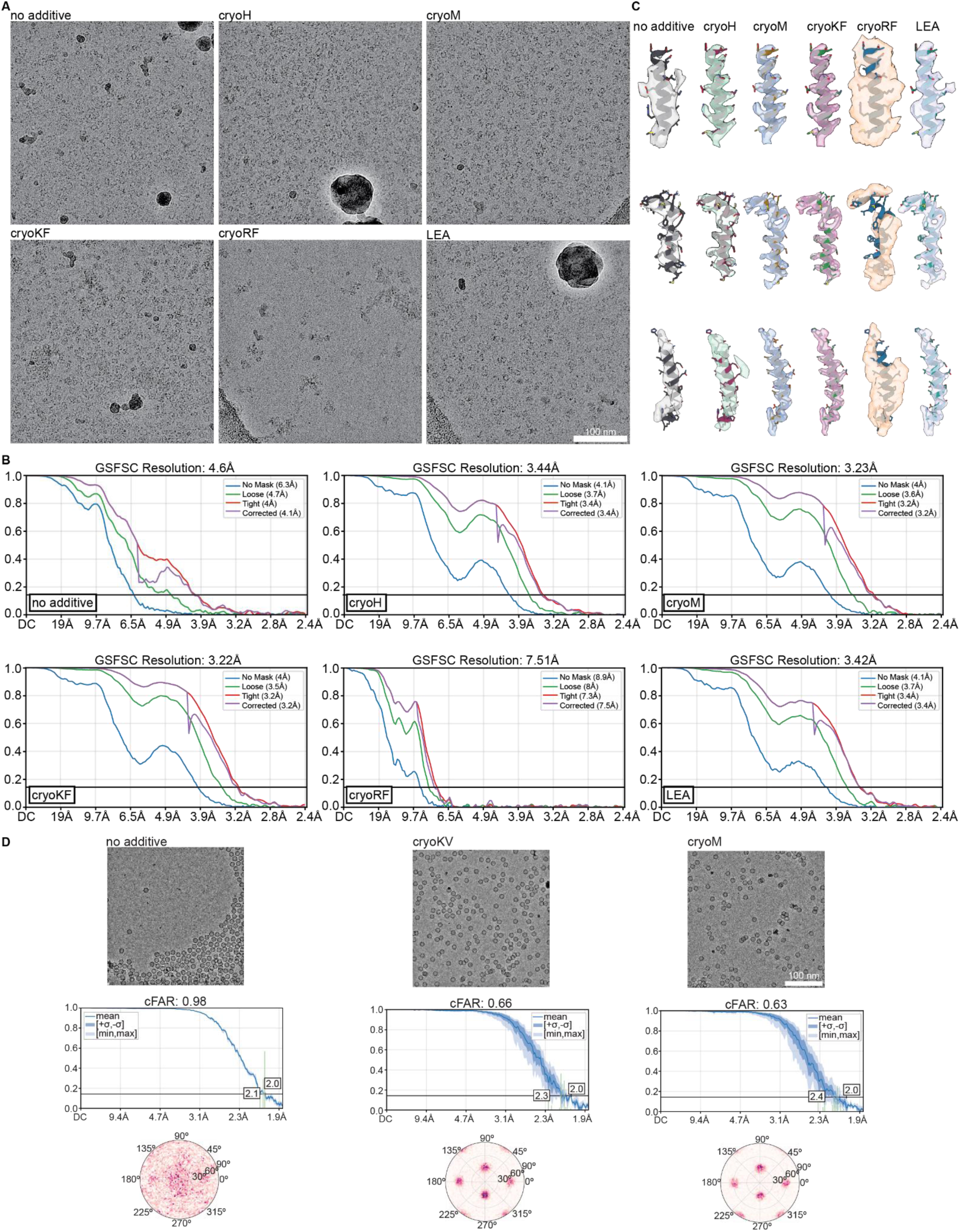
CryoSHIELDs effect on HUWE1 and AAVs. (A) Representative cryo-EM micrographs of HUWE1 prepared without additives or in the presence of cryoH, cryoM, cryoKF, cryoRF, or LEA protein. Micrographs were selected at comparable defocus values to illustrate differences in particle distribution and image contrast. Scale bar, 100 nm. (B) Gold-standard Fourier shell correlation (FSC) curves for HUWE1 reconstructions obtained from equivalent datasets under the conditions shown in (A). FSC curves calculated using different masking and correction procedures are shown as indicated. Global resolution estimates range from 3.22–3.44 Å for cryoH, cryoM, cryoKF, and LEA, compared with 4.6 Å without additives and 7.51 Å with cryoRF. (C) Representative atomic model-to-map fits illustrating local cryo-EM density quality for HUWE1 reconstructed under the indicated conditions. Corresponding structural regions are displayed across datasets to enable direct comparison of map interpretability and the resolution of secondary-structure and side-chain features. (D) Representative cryo-EM micrographs, conical FSC area ratio (cFAR) plots, and angular distribution maps for adeno-associated virus (AAV) particles prepared without additives or with cryoKV or cryoM. Related to Figure 4.

**Figure S8.**
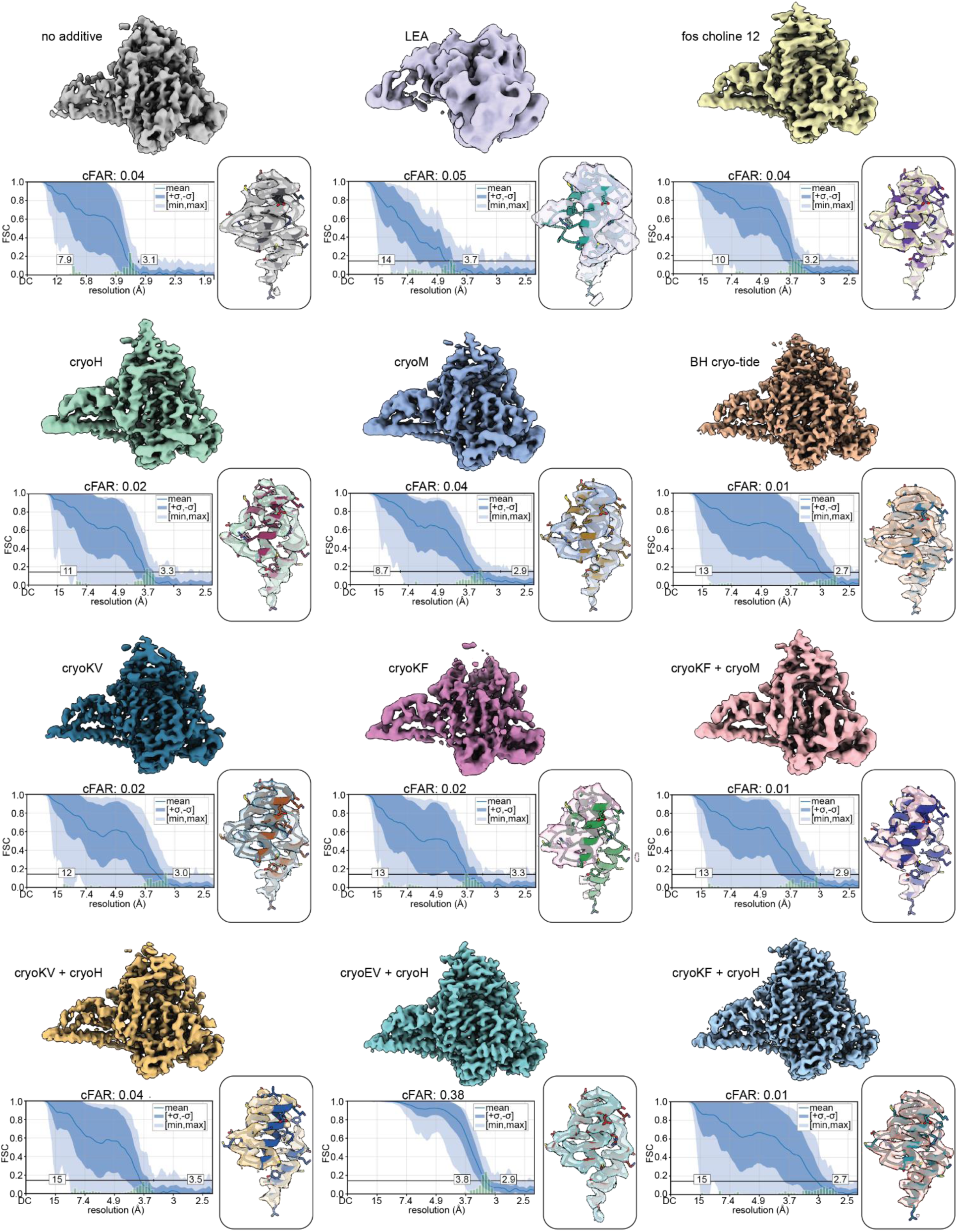
Effects of cryoSHIELDs on PI3Kα angular sampling and cryo-EM map quality. Comparison of PI3Kα cryo-EM reconstructions obtained without additives or in the presence of LEA protein, Fos-choline-12, BH cryo-tide, individual cryoSHIELDs (cryoH, cryoM, cryoKV, and cryoKF), or combinations of linear and helical peptides (cryoKF–cryoM, cryoKV–cryoH, cryoEV–cryoH, and cryoKF–cryoH), as indicated. For each condition, the overall cryo-EM reconstruction (top), directional Fourier shell correlation (FSC) analysis (bottom left), and representative atomic model-to-map fit (bottom right) are shown. Directional FSC plots display the mean FSC (solid blue line), ± standard deviation (dark blue shading), and minimum-to-maximum range (light blue shading), with the corresponding conical FSC area ratio (cFAR) indicated above each plot. Related to Figure 5

**Figure S9.**
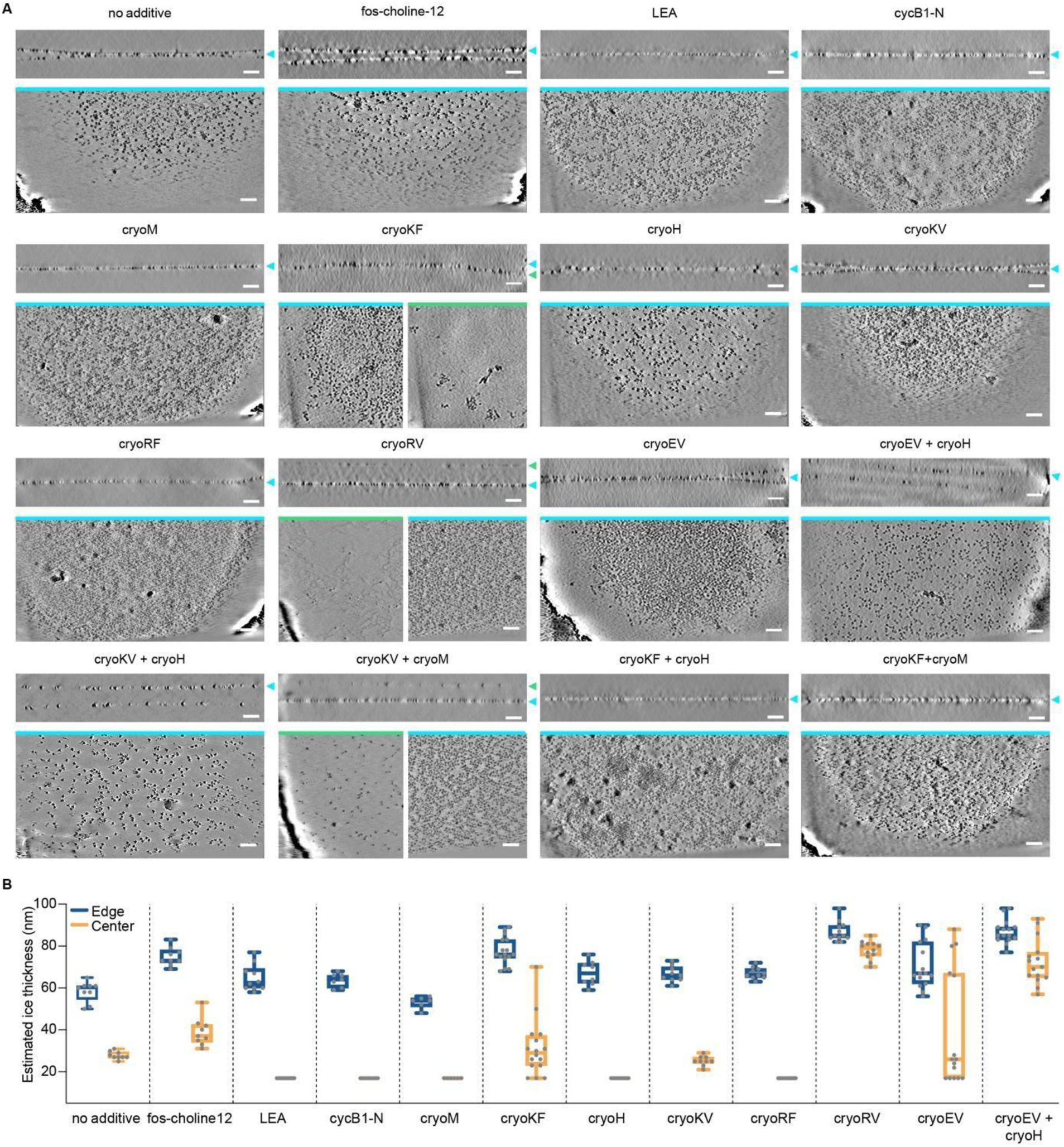
PI3Kα particle distribution. (A) Denoised tomographic views corresponding to the schematics shown in Figure 5, along with additional views for peptide combinations not presented in the main figure. For each condition, the top panel shows a cross-sectional (side-view) tomographic slice through the vitreous ice layer. Below, one or more planar (in-plane) tomographic slices are displayed, extracted at the heights indicated by color-coded arrows in the corresponding cross-section. The primary particle layer is shown for all conditions (cyan); where distinct additional layers were observed, arising from particle partitioning, aggregation, or cryoSHIELD-related artifacts, these are presented as separate planar slices (green). Border line colors of the in-plane views match the corresponding arrow colors in the cross-section. Scale bars, 50 nm. (B) Estimated ice thickness at the edge (blue) and center (orange) of the foil hole for each additive condition, calculated as described in the Methods. Box plots display median, interquartile range, and full data range. Number of measurements per condition: No Additive, *n* = 9; Fos-choline-12, *n* = 9; LEA, *n* = 10; BH cryo-tide, *n* = 7; cM, *n* = 6; cKF, *n* = 16; cH, *n* = 8; cKV, *n* = 9; cRF, *n* = 10; cRV, *n* = 12; cEV, *n* = 15; cEV+cH, *n* = 18. All individual data points are shown. Related to Figure 5.

**Figure S10.**
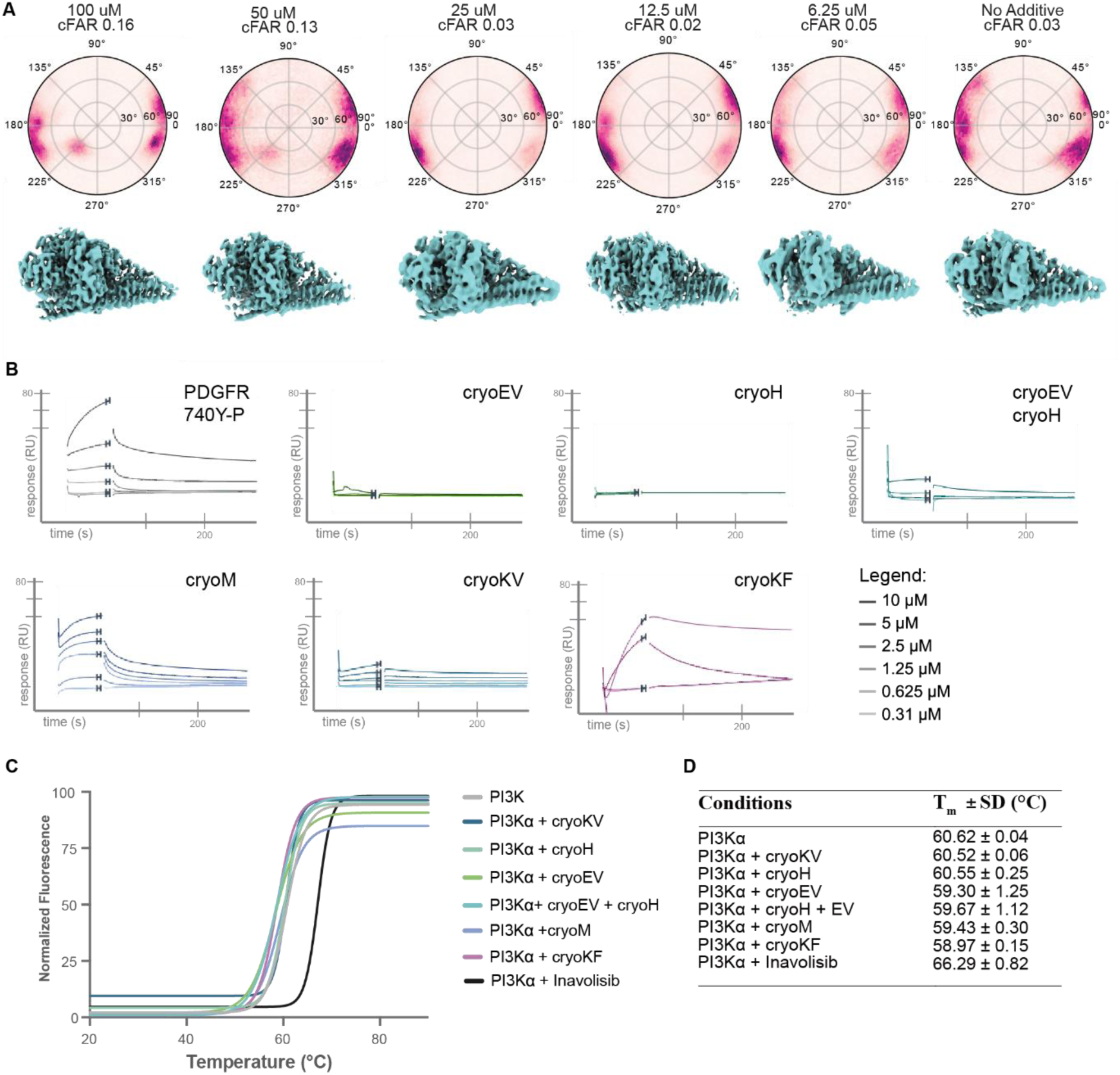
Effect of cryoSHIELDs on PI3Kα in solution. (A) Concentration-dependent effects of cryoEV–cryoH on PI3Kα particle orientation and cryo-EM reconstruction. Angular distribution plots and corresponding cryo-EM density maps for PI3Kα prepared in the presence of decreasing concentrations of cryoEV and cryoH (100, 50, 25, 12.5, and 6.25 µM each) or without additive. The corresponding cFAR values are indicated above each angular distribution plot. (B) Representative SPR sensorgrams examining cryoSHIELD peptides interactions with immobilized PI3Kα. A PDGFR-derived phosphopeptide (740Y-P) was used as a positive control. Peptides were injected as two-fold serial dilutions ranging from 0.31 to 10 µM. Most cryoSHIELDs showed no appreciable concentration-dependent binding, whereas cryoM exhibited detectable association and cryoKF produced a weaker, noncanonical apparent response. Representative sensorgrams from one of three independent experiments are shown (C) NanoDSF thermal unfolding profiles of PI3Kα in the absence or presence of individual cryoSHIELDs, a cryoEV–cryoH mixture (200 µM total peptide concentration), or the PI3Kα inhibitor inavolisib as a positive control. Curves represent the normalized integral of the baseline-corrected first derivative of the fluorescence ratio (F350/F330), fitted using sigmoidal Boltzmann nonlinear regression. (D) Apparent melting temperatures (Tₘ) of PI3Kα under the conditions shown in (C), calculated from the midpoint (V50) of the fitted thermal unfolding transitions. Values represent mean ± SD from two biologically independent replicate measurements. Related to Figure 5.

**Figure S11.**
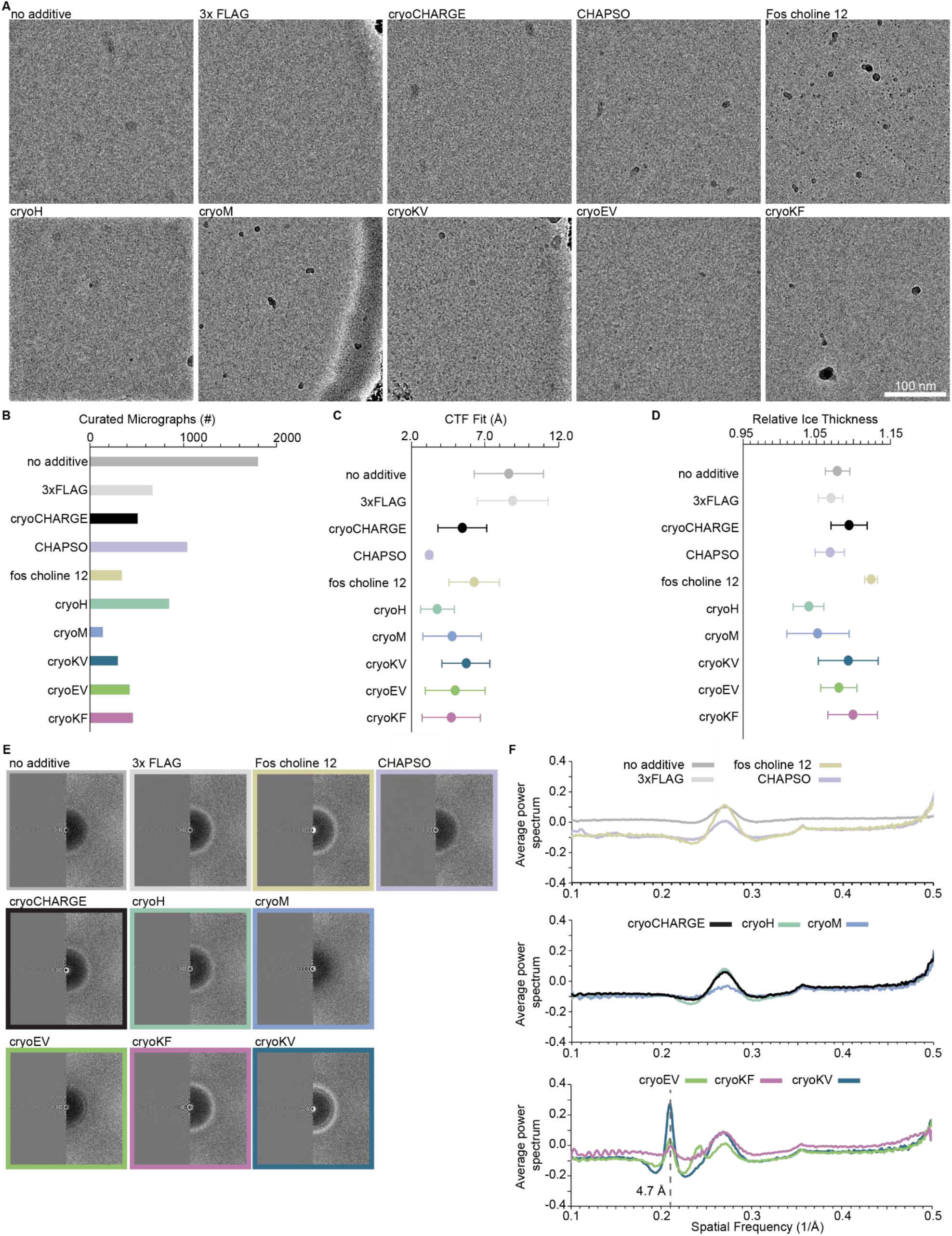
CryoSHIELDs do not substantially interfere with cryo-EM imaging and linear peptides exhibit characteristic diffraction features. (A) Representative cryo-EM micrographs of grids prepared with buffer alone (no additive), control peptides (3×FLAG and cryoCHARGE), conventional detergents (CHAPSO and fos-choline-12), or individual cryoSHIELDs, in the absence of target proteins. (B) Number of curated micrographs analyzed for each condition. Number of micrographs per condition: PBS, 1803; 3xFLAG, 671; cCHARGE, 510; CHAPSO, 1037; fos choline 12, 847; cH, 847; cM, 142; cKV, 300; cEV, 425; cKF, 459. (C) Comparison of contrast transfer function (CTF) fit resolution across the indicated conditions. Mean ± SD. (D) Relative ice-thickness estimates derived from the analyzed micrographs for each condition. Mean ± SD. (E) Representative two-dimensional fast Fourier transforms (FFTs) of micrographs collected under the indicated conditions. (F) Rotationally averaged one-dimensional power spectra corresponding to the conditions in (E), plotted against spatial frequency. Linear cryoSHIELDs (cryoEV, cryoKF, and cryoKV) exhibit a characteristic diffraction peak at approximately 4.7 Å, absent from helical peptides, control peptides, and conventional detergents. This periodicity is consistent with ordered β-sheet-like peptide assemblies. Importantly, this diffraction feature does not prevent CTF estimation, as demonstrated in (C). Related to Figure 6.

**Figure S12.**
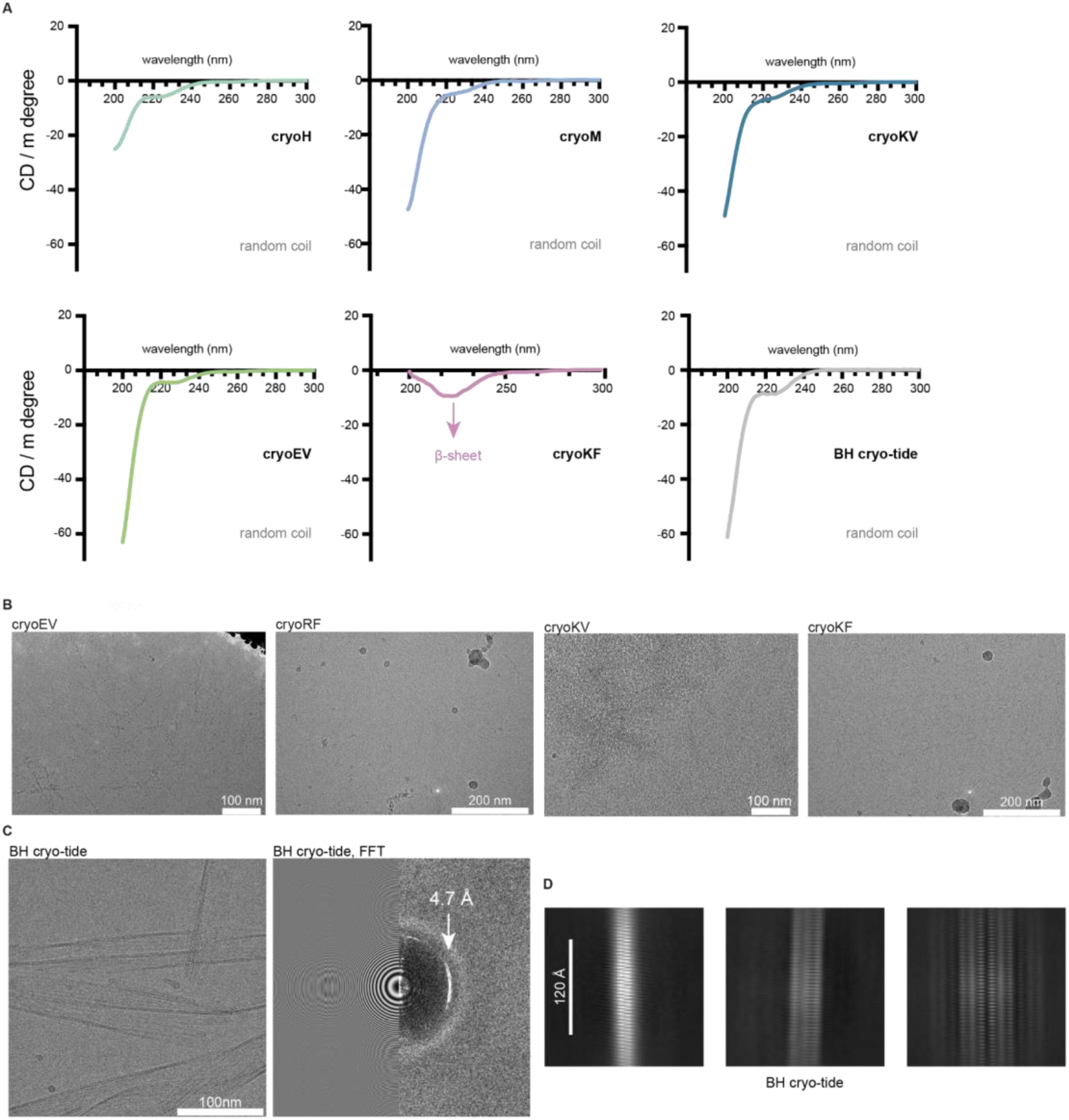
Secondary structure of cryoSHIELDs in solution and higher-order peptide assemblies observed during cryo-EM grid preparation. (A) Far-UV circular dichroism (CD) spectra of cryoSHIELD peptides (cryoH, cryoM, cryoKV, cryoEV, and cryoKF) and the BH cryo-tide control peptide. Peptides were analyzed at 0.3 mg/mL in PBS at room temperature. Spectra represent the average of three consecutive scans following buffer subtraction. Most peptides exhibited spectral profiles consistent with predominantly disordered conformations in solution, whereas cryoKF showed a weak negative band characteristic of β-sheet propensity. (B) Representative cryo-EM micrographs illustrating occasional higher-order peptide assemblies and filamentous or sheet-like artifacts observed with linear cryoSHIELDs (cryoEV, cryoRF, cryoKV, and cryoKF). These artifacts were particularly associated with prolonged peptide stock storage or extensive agitation. Scale bars, 100 or 200 nm, as indicated. (C) Representative cryo-EM micrograph of extended filamentous assemblies formed by BH cryo-tide (left) and the corresponding fast Fourier transform (FFT; right). The prominent diffraction feature at approximately 4.7 Å is highlighted. Scale bar, 100 nm. (D) Representative 2D class averages of BH cryo-tide filaments showing distinct periodic striations and variations in filament organization, consistent with the formation of ordered peptide assemblies with different packing arrangements. Scale bar, 120 Å. Related to Figure 6.

**Figure S13.**
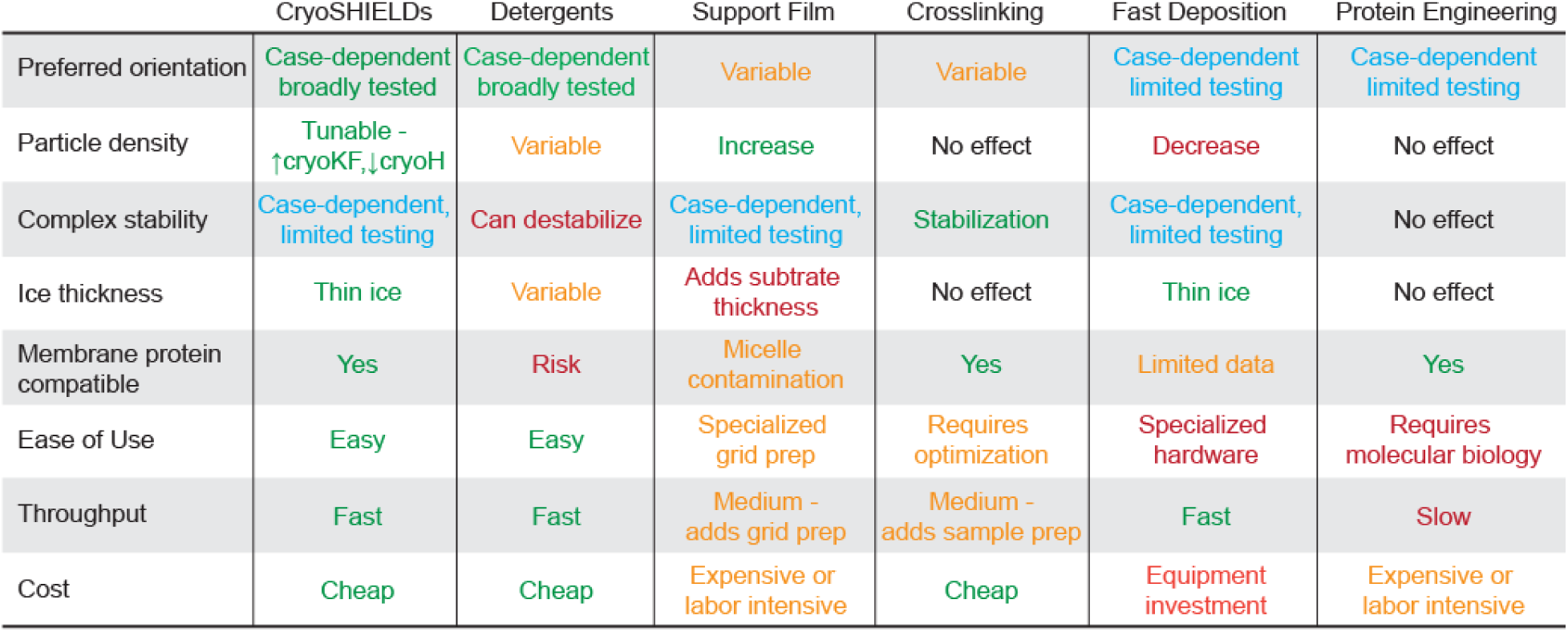
Comparing cryoSHIELDs to field standards for combatting the air-water interface. Table comparing and contrasting different techniques used to combat problems caused by the AWI. Color coding ranks from particularly good or easy (green) -> blue -> orange -> to particularly bad or difficult (red). Related to Figure 6.

**Table S1.**
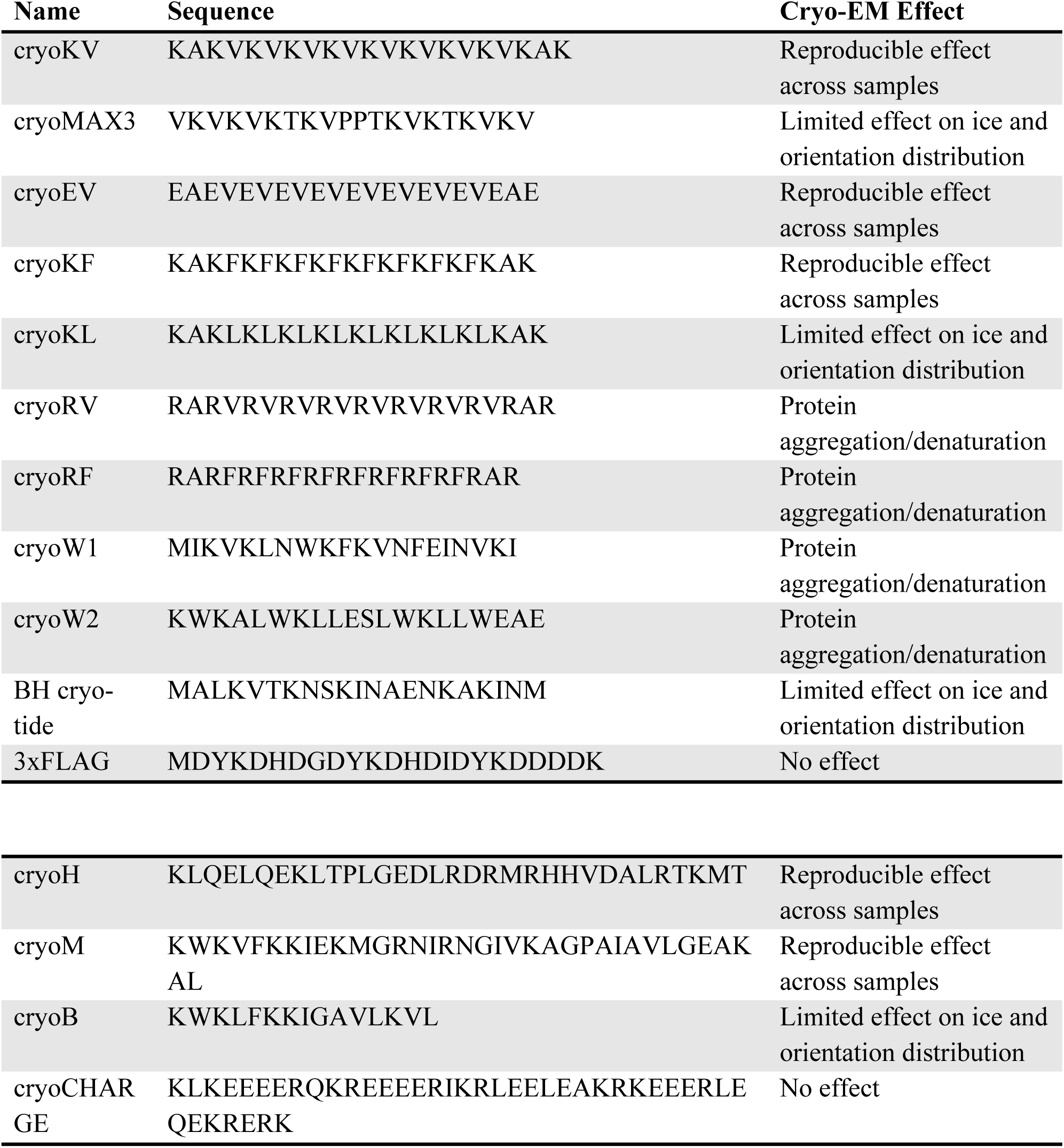
Peptide sequences used in this study. Related to Table 1.

| **Name** | **Sequence** | **Cryo-EM Effect** |
|---|---|---|
| cryoKV | KAKVKVKVKVKVKVKVKVKAK | Reproducible effect across samples |
| cryoMAX3 | VKVKVKTKVPPTKVKTKVKV | Limited effect on ice and orientation distribution |
| cryoEV | EAEVEVEVEVEVEVEVEVEAE | Reproducible effect across samples |
| cryoKF | KAKFKFKFKFKFKFKFKAK | Reproducible effect across samples |
| cryoKL | KAKLKLKLKLKLKLKLKLKAK | Limited effect on ice and orientation distribution |
| cryoRV | RARVRVRVRVRVRVRVRVRAR | Protein aggregation/denaturation |
| cryoRF | RARFRFRFRFRFRFRFRAR | Protein aggregation/denaturation |
| cryoW1 | MIKVKLNWKFKVNFEINVKI | Protein aggregation/denaturation |
| cryoW2 | KWKALWKLLESLWKLLWEAE | Protein aggregation/denaturation |
| BH cryo-tide | MALKVTKNSKINAENKAKINM | Limited effect on ice and orientation distribution |
| 3xFLAG | MDYKDHDGDYKDHDIDYKDDDDK | No effect |
| cryoH | KLQELQEKLTPLGEDLRDRMRHHVDALRTKMT | Reproducible effect across samples |
| cryoM | KWKVFKKIEKMGRNIRNGIVKAGPAIAVLGEAKAL | Reproducible effect across samples |
| cryoB | KWKLFKKIGAVLKVL | Limited effect on ice and orientation distribution |
| cryoCHARGE | KLKEEEERQKREEEERIKRLEELEAKRKEEERLEQEKRERK | No effect |

**Table S2.** Marangoni assay results for peptides and common detergents. Related to Table 1.

| **Name** | **Marangoni CSC (µM)** |
|---|---|
| cryoKV | 12.5 |
| cryoMAX3 | 25 |
| cryoEV | 12.5 |
| cryoKF | 12.5 |
| cryoKL | 6.3 |
| cryoRV | 12.5 |
| cryoRF | 12.5 |
| cryoW1 | 25 |
| cryoW2 | 100 |
| BH cryo-tide | 50 |
| 3xFLAG | >4000 |
| cryoH | 20 |
| cryoM | 4 |
| cryoB | 4 |
| cryoCHARGE | 100 |
| cryoKF + cryoH | 10 |
| cryoKV + cryoH | 3.3 |
| CHAPSO | 42 |

**Table S3.** μ-opioid–G protein complex structures deposited in the PDB. * DAMGO bound structures. Related to Figure 2.

| PDB | Resolution (Å) | Concentration (mg/mL) |
|---|---|---|
| 8EF5<sup>44</sup> | 3.3 | 12.5 |
| 8EF6<sup>44</sup> | 3.2 | 14.8 |
| 8EFB<sup>44</sup> | 3.2 | 9.8 |
| 8EFL<sup>44</sup> | 3.2 | 8.1 |
| 8EFO<sup>44</sup> | 2.8 | 15.2 |
| 8EFQ*<sup>44</sup> | 3.3 | 3.7 |
| 8F7Q<sup>61</sup> | 3.2 | 7.5 |
| 8F7R<sup>61</sup> | 3.3 | 6.8 |
| 8K9K*<sup>62</sup> | 3 | 5 |
| 8K9L*<sup>62</sup> | 3.1 | 5 |
| 8Y72<sup>63</sup> | 2.7 | 15 |
| 8Y73<sup>63</sup> | 2.8 | 15 |
| 9PXV<sup>64</sup> | 3 | 2 |
| 9PXW<sup>64</sup> | 3.8 | 2 |
| 9PXX<sup>64</sup> | 3.1 | 2 |
| 9PXY<sup>64</sup> | 3 | 2 |
| 9PY2<sup>64</sup> | 3.2 | 2 |
| 9PY3 | 3.2 | 2 |
| 9PY4<sup>64</sup> | 2.8 | 2 |
| 9WST*<sup>65</sup> | 2.8 | 6 |
| 9WSW<sup>65</sup> | 2.8 | 12 |
| 6DDF*<sup>66</sup> | 3.5 | 7 |
| 9O36<sup>67</sup> | 2.3 | 10 |
| 9ODF<sup>68</sup> | 2.4 | 15 |
| 9ODL<sup>68</sup> | 3.6 | 15 |
| 9ODK<sup>68</sup> | 3.9 | 15 |
| 9ODJ<sup>68</sup> | 3 | 15 |
| 9ODI<sup>68</sup> | 3.7 | 15 |
| 9ODH<sup>68</sup> | 3.8 | 15 |
| 9ODE<sup>68</sup> | 2.5 | 15 |
| 6DDE*<sup>66</sup> | 3.5 | 7 |
| 7T2G<sup>69</sup> | 2.5 | 10 |
| 7T2H<sup>69</sup> | 3.2 | 20 |

**Table S4.** CryoEM grid preparation conditions. Related to Figures 1-5.

| Sample | Concentration | Peptide | Grid Type | Figure with data |
|---|---|---|---|---|
| Class A GPCR-Fab in LMNG-CHS | 1.5 mg/mL | H, M, KF, EV | ANT 1.2/1.3 Au 300 mesh | Table 1 and Figure S5F |
| μ-opioid receptor + heterotrimeric Gαi1 in LMNG-CHS | 2.5 mg/mL | EV, KV, KL, H, M | UF 1.2/1.3<br>25nm Au 300 mesh | Table 1, Figure 2, S3 |
|  | 1 mg/mL | KF | UF 1.2/1.3<br>25nm Au 300 mesh | Table 1, Figure 2, S3 |
| Class B GPCR 1 + heterotrimeric Gαs in LMNG-CHS | 1.5 mg/mL | KF | UF 1.2/1.3<br>25nm Au 300 mesh | Table 1, Figure 2, S4 |
| GPCR dimer in GDN | 0.5 mg/mL | EV, KF+H | UF 1.2/1.3<br>25nm Au 300 mesh | Table 1, S5 |
| ABCC6 in GDN | 3 mg/mL | EV | UF 1.2/1.3 Au 300 mesh | Table1, Figure 3A&B |
|  | 0.5 mg/mL | KF | UF 1.2/1.3 Au 300 mesh | Table 1, Figure 3A-F, S5A-C |
| Ion channel (nanodisc) | 1 mg/mL | KV+H | UF 1.2/1.3 Au 300 mesh | Table 1 |
| SLC transporter (nanodisc) + Fab | 3.4 mg/mL | H, EV, KV | UF 1.2/1.3<br>25nm Au 300 mesh | Table 1, Figure S5F&G |
|  | 4 mg/mL | KV | UF 1.2/1.3<br>25nm Au 300 mesh | Figure S5G |
|  | 1 mg/mL | KL, EV+H | UF 1.2/1.3<br>25nm Au 300 mesh | Figure S5F&G |
| 32 kD membrane protein in LMNG-CHS | 0.5 mg/mL | H, M, KF, KV, EV | UF 1.2/1.3<br>25nm Au 300 mesh | Table 1 |
| BCDX2-CX3- Rad51 | 0.5 mg/mL | W2, H, KV, KF | UF 1.2/1.3<br>25nm Au 300 mesh | Table 1, Figure S1C |
| Rad55-Rad57- SHU | 1 mg/mL | H, KV, KF | UF 1.2/1.3<br>25nm Au 300 mesh | Table 1 |
| Degrader complex | 1 mg/mL | EV, KV, KF, H, M, RF, KF+H, EV+H | UF 1.2/1.3<br>25nm Au 300 mesh | Table 1; Figure 1C, D; S2I–K |
| CCAN (Kinetochore) | 1 mg/mL | KF, KV, EV, M, H, KV+H | UF 1.2/1.3 Au 300 mesh | Table 1, Figure 4A&B, S6A |
| TRIP12-USP7 | 0.5 mg/mL | H, M, W1 | UF 1.2/1.3<br>25nm Au 300 mesh | Table 1, Figure 4C&D, S1C, S6B-E |
|  | 0.25 mg/mL | KF | UF 1.2/1.3<br>25nm Au 300 mesh | Table 1 |
| rFab + 20kD immune receptor | 0.2 mg/mL | KV, KV+M, KV+H | ANT 1.2/1.3 Au 300 mesh | Table 1, Figure S2B |
| rFab + 18kD cytokine | 0.25 mg/mL | EV, EV+H, KV+H | UF 1.2/1.3<br>25nm Au 300 mesh | Table 1, Figure S2A |
| OSMR + Vixa.4DS | 0.7 mg/mL | EV+H | ANT 1.2/1.3 Au 300 mesh | Figure 1F–H; S2D–E |
| Fab + immune receptor ECD (40kD) | 0.3 mg/mL | EV, KV, KF, H, M, KV+H | QF 1.2/1.3<br>Cu 300 mesh | Table 1, Figure 1A,D,E. |
| 3 Fabs + TNF family trimer | 0.25 mg/mL | H, M, KV, EV, KF | UF 1.2/1.3<br>25nm Au 300 mesh | Table 1 |
| HUWE1 | 1 mg/mL | KF, RF, H, M, LEA | ANT 1.2/1.3 Au 300 mesh | Table 1, Figure 4E-G, S7A-C |
| 70kDa tyrosine phosphatase | 0.25 mg/mL | KF | UF 1.2/1.3<br>25nm Au 300 mesh | Table 1, Figure S1E |
|  | 1 mg/mL | KF+H | UF 1.2/1.3<br>25nm Au 300 mesh | Figure S1E |
| PI3Kα + inavolisib | 1 mg/mL | M, MAX3, B, H, KF, EV, EV+H, KF+H, KV+M, KV+H, RF, KF+M, KV, RV, BH cryo-tide, LEA | ANT 1.2/1.3 Au 300 mesh | Table 1, Figure 5, S8, S9, S10A |
| “Flat” enzyme | 1.5 mg/mL | EV, KV, KF, H, M, RV, RF | ANT 1.2/1.3 Au 300 mesh | Table 1, Figure 1B&D, S1B&D, S2I |
| Class B GPCR 2<br>+ heterotrimeric Gαs in LMNG, CHS | 5.8 mg/mL |  | ANT 1.2/1.3 Au 300 mesh | Figure 2, S4 |
|  | 1.5 mg/mL | KF | ANT 1.2/1.3 Au 300 mesh | Figure 2, S4 |
|  | 0.1 mg/mL | KF | ANT 1.2/1.3 Au 300 mesh | Figure 2, S4 |
| Trimeric ligand and receptor | 0.2 mg/mL | BH cryo-tide, CHARGE, H | UF 1.2/1.3<br>25nm Au 300 mesh | Figure S1A |
| Nucleosome binding protein | 1 mg/mL | RF | UF 1.2/1.3<br>25nm Au 300 mesh | Figure S1B |
| AAV | 2.4 mg/mL | M, H, KF, EV, RF, KV, RV | ANT 1.2/1.3 Au 300 mesh | Figure S7D |
| – | – | 3xFLAG, CHARGE, H, M, KV, EV, KF | ANT 1.2/1.3 Au 300 mesh | Figure S11 |

**Table S5.** CryoEM data collection conditions. Related to Figures 1-5.

| **Sample** | **Microscope** | **Mag (kx)** | **Pixel Size (Å/pix)** | **Electron Flux (e<sup>-</sup>/pix\*sec)** | **Total Dose (e<sup>-</sup>/Å<sup>2</sup>)** | **Figure panel containing data** |
|---|---|---|---|---|---|---|
| Fab + immune receptor ECD (40kD) | Glacios2 | 150 | 0.911 | 12 | 40 | Figure 1A&D S2C&I |
| "Flat" enzyme | Krios G2 | 81 | 1.467 | 10 | 10 | Figure 1B&D, S1D, S2C&I |
| Degrader complex | Krios G2 | 165 | 0.731 | 10 | 40 | Figure 1C&D, S2C,I-K |
| Class B GPCR 1 + heterotrimeric Gαs in LMNG, CHS | Krios G3i | 130 | 0.928 | 12 | 50 | Figure 2A&B |
| Class B GPCR 2 + heterotrimeric Gαs in LMNG, CHS | Glacios | 36 | 1.114 | 15 | 48 | Figure S4C |
| Class B GPCR 1 + heterotrimeric Gαs in LMNG, CHS | Glacios | 36 | 1.114 | 15 | 48 | Figure S4A |
| Class B GPCR 1 + heterotrimeric Gαs in LMNG, CHS | Krios G3i | 105 | 1.179 | 13 | 40 | Figure S4B |
| µ-opioid receptor + heterotrimeric Gαi in LMNG, CHS | Krios G3i | 105 | 1.179 | 10 | 40 | Figure 2C-I, S3 |
| ABCC6 in GDN | Glacios2 | 150 | 0.911 | 13 | 50 | Figure 3A&B, S5A |
| ABCC6 in GDN | Krios G3i | 130 | 0.928 | 12 | 60 | Figure 3C-F, S5B&C |
| CCAN (Kinetochore) | Glacios2 | 150 | 0.911 | 12 | 41 | Figure 4A&B, S6A |
| TRIP12 | Krios |  |  |  |  | Figure S6C |
| TRIP12-USP7 | Glacios | 36 | 1.114 | 15 | 40 | Figure 4C&D, S6 E |
| TRIP12-USP7 | Krios G2 | 165 | 0.731 | 10 | 48 | Figure S6D |
| Class A GPCR-Fab in LMNG, CHS | Glacios2 | 150 | 0.911 | 12 | 40 | Figure S6F |
| HUWE1 | Krios G3i | 130 | 0.928 | 13 | 45 | Figure 4E-G, S7A-C |
| PI3Kα + inavolisib | Krios G3i | 130 | 0.928 | 12 | 45 | Figure 5, S8 |
| PI3Kα + inavolisib tilt series | Krios G3i | 64 | 1.897 | 11 | 2.6/tilt | Figure 5A, S9 |
| PI3Ka + inavolisib | Glacios | 150 | 0.931 | 15 | 50 | Figure S10A |
| Trimeric ligand and receptor | Krios G3i | 165 | 0.725 | 12 | 61 | Figure S1A |
| Protein aggregation | Glacios & Krios G2 | 36 &<br>81 | 1.114<br>1.467 | 15<br>10 | 40<br>10 | Figure S1B-E |
| Fab + immune receptor ECD (40kD) | Krios G3i | 165 | 0.725 | 12 | 51 | Figure 1E |
| rFab + 18kD cytokine | Glacios2 | 150 | 0.911 | 13 | 45 | Figure S2A |
| OSMR + Vixa.4DS | Glacios2 | 150 | 0.911 | 13 | 45 | Figure 1F-H, Figure S2D&E |
| rFab + 20kD immune receptor | Glacios2 | 150 | 0.911 | 12 | 44 | Figure S2B,F-H |
| GPCR dimer in GDN | Glacios | 36 | 1.114 | 15 | 55 | Figure S4D,E |
| ABC transporter in GDN | Krios G3i | 130 | 0.928 | 12 | 41 | Figure S5D&E |
| SLC transporter (nanodisc) + Fab | Glacios & Krios G2 | 36 &<br>105 | 1.114<br>1.179 | 15<br>11 | 40<br>40 | Figure S5F&G |
| AAV | Krios G2 | 81 | 1.467 | 10 | 20 | Figure S7D |
| Peptide self-association | Glacios | 36 | 1.114 | 15 | 40 | Figure S12B-D |
| cryoSHIELDs or additives alone | Krios G2 | 165 | 0.731 | 11 | 20 | Figure S11 |

**Table S6.** Cryo-EM data collection, refinement, and validation statistics. Related to Figures 2, 3 and 5.

|  | ABCC6 (EMD-77282) (PDB 35XZ) | OSMR-Vixa.4DS (EMD-79601) (PDB 39MH) | PI3Kα (EMD-77258, EMD-77259) |  | µ-opioid-Gi DAMGO complex (EMD-79600, EMD-77232, EMD-79365) |  |  |
|---|---|---|---|---|---|---|---|
| **Data collection and processing** | cryoKF | cryoEV/cryoH | No peptide | cryoEV / cryoH | cryoKF<br>1 mg/mL | cryoKF<br>0.5 mg/mL | No peptide<br>5 mg/mL |
| **Magnification** | 130,000x | 150,000x | 130,000x |  | 105,000x |  |  |
| **Voltage (kV)** | 300 | 200 | 300 |  | 300 |  |  |
| **Electron exposure (e–/Å2)** | 51 | 45 | 45 |  | 40 |  |  |
| **Defocus range (µm)** | -1.0 to -2.6 | -1.0 to -2.2 | -0.8 to -1.8 |  | -0.8 to -1.8 |  |  |
| **Pixel size (Å)** | 0.928 | 1.15 | 0.928 |  | 1.179 | 0.769 | 1.04 |
| **Symmetry imposed** | C1 | C1 | C1 |  | C1 |  |  |
| **Initial particle images (no.)** | 4,517,311 | 1,742,558 | 3,726,258 | 4,941,335 | 4,187,275 | 5,280,910 | 5,089,833 |
| **Final particle images (no.)** | 947,054 | 142,234 | 127,646 | 383,417 | 177,631 | 864,043 | 383,987 |
| **Map resolution (Å)**<br>**FSC threshold** | 2.49<br>0.143 | 3.4<br>0.143 | 3.5<br>0.143 | 3.2<br>0.143 | 2.6<br>0.143 | 2.3<br>0.143 | 2.6 |
| **Map resolution range (Å)** | 2.05-30 | 2.9-52.5 | 3.2-57.7 | 2.8-41.7 | 2.33-44.5 | 2.2-35.7 | 2.3-35.5 |
| **Refinement** |  |  |  |  |  |  |  |
| **Initial model used (PDB code)** | AlphaFold2<br>ab initio | AlphaFold2<br>ab initio |  |  |  |  |  |
| **Model resolution (Å)**<br>**FSC threshold** | 2.5<br>0.143 | 3.3<br>0.143 |  |  |  |  |  |
| **Model resolution range (Å)** | 2.4-3.1 | - |  |  |  |  |  |
| **Map sharpening B factor (Å2)** | 90.5 | 120.5 |  |  |  |  |  |
| **Model composition**<br>**Non-hydrogen atoms**<br>**Protein residues**<br>**Nucleotides**<br>**Ligands** | 10499<br>1340<br>0<br>PEF: 2 | 3830<br>488<br>0<br>0 |  |  |  |  |  |
| **B factors (Å2)**<br>**Protein**<br>**Nucleotide**<br>**Ligand** | 148.04<br>-<br>100.3 | 90.36<br>-<br>- |  |  |  |  |  |
| **R.m.s. deviations**<br>**Bond lengths (Å)**<br>**Bond angles (°)** | 0.004<br>0.821 | 0.009<br>0.96 |  |  |  |  |  |
| **Validation**<br>**MolProbity score**<br>**Clashscore**<br>**Poor rotamers (%)** | 1.32<br>5.83<br>0.89 | 1.03<br>1.07<br>0.23 |  |  |  |  |  |
| **Ramachandran plot**<br>**Favored (%)**<br>**Allowed (%)**<br>**Disallowed (%)** | 99.24<br>0.76<br>0.00 | 96.61<br>3.39<br>0.00 |  |  |  |  |  |

**Movie S1. Air-water interface effects mitigation by cryoSHIELDs. Related to Figure 6**.

